# A Mathematical Genomics Perspective on the Moonlighting Role of Glyceraldehyde-3-Phosphate Dehydrogenase (GAPDH)

**DOI:** 10.1101/2025.07.06.663364

**Authors:** Sk. Sarif Hassan, Debaleena Nawn, Nabanita Mukherjee, Arunava Goswami, Vladimir N. Uversky

**Author notes:** These authors contributed equally in the work (Sk. Sarif Hassan, Debaleena Nawn). Email addresses:* (Sk. Sarif Hassan), (Debaleena Nawn), (Nabanita Mukherjee), (Arunava Goswami), (Vladimir N. Uversky).

## Abstract

Glyceraldehyde-3-phosphate dehydrogenase (GAPDH) is a well-conserved enzyme across Archaea, Bacteria, and Eukarya, known not only for its canonical role in glycolysis, but also for diverse moonlighting functions including transcription regulation, host-pathogen interactions, and immune modulation. Studying GAPDH quantitatively is crucial for understanding how subtle variations at the sequence and structural levels drive such functional diversity across evolutionary lineages. In this study, 165 GAPDH protein sequences from 158 organisms were analyzed to uncover conserved and divergent features underlying multifunctionality. While core catalytic residues were strongly preserved, selective enrichment of small non-polar residues such as valine and alanine suggested a structural basis for flexibility and adaptive potential. The balanced distribution of order- and disorder-promoting residues and the avoidance of long homopolymeric stretches indicated evolutionary selection for both structural coherence and local flexibility. Spatial distribution of amino acids in GAPDH sequences revealed low fractal variance across sequences, with moderate differences in residue clustering patterns pointing to localized adaptations without compromising overall organization. These findings demonstrate that GAPDH multifunctionality is encoded through compositional signatures and conserved spatial architecture, allowing the coexistence of metabolic stability and regulatory plasticity. The results have broad implications for understanding protein evolution, structural adaptability in extreme environments, and functional versatility in pathogenic contexts. This study establishes GAPDH as a model for exploring principles of protein moonlighting and highlights the potential of quantitative compositional analysis in uncovering hidden functional layers.

## 1 Introduction

The central dogma of molecular biology once dictated that each gene encodes a single protein with a distinct, specialized function [1, 2]. Although, alternating splicing results in the production of multiple distinct mRNA transcripts — thus structurally different proteins from a single gene [3]. However, the discovery of moonlighting proteins — single polypeptides capable of performing two or more distinct and physiologically relevant functions without partitioning those roles into separate structural domains — challenges the classical view that protein structure strictly determines a single function, revealing that identical structure can support functional diversity [4, 5, 6]. These multifunctional proteins, also referred to as multitasking proteins, have now been identified across all domains of life, from bacteria to humans, revealing a widespread and conserved evolutionary strategy for expanding functional diversity without increasing genome size [7, 8, 9]. One of the best-studied and paradigmatic examples of moonlighting proteins is ‘glyceraldehyde-3-phosphate dehydrogenase’ (GAPDH). Historically known as a key glycolytic enzyme, catalyzing the conversion of glyceraldehyde-3-phosphate to 1, 3-bisphosphoglycerate, GAPDH is now widely recognized as a quintessential moonlighting protein with roles far beyond energy metabolism [10, 11, 12, 13]. GAPDH’s functional versatility spans multiple cellular compartments, including the cytoplasm, nucleus, membrane, and extracellular matrix, reflecting a remarkable degree of spatial and functional plasticity [14, 15]. In these diverse locales, GAPDH is involved in apoptosis regulation, membrane fusion, endocytosis, iron homeostasis, cytoskeletal dynamics, tubulin regulation, transcriptional control, mRNA stability, DNA repair, immune evasion, and host-pathogen interactions, among others [6]. These moonlighting activities are often modulated by post-translational modifications such as phosphorylation, nitrosylation, acetylation, and O-GlcNAcylation, as well as by conformational changes, binding partners, and subcellular localization [16, 17].

GAPDH moonlighting is not limited to eukaryotic cells [18, 19]. In bacterial and fungal pathogens, GAPDH has been co-opted as a virulence factor, functioning as an adhesin, a receptor for host proteins like transferrin and plasminogen, and a mediator of immune modulation [20, 4]. Its presence on the surface of pathogens, despite lacking classical secretion signals, facilitates host tissue colonization, immune evasion, and systemic dissemination [21, 22]. In probiotic bacteria such as *Lactobacillus gasseri*, moonlighting GAPDH has been shown to alleviate allergic asthma by modulating dendritic cell signaling [23, 24]. Similarly, in parasitic organisms like *Fasciola gigantica*, GAPDH supports energy metabolism and exhibits structural adaptations that may affect host-parasite interactions [25, 26].

The evolutionary conservation and diversification of GAPDH across kingdoms underscores its fundamental importance and functional adaptability [27, 15]. Despite strong conservation in its glycolytic catalytic core, sequence variations — especially in non-catalytic regions — may underpin the emergence of novel moonlighting functions [10, 28, 29, 30]. These variations are likely to reflect species-specific pressures, such as environmental stress, immune challenge, or metabolic demands, contributing to organismal fitness and pathogenicity [31, 32].

In this context, a comprehensive comparative analysis of GAPDH protein sequences from diverse organisms not only facilitates the identification of evolutionary conserved motifs and structural domains, but also allows us to pinpoint sequence heterogeneities that may correspond to moonlighting adaptations [33, 30]. Such quantitative insights are crucial to unravel the structure–function relationship of GAPDH, to understand the molecular logic of functional diversification, and to trace the evolutionary history of its secondary roles.

Moreover, given GAPDH’s implications in cancer, neurodegeneration, autoimmune disorders, and infectious disease, a deeper understanding of its multifunctionality at the sequence level could inform novel therapeutic strategies, such as selectively targeting disease-related moonlighting functions without disrupting essential metabolic activity [19]. This systems-level understanding demands quantitative approaches rooted in comparative genomics, structural bioinformatics, and functional annotation [33, 30].

This study, therefore, undertakes a quantitative genomic analysis of 165 GAPDH sequences from 158 organisms, aiming to dissect sequence conservation, variation, and potential functional divergence. By integrating evolutionary, structural, and functional perspectives, we offer a novel lens to understand the moonlighting nature of GAPDH along with its biological significance across taxa. This approach promises to contribute meaningfully to the broader field of moonlighting protein biology, especially in relation to functional genomics and molecular evolution.

## 2 Data

In the present study, 165 Glyceraldehyde-3-phosphate dehydrogenase (GAPDH) protein sequences from 158 distinct organisms were extracted from the UniPort database (Table 2). Note that all these sequences were UniProtKB reviewed (Swiss-Prot).

**Table 1.**
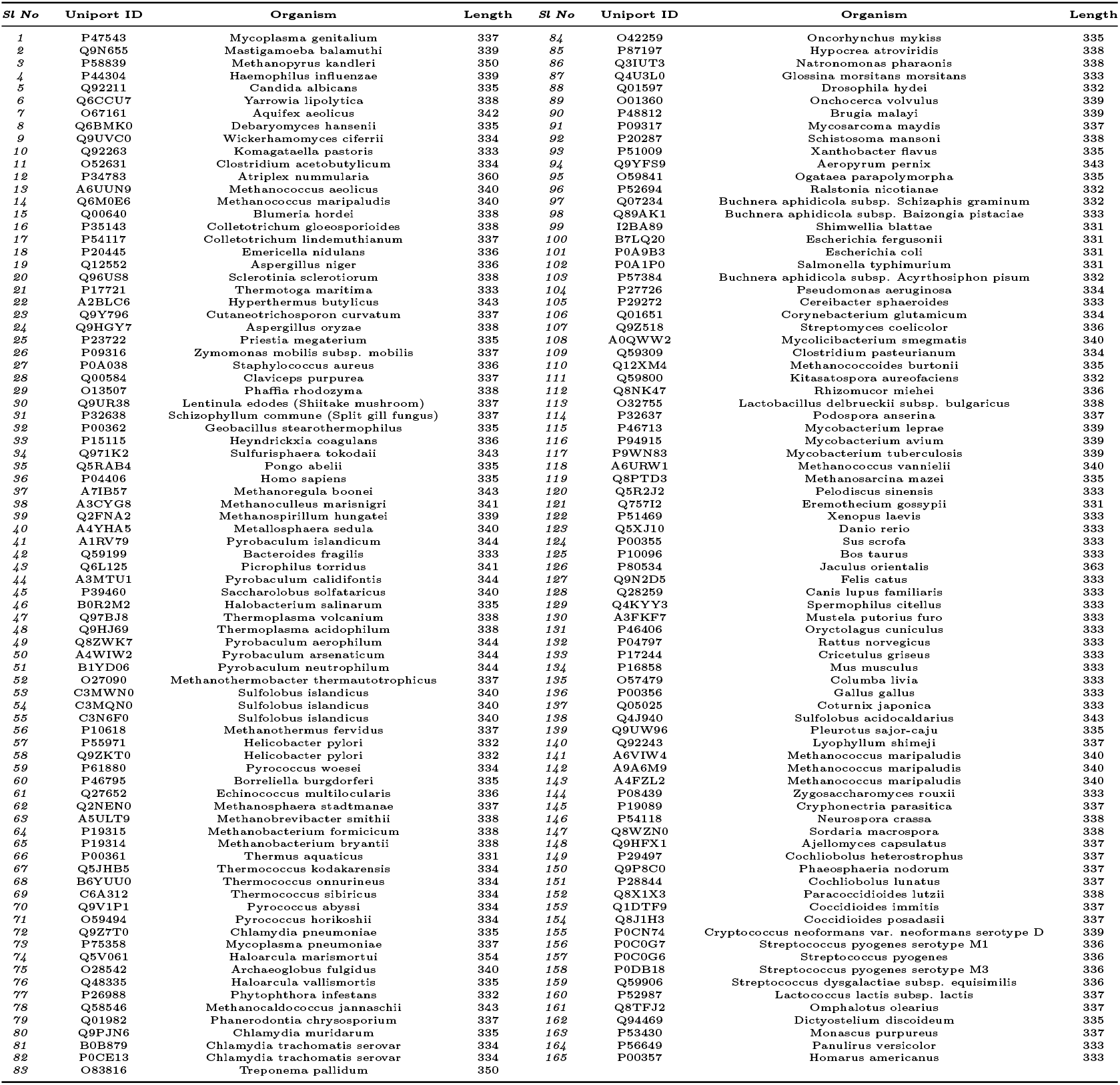
List of 165 GAPDH moonlighting proteins from 158 different organisms with their associated Uniprot ID (Hyperlinked with respective Uniprot webpage)

**Table 2.** List of sequence variations at various positions based on aligned sequence.

| Variations | Transitions |  |  |  |  |
| --- | --- | --- | --- | --- | --- |
|  | From AA | To AA | From Type | To Type | Change Type |
| **I12V** | Isoleucine | Valine | Non-polar | Non-polar | Conservative |
| **F15Y** | Phenylalanine | Tyrosine | Non-polar | Polar | **Non-polar → Polar** |
| **I18V** | Isoleucine | Valine | Non-polar | Non-polar | Conservative |
| **R20K** | Arginine | Lysine | Polar | Polar | Conservative |
| **V40I** | Valine | Isoleucine | Non-polar | Non-polar | Conservative |
| **D45E** | Aspartic acid | Glutamic acid | Polar | Polar | Conservative |
| **D45K** | Aspartic acid | Lysine | Polar | Polar | Conservative |
| **L111F** | Leucine | Phenylalanine | Non-polar | Non-polar | Conservative |
| **L111I** | Leucine | Isoleucine | Non-polar | Non-polar | Conservative |
| **L111V** | Leucine | Valine | Non-polar | Non-polar | Conservative |
| **L111M** | Leucine | Methionine | Non-polar | Non-polar | Conservative |
| **E148D** | Glutamic acid | Aspartic acid | Polar | Polar | Conservative |
| **N199T** | Asparagine | Threonine | Polar | Polar | Conservative |
| **N199S** | Asparagine | Serine | Polar | Polar | Conservative |
| **C216S** | Cysteine | Serine | Polar | Polar | Conservative |
| **T217N** | Threonine | Asparagine | Polar | Polar | Conservative |
| **T218N** | Threonine | Asparagine | Polar | Polar | Conservative |
| **T218I** | Threonine | Isoleucine | Polar | Non-polar | **Polar → Non-polar** |
| **L221I** | Leucine | Isoleucine | Non-polar | Non-polar | Conservative |
| **D225E** | Aspartic acid | Glutamic acid | Polar | Polar | Conservative |
| **D225K** | Aspartic acid | Lysine | Polar | Polar | Conservative |
| **L228I** | Leucine | Isoleucine | Non-polar | Non-polar | Conservative |
| **L228V** | Leucine | Valine | Non-polar | Non-polar | Conservative |
| **L228F** | Leucine | Phenylalanine | Non-polar | Non-polar | Conservative |
| **L228M** | Leucine | Methionine | Non-polar | Non-polar | Conservative |
| **H244R** | Histidine | Arginine | Polar | Polar | Conservative |
| **T282S** | Threonine | Serine | Polar | Polar | Conservative |
| **T282N** | Threonine | Asparagine | Polar | Polar | Conservative |
| **I292L** | Isoleucine | Leucine | Non-polar | Non-polar | Conservative |
| **I292M** | Isoleucine | Methionine | Non-polar | Non-polar | Conservative |
| **I292F** | Isoleucine | Phenylalanine | Non-polar | Non-polar | Conservative |
| **L299M** | Leucine | Methionine | Non-polar | Non-polar | Conservative |
| **L299I** | Leucine | Isoleucine | Non-polar | Non-polar | Conservative |
| **L299V** | Leucine | Valine | Non-polar | Non-polar | Conservative |
| **A303S** | Alanine | Serine | Non-polar | Polar | **Non-polar → Polar** |
| **A303G** | Alanine | Glycine | Non-polar | Non-polar | Conservative |
| **A303S** | Alanine | Serine | Non-polar | Polar | **Non-polar → Polar** |
| **I330L** | Isoleucine | Leucine | Non-polar | Non-polar | Conservative |
| **I330V** | Isoleucine | Valine | Non-polar | Non-polar | Conservative |
| **K393E** | Lysine | Glutamic acid | Polar | Polar | Conservative |
| **K393R** | Lysine | Arginine | Polar | Polar | Conservative |
| **W397F** | Tryptophan | Phenylalanine | Non-polar | Non-polar | Conservative |

## 3 Methods

### 3.1 Amino acid sequence homology and phylogeny

Amino acid sequence homology among 165 GAPDH sequences is analyzed using multiple sequence alignment performed via Clustal Omega to identify conserved residues and sequence variations [34]. Furthermore, type of change based on hydrophobicity was determined for each sequence variations.

#### 3.1.1 Amino acid frequency-based Shannon variability for each residue of GAPDH sequences

Shannon entropy is a fundamental measure for assessing the diversity of amino acid residues at individual positions within the multiple sequence alignment of GAPDH sequences [35]. The entropy measure of variability at each alignment position is quantified using Shannon variability (*H*_*v*_), calculated as:

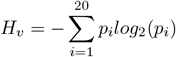

where *p*_*i*_ denotes the proportion of residues corresponding to amino acid type *i* at a given position. The Shannon variability score (*H*_*v*_) ranges from 0 (indicating complete conservation with only one residue present) to a maximum of 4.322 (reflecting equal representation of all 20 amino acids) [36]. Positions with *H*_*v*_ *>* 2.0 are classified as *variable*, those with *H*_*v*_ *<* 2.0 as *conserved*, and positions with *H*_*v*_ *<* 1.0 are specifically identified as *highly conserved* [36, 35].

### 3.2 Determining amino acid frequency composition in GAPDH protein sequences

The amino acid frequency—defined as the number of occurrences of each amino acid within a protein sequence—was computed for all 165 GAPDH moonlighting proteins [37, 38, 39]. To normalize for differences in sequence length, the relative frequency of each amino acid was calculated by dividing its absolute frequency by the total length of the corresponding protein sequence and multiplying by 100. This relative frequency reflects the percentage composition of each amino acid within the sequence. Accordingly, each protein sequence is represented as a 20-dimensional vector, comprising the relative frequencies of the 20 standard amino acids.

### 3.3 Composition profiler of GAPDH protein sequences

Composition Profiler represents a useful web-based tool for visualization of these amino acid composition biases providing means for the semi-automatic discovery of enrichment or depletion of amino acids in query proteins [40]. The Composition Profiler is used to generate an amino acid composition profile of all the GAPDH proteins analyzed in this study [40]. This set of amino acid sequences is the query set and the ‘Protein Data Bank Select 25’ is the background set. We also generated a composition profile for experimentally validated disordered proteins from the DisProt [41, 42, 43]. The generated profiles represent plots showing normalized enrichment or depletion of a given residue calculated as 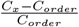, where *C*_*x*_ is the content of a given residue in its query protein, and *C*_*order*_ is the content of the same residue in the PDB Select 25.

### 3.4 Determining homogeneous poly-string frequency of amino acids in GAPDH protein sequences

A homogeneous poly-string of length *n* is defined as a subsequence consisting of n consecutive repetitions of the same amino acid [44, 45]. For instance, in the sequence “PPPWWPWW”, there is one homogeneous poly-string of ‘P’ with length 3, another of ‘P’ with length 1, and two homogeneous poly-strings of ‘W’, each of length 2. When identifying homogeneous poly-strings of a specific length *n*, only those with exactly length n are counted—longer strings are not subdivided. To systematically analyze these poly-strings across all GAPDH protein sequences, we first determined the maximum length of homogeneous poly-strings observed for any amino acid in the dataset. Using this maximum length as a reference, we then counted the number of homogeneous poly-strings for each amino acid at each possible length from 1 up to the maximum in every protein sequence [45].

### 3.5. Evaluating polar, nonpolar residue profiles of GAPDH sequences

Each amino acid in a given GAPDH moonlighting protein sequence is categorized as either polar (P) or nonpolar (N). Thus, the original protein sequence is transformed into a binary sequence composed of the symbols P and N, representing the spatial distribution of polar and non-polar residues in the amino acid sequence [46].

#### 3.5.1. Determining homogeneous poly-string frequency of polar, nonpolar residues

The frequencies of homogeneous poly-strings composed of polar and nonpolar residues are determined across all 165 GAPDH sequences, following the same approach used for computing homogeneous poly-string frequencies of individual amino acids. For an instance, consider an amino acid sequence ‘DEAFAHSVW’ which transformed into into a binary polar-nonpolar profile–’PPNNNPPNN’. It consists of two homogeneous poly-strings of ‘P’ of length 2, two homogeneous poly-strings of ‘N’ of length 2, and one homogeneous poly-string of ‘N’ of length 3 [46].

#### 3.5.2. Frequency of changes between adjacent residues from polar-nonpolar profiles of GAPDH sequences

In the binary polar, nonpolar profile of each GAPDH moonlighting protein sequence, four possible changes can occur between two adjacent residues: polar to polar (PP), polar to nonpolar (PN), nonpolar to nonpolar (NN), and nonpolar to polar (NP). The frequency of each of four changes (PP, PN, NN, and NP) is calculated for each GAPDH protein from its polar-nonpolar profile and the frequency further be divided by length of the respective GAPDH protein sequence [45].

#### 3.5.2. Walk based on polar, non-polar profiles of GAPDH sequences

A protein walk based on polar-nonpolar profile, termed as ‘P-NP walk’, is defined as:

In cartesian coordinate system, starting from the origin (0, 0), for the first residue in polar-nonpolar profile, the walker moves one unit along positive x-axis, if the residue is polar and else it moves one unit along positive y-axis and, walker moves forward accordingly till last residue of the polar-nonpolar profile. For all GAPDH sequences, P-NP walks are generated [47].

### 3.6. Evaluating intrinsic protein disorder in GAPDH sequences

The intrinsic disorder propensity of all GAPDH protein sequences are evaluated using a suite of well-established per-residue disorder predictor, the Rapid Intrinsic Disorder Analysis Online (RIDAO) platform. Among all, VSL2B predictor result are considered for further analyses [48, 49, 50].

Disorder scores of each residue range from 0 (completely ordered) to 1 (completely disordered). Residues with scores above 0.5 are classified as disordered (‘D’). Those with scores between 0.25 and 0.5 are classified as highly flexible (‘HF’), while scores from 0.1 to 0.25 are categorized as moderately flexible (‘MF’). Residues with scores below 0.1 are grouped under the ‘other’ (‘O’) category [48, 51].

#### 3.6.1. Percentage of four intrinsic protein disorder residue types in each amino acid

As the distributions of amino acids are non-uniform (relative frequency of some amino acids are very small, while some possess very large frequency), counts of each of the four residue types (‘D’, ‘HF’, ‘MF’, and ‘O’) for a particular amino acid are divided by the corresponding amino acid frequency and multiplied by 100 in each sequence. It is to be noted that percentages are calculated based on individual amino acid frequency and not with respect to the total number of residues of a given type in a sequence. Hence the sum of the percentages of different amino acids will not be 100 for a given residue type in a sequence. Even if the frequency of an amino acid is much lower in any sequence, its distribution among four types of residues can be better reflected in this way [51].

#### 3.6.2. Frequency of changes between adjacent residues of intrinsic protein disorder profiles of GAPDH sequences

There are sixteen possible changes between two adjacent residues in a sequence, based on their classification as disordered (D), highly flexible (HF), moderately flexible (MF), or other (O). These transitions include: D_D, D_HF, D_MF, D_O, HF_D, HF_HF HF_MF, HF_O, MF_D, MF_HF, MF_MF, MF_O, O_D, O_HF, O_MF, and O_O. The frequency of each of the sixteen transitions is quantified for each protein, offering insights into the dynamics of flexibility and disorder within the protein sequences [30].

#### 3.6.3. Walk based on intrinsic protein disorder profiles of GAPDH sequences

A protein walk based on intrinsic protein disorder profiles henceforth termed as ‘IPD Walk’ is defined as: In the cartesian coordinate system, starting from the origin (0, 0), for the first residue in intrinsic protein disorder profiles, the walker moves five (for better visualization) units along positive x-axis, if the residue is disordered. If the residue is highly flexible, walker moves five units at an angle 30 degree with positive x-axis. If the residue is moderately flexible, walker moves five units at an angle 60 degree with positive x-axis. If the residue type is ‘other’, walker moves five units along positive y-axis. Walker moves accordingly till last residue of the intrinsic protein disorder profiles. Note that, angles are chosen in such a way that no loop is formed along the walk. For all GAPDH sequences, IPD walks are generated.

### 3.7. Indicator matrix generation and its derivatives

For a given GAPDH amino acid sequence, The indicator function is defined as:

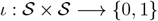

such that

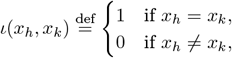

Here *x*_*h*_, *x*_*k*_ ∈ 𝒮 and 𝒮 is the set of all twenty amino acids. Clearly, *ι*(*x*_*h*_, *x*_*k*_) = *ι*(*x*_*k*_, *x*_*h*_) and *ι*(*x*_*h*_, *x*_*h*_) = 1. Hence, According to the indicator function definition, the indicator of an *l* length amino acid sequence can be easily represented by *l × l* sparse symmetric matrix of binary values of 0 and 1, which results from the indicator matrix ℐ _*hk*_ = *ι*(*x*_*h*_, *x*_*k*_), and *h, k ∈ {*1, 2, 3, …, *l}*. The resulting square matrix is a binary image, where white pixels represent ℐ _*hk*_ = 1, and black represents ℐ _*hk*_ = 0 [52].

Fractal dimension, density of ones and frequency of clusters with 4 and 8 connectivity are enumerated from the indicator matrix [52, 53]. Note that, frequency of clusters with 4 and 8 connectivity of a GAPDH sequence are normalized by multiplying with the ratio of the highest length among all 165 GAPDH sequences and length of the respective GAPDH sequence.

### 3.8. Formation of distance matrices and dendrograms

MSA based distance matrix: Similarity matrix is obtained from pairwise GAPDH protein sequences identity percentages and the corresponding distance matrix is obtained by subtracting similarity matrix entries from 100.

Euclidean distances are computed between the feature-vectors of all pairs of GAPDH protein sequences for each of the following three features:

- Relative frequency of amino acids (dimension 20)
- Relative frequency of changes from polar-nonpolar profiles (dimension 4)
- Relative frequency of changes from intrinsic protein disorder profiles (dimension 16)

Furthermore, Hausdorff distance is calculated between all pairs of GAPDH protein sequences for:

- P-NP Walks based on polar-nonpolar residue profiles
- IPD Walks based on intrinsic protein disorder profiles

As the GAPDH sequences are of varying lengths, P-NP and IPD walks are scaled. Prior to calculation of Hausdorff distance between a pair of GAPDH sequences *Gs*_1_ and *Gs*_2_ having lengths *l*_1_ and *l*_2_ such that (*l*_1_ *< l*_2_), walk of *Gs*_2_ is scaled down by the scaling factor 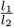.

A clustering threshold (empirically chosen) is applied to a dendrogram to define distinct clusters of GAPDH proteins, and a smaller secondary threshold (typically 20% of the primary threshold) is used to identify subsets containing proximal GAPDH sequences (these subsets are called as proximal sets) within these clusters.

### 3.9. Proximal relationships of GAPDH proteins

To identify the final proximal GAPDH proteins, a maximal intersecting family of proximal sets are obtained by those GAPDH sequences which are turned out to be proximal with regards to all six dendrograms with respect to the secondary thresholds.

## 4 Results and analyses

### 4.1. Sequence-based phylogenetic relationships among GAPDH moonlighting protein sequences

### 4.1.1 Sequence variations and invariance

Multiple sequence alignment (MSA) of 165 GAPDH protein sequences revealed that four invariant residues viz. G14G, G16G, G19G, and S215S were identified. In Table 2, list of sequence variations as mentioned in section 3.1 were presented. Note that, here positions were noted based on aligned sequences, so no reference sequence was needed. Only three sequence variations were noted F15Y, T218I, and A303S, which were typically non-polar to polar, whereas only one change T218I found to be of polar to non-polar type. Remaining transitions were all conservative as listed in Table 2.

An examination of the multiple sequence alignment (MSA) of 165 GAPDH protein sequences revealed that 27.25% of the residues were highly conserved, 38.16% were moderately conserved, and the remaining 34.59% exhibited variability across the sequences (Figure 1).

**Figure 1.**
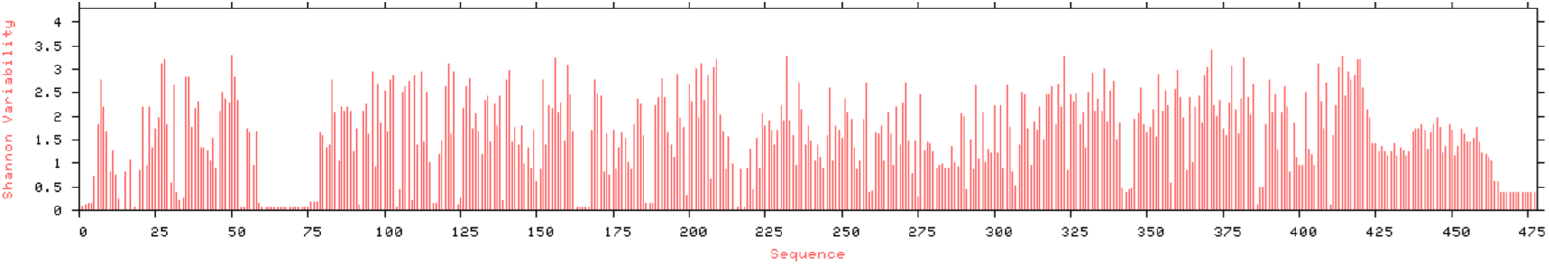
Shannon entropy of GAPDH sequence variability

#### 4.1.2 Phylogenetic relationships among GAPDH sequences based on amino acid sequence homolog*y*

Thirteen clusters were emerged based on sequence homology as derived in the phylogeny (Figure 2) by setting a distance threshold of 45. The largest cluster consists of 86 GAPDH sequences, represented in light green in the dendrogram (Figure 4). When the threshold was set to 10, a total of 65 moonlighting sequences were grouped into 20 disjoint sets, as noticed in the dendrogram (Figure 2) and detailed in Table 3.

**Table 3.** List of sets consist of proximal GAPDH protein sequences based on amino acid sequence homology.

| Sets based on the proximity derived from amino acid sequence homology |  |  |  |
| --- | --- | --- | --- |
| **Set-1** | {53, 54, 55} | **Set-11** | {87, 88} |
| **Set-2** | {59, 70, 71} | **Set-12** | {164, 165} |
| **Set-3** | {67, 68} | **Set-13** | {89, 90} |
| **Set-4** | {14, 141, 142, 143} | **Set-14** | {16, 17} |
| **Set-5** | {64, 65} | **Set-15** | {146, 147} |
| **Set-6** | {11, 109} | **Set-16** | {149, 151, 150} |
| **Set-7** | {156, 157, 158, 159} | **Set-17** | {18, 19} |
| **Set-8** | {116, 117} | **Set-18** | {153, 154} |
| **Set-9** | {99, 100, 101, 102} | **Set-19** | {80, 81, 82} |
| **Set-10** | {35, 36, 124, 131, 125, 127, 128, 129, 130, 132, 134, 133, 126, 135, 136, 137, 120} | **Set-20** | {57, 58} |

**Figure 2.**
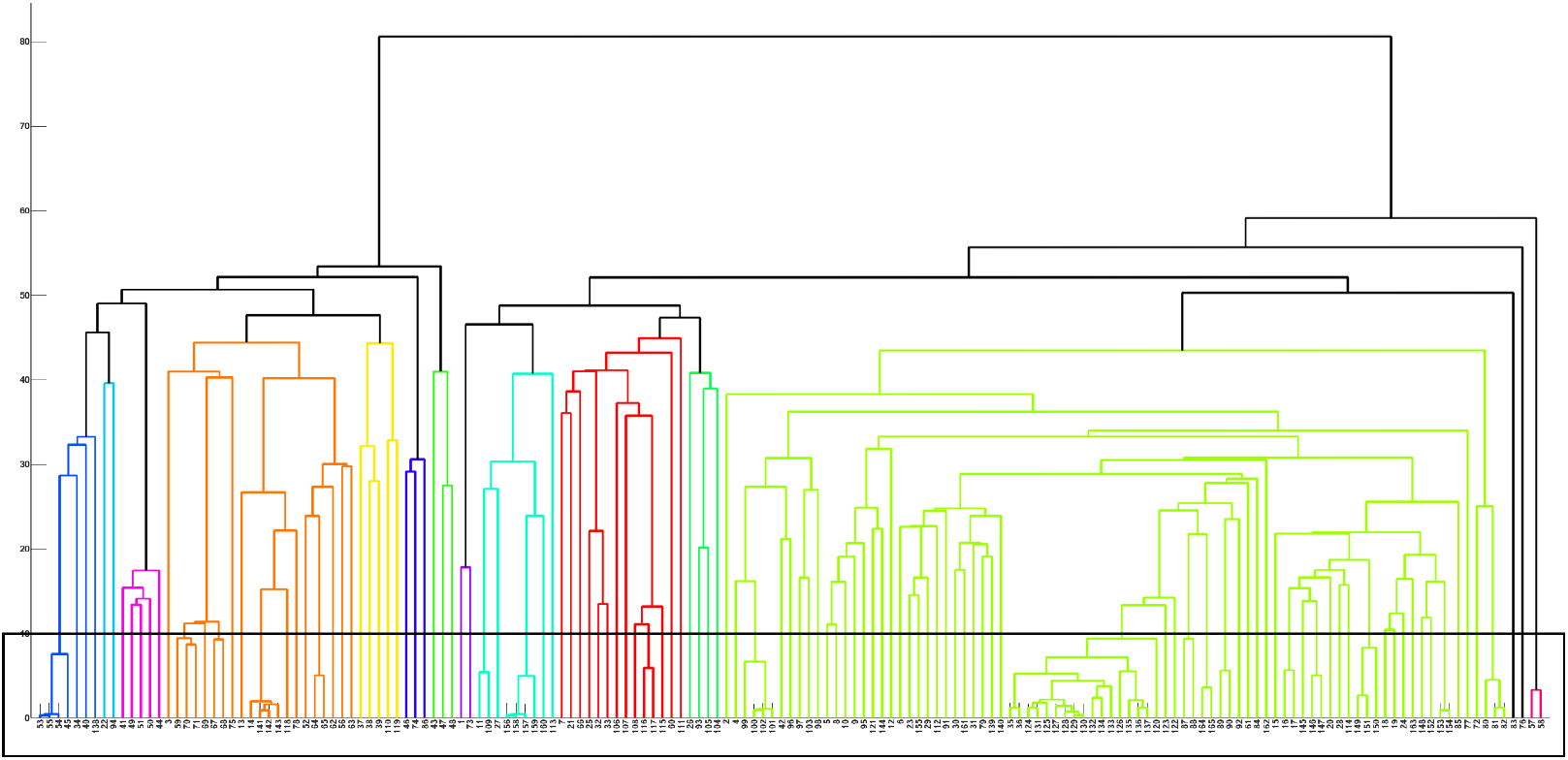
Phylogenetic relationships among GAPDH sequences based on sequence homology

Among these, all the members of the Sets-1, 2, 3, 4, and 5 fell under the domain Archaea, while the GAPDH sequences of the sets 6, 7, 8 9, 19 and 20 belonged to the domain Bacteria. The GAPDH proteins of the remaining nine sets (Sets-10, 11, 12, … 18) were grouped under the domain Eukaryota, comprising representatives from the kingdoms Fungi and Animalia. Specifically,

- Set-10 includes organisms from Animalia, with members classified under phylum Chordata, specifically from the classes Mammalia, Reptilia, and Aves.
- Members of the Set-11 and Set-12 were from Animalia, phylum Arthropoda, with classes Insecta and Malacostraca, respectively.
- Organisms of Set-13 also belonged to Animalia, but from the phylum Nematoda, class Chromadorea.
- Set-14 to Set-18 consisted entirely of organisms from the kingdom Fungi.

This domain-wise and taxonomic distribution highlights the evolutionary span and functional diversity of GAPDH moonlighting proteins across all three domains of life.

### 4.2 Relative frequency of amino acids in GAPDH moonlighting protein sequences

Relative frequencies of amino acids across 165 moonlighting protein sequences were enumerated. Due to the skewed distribution patterns observed across most amino acids in 165 GAPDH moonlighting protein sequences, median values were preferred over means for central tendency (Figure 3).

**Figure 3.**
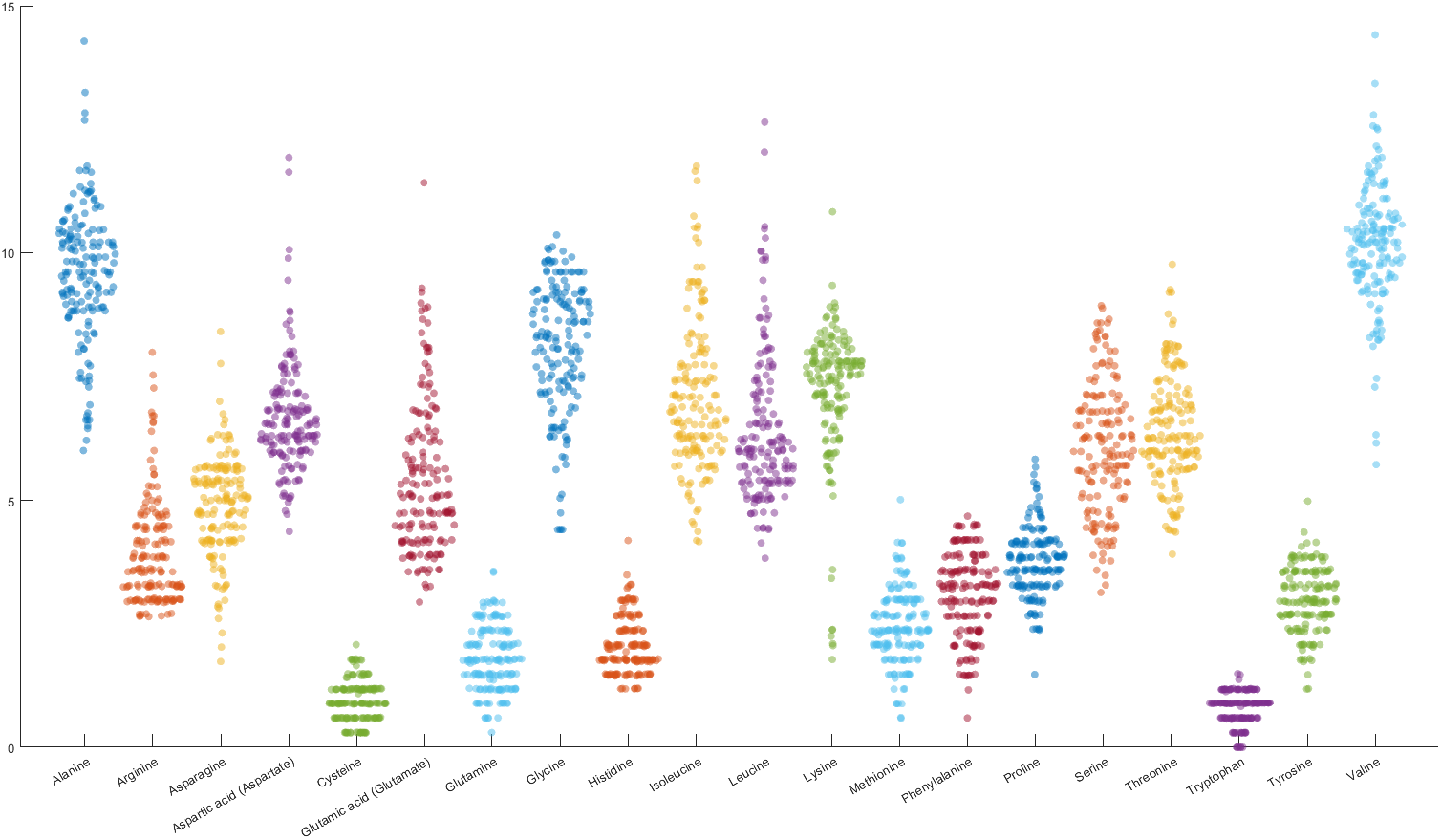
Histogram of relative frequency of each amino acid present in GAPDH protein sequences.

Among all amino acids, valine emerged as the most predominant in terms of relative abundance, displaying the highest median value (10.210) across the 165 moonlighting protein sequences analyzed. Alanine was the second most abundant, with a median value of 9.610. The maximum percentage of valine was recorded in the protein sequence with UniProt ID A4YHA5 (14.412%), while the highest percentage of alanine was observed in the sequence P15115 (14.286%). These results indicate a consistent enrichment of valine and alanine across moonlighting proteins from diverse organisms.

However, only a few sequences showed other amino acids as the most abundant. These included:

- Leucine was most abundant in the sequence with UniProt ID Q9ZKT0 (12.651%), where the percentages of valine and alanine were approximately half that of leucine.
- Isoleucine showed its highest percentage in A6URW1 (11.765%), while valine and alanine were present at 8.824% and 8.235%, respectively.
- Lysine reached its maximum proportion in Q07234 (10.843%), with comparable percentages of isoleucine (9.036%), leucine (8.735%), and valine (8.434%).
- Glycine was most abundant in Q6CCU7 (10.355%), where the percentage of valine was equal, and the alanine content was also very close.

In contrast to the predominant amino acids, tryptophan and cysteine were the least frequent across the majority of the sequences, both exhibiting the lowest median value of 0.896. Remarkably, tryptophan was completely absent in several sequences, including Q971K2, A4YHA5, P39460, C3MWN0, C3MQN0, C3N6F0, and Q4J940. In 163 out of the 165 sequences analyzed, either tryptophan or cysteine represented the minimum frequency amino acid. For the remaining two sequences—Q01982 and Q94469—glutamine was found to be the least abundant, with a median value of 1.780.

In terms of variation, glutamic acid exhibited the highest median standard deviation (1.540), followed by leucine (median standard deviation 1.537) indicating considerable variability in its representation across sequences.

Based on a distance threshold of 5, the largest cluster consisted of 71 sequences, represented in green in the dendrogram (Figure 4). Based on a distance threshold of 1, 30 GAPDH moonlighting protein sequences were organized into 11 distinct sets (Figure 4, Table 4). Taxonomic profiling revealed that members of the Set-1 and Set-2 belong to the domain Archaea, while members belonging to the Set-3, Set-4, and Set-5 were of Bacterial origin. The remaining six sets (Set-6, 7, … 11) fell under the domain Eukaryota. Among these members of the Set-6 and Set-7 were fungal, whereas Set-8, Set-9, and Set-10 represent mammalian GAPDH sequences from the phylum Chordata, and members of the Set-11 correspond to Aves. This classification underscores the evolutionary breadth of GAPDH moonlighting proteins and their conserved presence across all domains of life.

**Figure 4.**
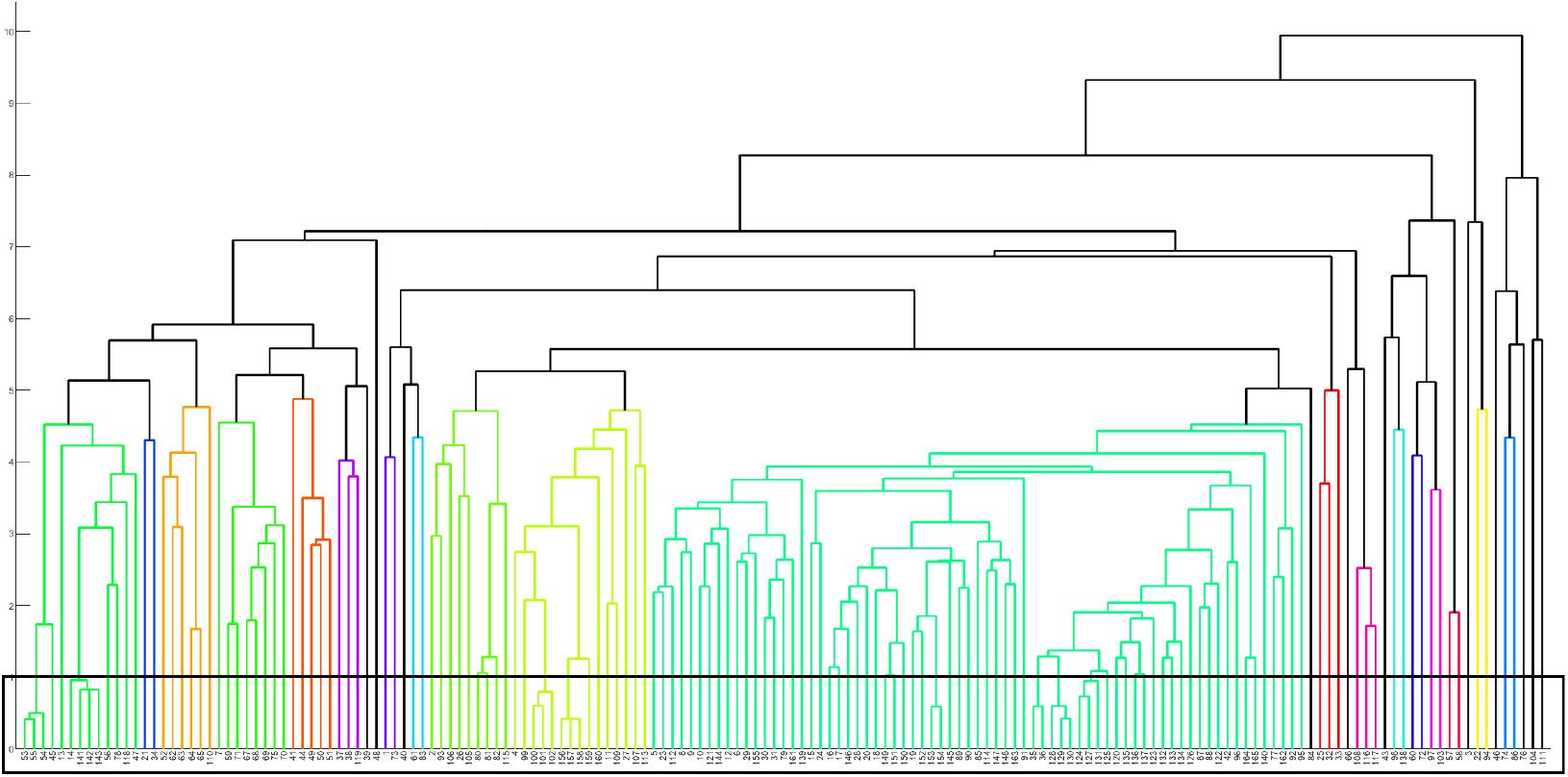
Phylogenetic relationship among the GAPDH moonlighting proteins based on relative frequency of amino acids.

**Table 4.**
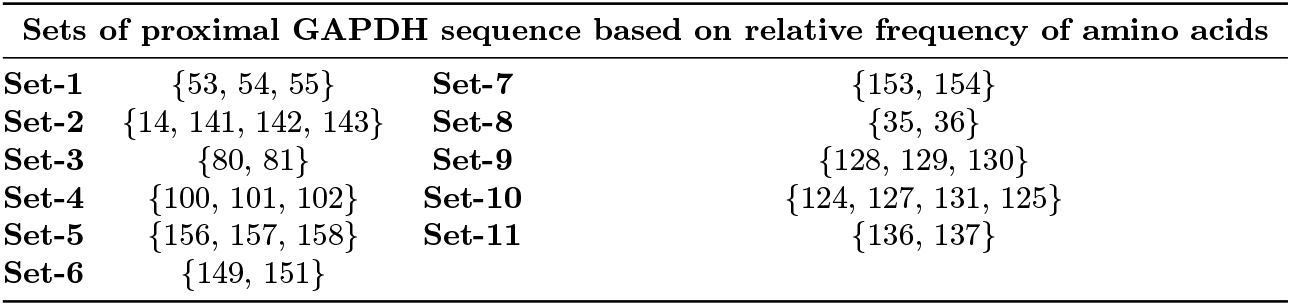
List of sets consist of proximal GAPDH protein sequences based on relative frequency of amino acids.

### 4.3 Composition profiler of GAPDH protein sequences

Figure 5 represents the composition profile generated for a set of 165 GAPDH proteins using Composition Profiler [40]. Logistics behind this analysis is rooted in the existence of noticeable differences between the amino acid compositions of amino acid sequences coding for ordered proteins/domains and intrinsically disordered proteins/regions [54, 55, 56, 57, 40, 58]. In fact, in comparison with ordered proteins, disordered proteins/regions are known to be significantly depleted in so-called order-promoting amino acids, such as C, W, I, Y, F, L, H, V, N, and M, being instead noticeably enriched in disorder-promoting amino acids, R, T, D, G, A, K, Q, S, E, and P [54, 55, 56, 57, 40, 58].

**Figure 5.**
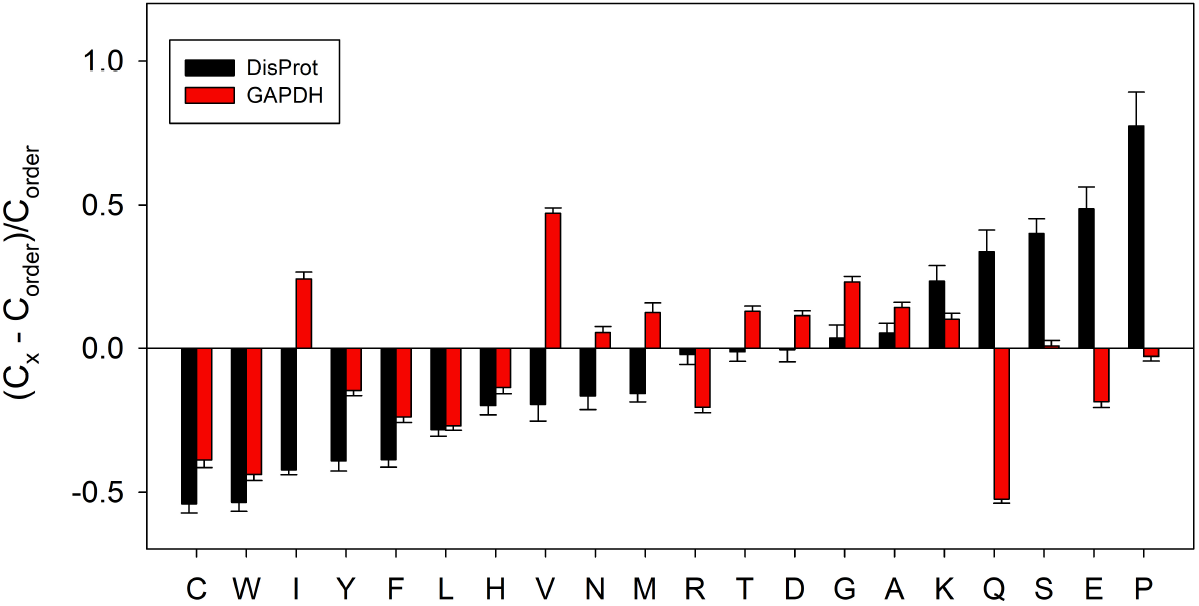
Amino acid composition profile of 165 GAPDH proteins (red bars). The fractional difference is calculated as 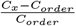, where *C*_*x*_ is the content of a given amino acid in the query set (GAPDH proteins or known intrinsically disordered proteins) and *C*_*order*_ is the content of a given amino acid in the background set (Protein Data Bank Select 25). The amino acid residues are ranked from most order-promoting residue to most disorder-promoting residue. Positive values indicate enrichment and negative values indicate depletion of a particular amino acid. The composition profile generated for experimentally validated disordered proteins from the DisProt database (black bars) is shown for comparison. In both cases, error bars correspond to standard deviations over 10,000 bootstrap iterations.

Figure 5 shows that GAPDH proteins analyzed in this study are depleted in several order-promoting residues, including C, W, Y, F, L, and H. These proteins are also depleted in three disorder-promoting residues, R, Q, and E. On the other hand, GAPDH proteins are enriched in four order promoting residues (I, V, N, and M) and five disorder-promoting residues (T, S, G, A, and K).

### 4.4. Homogeneous poly-string frequency of amino acids in GAPDH moonlighting protein sequences

The longest homogeneous poly-string observed across all amino acids in the 165 GAPDH moonlighting sequences is of length 4. Accordingly, the frequencies of homogeneous poly-strings of lengths 1 to 4 for each of the twenty amino acids in every sequence were recorded (**Supplementary File-1**). The total frequency of poly-strings, aggregated across all amino acids for each sequence, are summarized in Table 5.

**Table 5.** Frequency of homogeneous poly-string of amino acids of length 1, 2, 3, and 4 of *GAPDH* moonlighting proteins.

| *Sl No* | Uniport ID | l=1 | l=2 | l-3 | l-4 | *Sl No* | Uniport ID | l=1 | l=2 | l-3 | l-4 | *Sl No* | Uniport ID | l=1 | l=2 | l-3 | l-4 |
| --- | --- | --- | --- | --- | --- | --- | --- | --- | --- | --- | --- | --- | --- | --- | --- | --- | --- |
| 1 | P47543 | 285 | 23 | 2 | 0 | 56 | P10618 | 297 | 20 | 0 | 0 | 111 | Q59800 | 285 | 19 | 3 | 0 |
| 2 | Q9N655 | 285 | 24 | 2 | 0 | 57 | P55971 | 293 | 15 | 3 | 0 | 112 | Q8NK47 | 290 | 23 | 0 | 0 |
| 3 | P58839 | 295 | 26 | 1 | 0 | 58 | Q9ZKT0 | 293 | 15 | 3 | 0 | 113 | O32755 | 299 | 18 | 1 | 0 |
| 4 | P44304 | 305 | 17 | 0 | 0 | 59 | P61880 | 294 | 20 | 0 | 0 | 114 | P32637 | 300 | 17 | 1 | 0 |
| 5 | Q92211 | 281 | 27 | 0 | 0 | 60 | P46795 | 295 | 17 | 2 | 0 | 115 | P46713 | 292 | 19 | 3 | 0 |
| 6 | Q6CCU7 | 290 | 24 | 0 | 0 | 61 | Q27652 | 292 | 22 | 0 | 0 | 116 | P94915 | 284 | 23 | 3 | 0 |
| 7 | O67161 | 291 | 21 | 3 | 0 | 62 | Q2NEN0 | 293 | 22 | 0 | 0 | 117 | P9WN83 | 282 | 24 | 3 | 0 |
| 8 | Q6BMK0 | 284 | 24 | 1 | 0 | 63 | A5ULT9 | 289 | 23 | 1 | 0 | 118 | A6URW1 | 288 | 26 | 0 | 0 |
| 9 | Q9UVC0 | 284 | 25 | 0 | 0 | 64 | P19315 | 295 | 20 | 1 | 0 | 119 | Q8PTD3 | 291 | 22 | 0 | 0 |
| 10 | Q92263 | 285 | 24 | 0 | 0 | 65 | P19314 | 293 | 21 | 1 | 0 | 120 | Q5R2J2 | 289 | 19 | 2 | 0 |
| 11 | O52631 | 288 | 20 | 2 | 0 | 66 | P00361 | 284 | 19 | 3 | 0 | 121 | Q757I2 | 283 | 24 | 0 | 0 |
| 12 | P34783 | 303 | 27 | 1 | 0 | 67 | Q5JHB5 | 298 | 18 | 0 | 0 | 122 | P51469 | 292 | 19 | 1 | 0 |
| 13 | A6UUN9 | 290 | 25 | 0 | 0 | 68 | B6YUU0 | 298 | 18 | 0 | 0 | 123 | Q5XJ10 | 298 | 16 | 1 | 0 |
| 14 | Q6M0E6 | 298 | 21 | 0 | 0 | 69 | C6A312 | 302 | 16 | 0 | 0 | 124 | P00355 | 299 | 17 | 0 | 0 |
| 15 | Q00640 | 289 | 23 | 1 | 0 | 70 | Q9V1P1 | 288 | 23 | 0 | 0 | 125 | P10096 | 299 | 17 | 0 | 0 |
| 16 | P35143 | 288 | 22 | 2 | 0 | 71 | O59494 | 290 | 22 | 0 | 0 | 126 | P80534 | 329 | 17 | 0 | 0 |
| 17 | P54117 | 294 | 20 | 1 | 0 | 72 | Q9Z7T0 | 295 | 20 | 0 | 0 | 127 | Q9N2D5 | 297 | 18 | 0 | 0 |
| 18 | P20445 | 288 | 24 | 0 | 0 | 73 | P75358 | 285 | 20 | 4 | 0 | 128 | Q28259 | 297 | 18 | 0 | 0 |
| 19 | Q12552 | 275 | 29 | 1 | 0 | 74 | Q5V061 | 319 | 16 | 1 | 0 | 129 | Q4KYY3 | 297 | 18 | 0 | 0 |
| 20 | Q96US8 | 283 | 26 | 1 | 0 | 75 | O28542 | 300 | 20 | 0 | 0 | 130 | A3FKF7 | 297 | 18 | 0 | 0 |
| 21 | P17721 | 280 | 22 | 3 | 0 | 76 | Q48335 | 284 | 19 | 3 | 1 | 131 | P46406 | 295 | 19 | 0 | 0 |
| 22 | A2BLC6 | 304 | 18 | 1 | 0 | 77 | P26988 | 288 | 22 | 0 | 0 | 132 | P04797 | 297 | 18 | 0 | 0 |
| 23 | Q9Y796 | 292 | 21 | 1 | 0 | 78 | Q58546 | 297 | 23 | 0 | 0 | 133 | P17244 | 299 | 17 | 0 | 0 |
| 24 | Q9HGY7 | 285 | 25 | 1 | 0 | 79 | Q01982 | 292 | 21 | 1 | 0 | 134 | P16858 | 299 | 17 | 0 | 0 |
| 25 | P23722 | 281 | 24 | 2 | 0 | 80 | Q9PJN6 | 282 | 25 | 1 | 0 | 135 | O57479 | 296 | 17 | 1 | 0 |
| 26 | P09316 | 298 | 18 | 1 | 0 | 81 | B0B879 | 279 | 26 | 1 | 0 | 136 | P00356 | 296 | 17 | 1 | 0 |
| 27 | P0A038 | 292 | 19 | 2 | 0 | 82 | P0CE13 | 281 | 25 | 1 | 0 | 137 | Q05025 | 296 | 17 | 1 | 0 |
| 28 | Q00584 | 290 | 22 | 1 | 0 | 83 | O83816 | 297 | 25 | 1 | 0 | 138 | Q4J940 | 303 | 17 | 2 | 0 |
| 29 | O13507 | 286 | 26 | 0 | 0 | 84 | O42259 | 295 | 17 | 2 | 0 | 139 | Q9UW96 | 288 | 22 | 1 | 0 |
| 30 | Q9UR38 | 291 | 23 | 0 | 0 | 85 | P87197 | 285 | 25 | 1 | 0 | 140 | Q92243 | 283 | 27 | 0 | 0 |
| 31 | P32638 | 292 | 21 | 1 | 0 | 86 | Q3IUT3 | 303 | 16 | 1 | 0 | 141 | A6VIW4 | 298 | 21 | 0 | 0 |
| 32 | P00362 | 276 | 25 | 3 | 0 | 87 | Q4U3L0 | 290 | 20 | 1 | 0 | 142 | A9A6M9 | 298 | 21 | 0 | 0 |
| 33 | P15115 | 280 | 25 | 2 | 0 | 88 | Q01597 | 291 | 19 | 1 | 0 | 143 | A4FZL2 | 300 | 20 | 0 | 0 |
| 34 | Q971K2 | 294 | 23 | 1 | 0 | 89 | O01360 | 290 | 23 | 1 | 0 | 144 | P08439 | 288 | 21 | 1 | 0 |
| 35 | Q5RAB4 | 299 | 18 | 0 | 0 | 90 | P48812 | 288 | 24 | 1 | 0 | 145 | P19089 | 286 | 24 | 1 | 0 |
| 36 | P04406 | 299 | 18 | 0 | 0 | 91 | P09317 | 287 | 20 | 2 | 1 | 146 | P54118 | 285 | 25 | 1 | 0 |
| 37 | A7IB57 | 296 | 22 | 1 | 0 | 92 | P20287 | 287 | 21 | 3 | 0 | 147 | Q8WZN0 | 289 | 23 | 1 | 0 |
| 38 | A3CYG8 | 305 | 18 | 0 | 0 | 93 | P51009 | 302 | 12 | 3 | 0 | 148 | Q9HFX1 | 292 | 21 | 1 | 0 |
| 39 | Q2FNA2 | 294 | 21 | 1 | 0 | 94 | Q9YFS9 | 288 | 26 | 1 | 0 | 149 | P29497 | 290 | 22 | 1 | 0 |
| 40 | A4YHA5 | 299 | 19 | 1 | 0 | 95 | O59841 | 281 | 27 | 0 | 0 | 150 | Q9P8C0 | 289 | 21 | 2 | 0 |
| 41 | A1RV79 | 305 | 18 | 1 | 0 | 96 | P52694 | 285 | 22 | 1 | 0 | 151 | P28844 | 286 | 24 | 1 | 0 |
| 42 | Q59199 | 290 | 20 | 1 | 0 | 97 | Q07234 | 281 | 24 | 1 | 0 | 152 | Q8X1X3 | 290 | 21 | 2 | 0 |
| 43 | Q6L125 | 297 | 19 | 2 | 0 | 98 | Q89AK1 | 292 | 19 | 1 | 0 | 153 | Q1DTF9 | 295 | 18 | 2 | 0 |
| 44 | A3MTU1 | 301 | 17 | 3 | 0 | 99 | I2BA89 | 291 | 20 | 0 | 0 | 154 | Q8J1H3 | 296 | 19 | 1 | 0 |
| 45 | P39460 | 298 | 21 | 0 | 0 | 100 | B7LQ20 | 292 | 18 | 1 | 0 | 155 | P0CN74 | 291 | 21 | 2 | 0 |
| 46 | B0R2M2 | 294 | 16 | 3 | 0 | 101 | P0A9B3 | 290 | 19 | 1 | 0 | 156 | P0C0G7 | 288 | 24 | 0 | 0 |
| 47 | Q97BJ8 | 287 | 24 | 1 | 0 | 102 | P0A1P0 | 290 | 19 | 1 | 0 | 157 | P0C0G6 | 290 | 23 | 0 | 0 |
| 48 | Q9HJ69 | 304 | 17 | 0 | 0 | 103 | P57384 | 284 | 21 | 2 | 0 | 158 | P0DB18 | 288 | 24 | 0 | 0 |
| 49 | Q8ZWK7 | 306 | 19 | 0 | 0 | 104 | P27726 | 312 | 8 | 2 | 0 | 159 | Q59906 | 289 | 22 | 1 | 0 |
| 50 | A4WIW2 | 305 | 18 | 1 | 0 | 105 | P29272 | 288 | 18 | 3 | 0 | 160 | P52987 | 291 | 20 | 2 | 0 |
| 51 | B1YD06 | 307 | 17 | 1 | 0 | 106 | Q01651 | 290 | 19 | 2 | 0 | 161 | Q8TFJ2 | 292 | 21 | 1 | 0 |
| 52 | O27090 | 297 | 20 | 0 | 0 | 107 | Q9Z518 | 292 | 19 | 2 | 0 | 162 | Q94469 | 291 | 22 | 0 | 0 |
| 53 | C3MWN0 | 300 | 20 | 0 | 0 | 108 | A0QWW2 | 297 | 17 | 3 | 0 | 163 | P53430 | 289 | 24 | 0 | 0 |
| 54 | C3MQN0 | 300 | 20 | 0 | 0 | 109 | Q59309 | 291 | 20 | 1 | 0 | 164 | P56649 | 285 | 24 | 0 | 0 |
| 55 | C3N6F0 | 300 | 20 | 0 | 0 | 110 | Q12XM4 | 300 | 16 | 1 | 0 | 165 | P00357 | 283 | 25 | 0 | 0 |

Poly-strings of length 4 were identified in sequences *Q*48335 and *P* 09317, consisting of four consecutive alanines (AAAA) and four consecutive arginines (RRRR), respectively, each occurring with a frequency of 1. The highest observed frequency for a poly-string of length 3 was 4, found in sequence P75358, comprising one AAA, two KKK, and one TTT repeats. Tables 6, 7, and 8 list sequences containing poly-strings of length 3 with frequencies of 3, 2, and 1, respectively. These tables include only those amino acids for which at least one non-zero value was observed. It is noteworthy that among the poly-strings of length 3, only alanine (AAA) appeared with a frequency greater than 1, as shown in Tables 6 and 7. A total of 64 GAPDH sequences exhibited only a single occurrence of poly-string of length 3 across all amino acids (Tables 5 and 8). FFF, GGG, and KKK were found only in Q2FNA2, Q59906 and Q97BJ8 respectively among all 165 sequences (Table 8). Notably, no poly-string of length 3 was detected in 62 GAPDH sequences. None of 165 sequences contained CCC, HHH, MMM, PPP, QQQ, RRR, WWW and YYY.

**Table 6.** Sequences having poly-strings of of amino acids length 3 with frequency 3 considering all twenty amino acids.

| Uniport ID | AAA | DDD | EEE | LLL | NNN | SSS | TTT | VVV |
| --- | --- | --- | --- | --- | --- | --- | --- | --- |
| O67161 | 1 | 0 | 0 | 0 | 0 | 0 | 1 | 1 |
| P17721 | 1 | 0 | 0 | 1 | 0 | 0 | 1 | 0 |
| P00362 | 2 | 0 | 0 | 0 | 0 | 0 | 1 | 0 |
| A3MTU1 | 1 | 0 | 0 | 1 | 0 | 0 | 0 | 1 |
| B0R2M2 | 1 | 1 | 0 | 0 | 0 | 0 | 0 | 1 |
| P55971 | 1 | 0 | 0 | 1 | 0 | 0 | 1 | 0 |
| Q9ZKT0 | 1 | 0 | 0 | 1 | 0 | 0 | 1 | 0 |
| P00361 | 2 | 0 | 0 | 0 | 0 | 0 | 1 | 0 |
| Q48335 | 2 | 1 | 0 | 0 | 0 | 0 | 0 | 0 |
| P20287 | 1 | 0 | 0 | 0 | 1 | 1 | 0 | 0 |
| P51009 | 3 | 0 | 0 | 0 | 0 | 0 | 0 | 0 |
| P29272 | 3 | 0 | 0 | 0 | 0 | 0 | 0 | 0 |
| A0QWW2 | 2 | 0 | 0 | 0 | 0 | 0 | 0 | 1 |
| Q59800 | 2 | 0 | 0 | 0 | 0 | 0 | 1 | 0 |
| P46713 | 1 | 0 | 1 | 0 | 0 | 0 | 0 | 1 |
| P94915 | 2 | 0 | 0 | 0 | 0 | 0 | 0 | 1 |
| P9WN83 | 2 | 0 | 0 | 0 | 0 | 0 | 0 | 1 |

**Table 7.** Sequences having poly-strings of amino acids length 3 with frequency 2 considering all twenty amino acids.

| Uniport ID | AAA | DDD | KKK | III | LLL | NNN | SSS | TTT | VVV |
| --- | --- | --- | --- | --- | --- | --- | --- | --- | --- |
| P47543 | 1 | 0 | 0 | 0 | 0 | 0 | 0 | 1 | 0 |
| Q9N655 | 1 | 0 | 0 | 0 | 1 | 0 | 0 | 0 | 0 |
| O52631 | 1 | 0 | 0 | 1 | 0 | 0 | 0 | 0 | 0 |
| P35143 | 0 | 0 | 0 | 0 | 0 | 1 | 0 | 1 | 0 |
| P23722 | 1 | 0 | 0 | 0 | 0 | 0 | 0 | 0 | 1 |
| P0A038 | 1 | 1 | 0 | 0 | 0 | 0 | 0 | 0 | 0 |
| P15115 | 1 | 0 | 0 | 0 | 0 | 0 | 0 | 0 | 1 |
| Q6L125 | 0 | 1 | 0 | 1 | 0 | 0 | 0 | 0 | 0 |
| P46795 | 1 | 0 | 0 | 1 | 0 | 0 | 0 | 0 | 0 |
| O42259 | 0 | 0 | 0 | 0 | 1 | 0 | 0 | 0 | 1 |
| P09317 | 1 | 0 | 0 | 0 | 0 | 1 | 0 | 0 | 0 |
| P57384 | 0 | 0 | 1 | 0 | 0 | 0 | 0 | 0 | 1 |
| P27726 | 2 | 0 | 0 | 0 | 0 | 0 | 0 | 0 | 0 |
| Q01651 | 1 | 1 | 0 | 0 | 0 | 0 | 0 | 0 | 0 |
| Q9Z518 | 1 | 0 | 0 | 0 | 0 | 0 | 0 | 1 | 0 |
| Q5R2J2 | 1 | 0 | 0 | 0 | 0 | 0 | 0 | 1 | 0 |
| Q4J940 | 0 | 0 | 1 | 0 | 1 | 0 | 0 | 0 | 0 |
| Q9P8C0 | 1 | 0 | 0 | 0 | 0 | 0 | 0 | 1 | 0 |
| Q8X1X3 | 0 | 0 | 0 | 0 | 0 | 0 | 1 | 1 | 0 |
| Q1DTF9 | 1 | 0 | 0 | 0 | 0 | 0 | 0 | 1 | 0 |
| P0CN74 | 1 | 0 | 0 | 0 | 0 | 0 | 0 | 1 | 0 |
| P52987 | 2 | 0 | 0 | 0 | 0 | 0 | 0 | 0 | 0 |

**Table 8.**
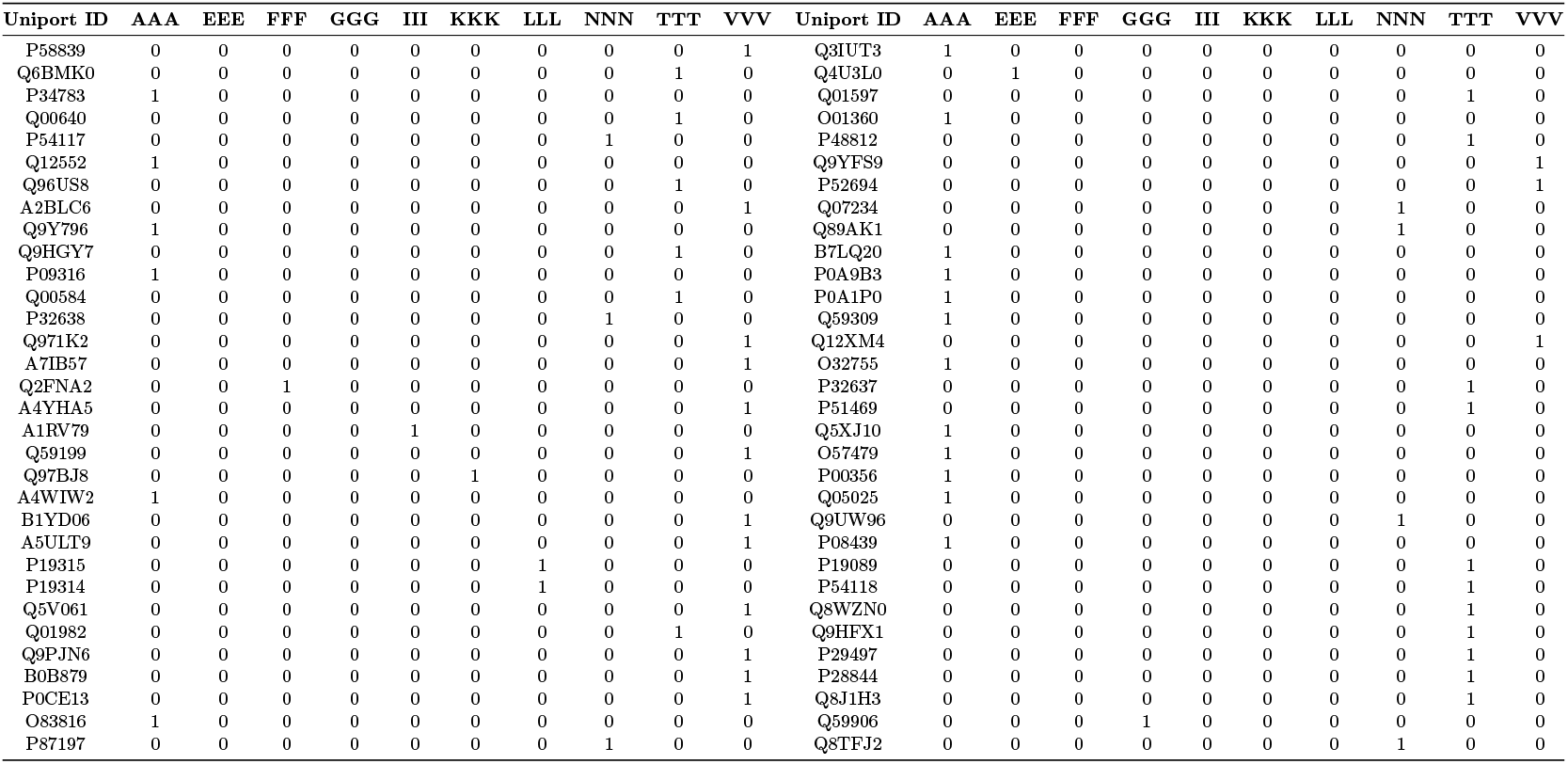
Sequences having poly-strings of of amino acids length 3 with frequency 1 considering all twenty amino acids.

Considering all amino acids across 165 sequences, the maximum count of poly-string of length 2 was 8, which was noted for EE (in P58839) and VV (in O13507, P20287 and P51469)(**Supplementary File-1**). All GAPDH sequences had at least one TT, with a maximum of five occurrences in a single sequence. Furthermore, poly-strings of amino acids of length 2, other than glutamic acid, had all the frequencies ranging from 1 to their respective maximum. None of the GAPDH sequences showed EE with frequencies 5, 6, or 7. WW was not present in any of the 165 sequences.

- Two sequences viz. Q01982 and Q8NK47 possessed CC with frequency 1 and no other sequence possessed CC.
- One sequence (Q01651) had two FF, whereas twenty sequences had FF with frequency 1. Remaining sequences did not possess any FF.
- One sequence (Q9YFS9) contained two HH, whereas fifty-two sequences contained HH with frequency 1. Remaining sequences did not possess any HH.
- MM, PP, and YY were present in only 14, 11, and 22 GAPDH sequences respectively, all with frequency one. Rest 151, 154 and 143 sequences did not have any MM, PP, and YY respectively.
- VV was absent only in two sequences viz. P55971 and Q9ZKT0.

### 4.5 Polar, non-polar residue profiles of GAPDH moonlighting proteins

Figure 6 shows percentage distributions of polar and non-polar residues in each GAPDH sequence. The ratio of polar to non-polar residues was 0.943 *±* 0.07. A4WIW2 (Q6L125) exhibited the lowest (the highest) ratio of 0.801 (1.243). Furthermore, exactly identical percentage (50%) of polar and non-polar residues was noticed in four sequences viz. Q00640, Q5V061, Q01651, and A4FZL2.

**Figure 6.**
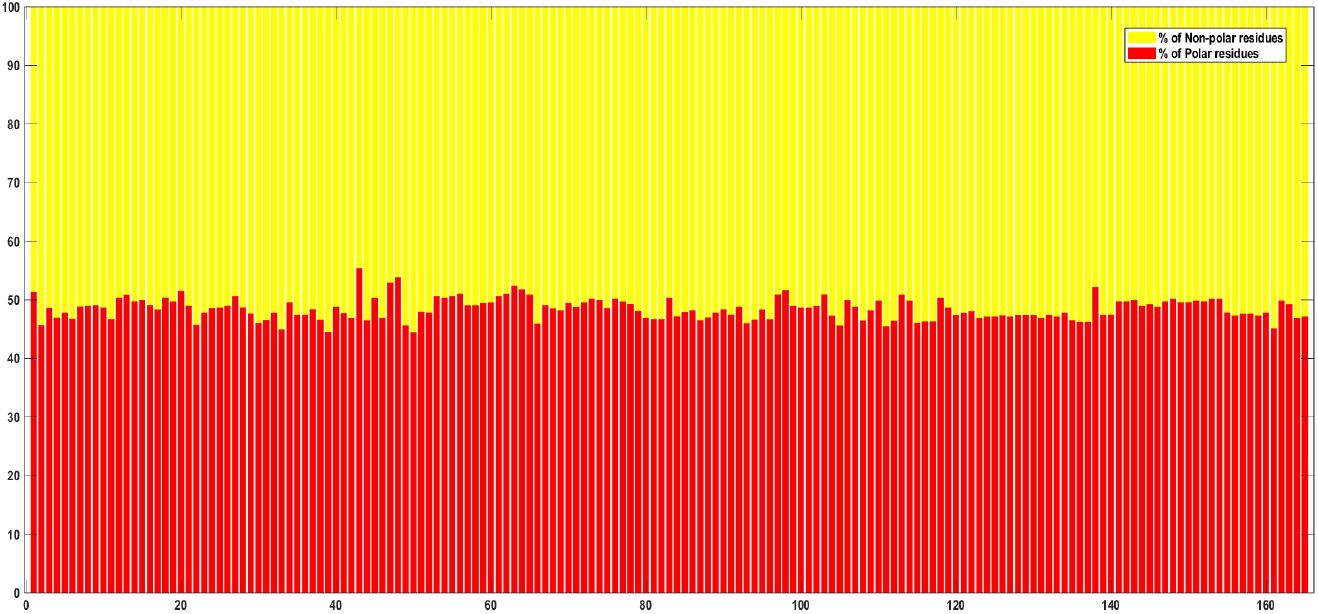
Percentages of polar, non-polar residues in GAPDH moonlighting proteins

Altogether 165 GAPDH sequences could be grouped into three disjoint groups as follows:

- **Group-I**: {50, 39, 33, 161, 111, 49, 105, 2, 22, 66, 93, 30, 115, 136, 137, 116, 117, 112, 108, 44, 87, 135, 31, 38, 94, 96, 11, 81, 82, 6, 42, 123, 131, 164, 46, 80, 4, 88, 124, 125, 127, 133, 165, 84, 104, 156, 159}
- **Group-II**: {126, 120, 128, 129, 130, 132, 35, 36, 139, 91, 140, 157, 158, 29, 41, 121, 134, 5, 32, 23, 52, 160, 89, 155, 85, 51, 122, 79, 69, 109, 86, 95, 17, 90, 37, 68, 24, 75, 3, 100, 101, 10, 25, 119, 28, 71, 107, 92, 146, 40, 7, 99, 102, 21, 144, 8, 26, 57, 58, 9, 67, 16, 145, 163, 78, 59, 70, 60, 72, 149, 150, 34, 77, 19, 147, 152, 14, 141, 142, 110, 162, 114, 151, 15, 74, 106, 143, 73, 148, 153, 154, 76, 12, 83, 45, 54, 118, 18, 53, 55, 27, 61, 13, 65, 113, 97, 103, 56, 62, 1, 20, 98, 64, 138, 63, 47, 48,}
- **Group-III**: {43}

GAPDH sequences of the **Group-I** were found to be *moderately hydrophobic*, whereas members of the **Group-II** were *balanced*. GAPDH sequence 43 (Q6L125) stood out to be *moderately hydrophilic* in nature.

#### 4.5.1 Homogeneous poly-string frequency of polar, non-polar residues in GAPDH sequences

Among all 165 GAPDH sequences, the longest homogeneous poly-strings observed for polar and non-polar residues were 13 and 11, respectively (Tables 9, 10, and 11). Notably, none of the sequences contained polar residue poly-strings of lengths 11 or 12, nor non-polar residue poly-strings of lengths 9 or 10.

**Table 9.** Frequency of homogeneous poly-string of polar and non-polar residues of *GAPDH* moonlighting proteins.

| Sl No | Uniport ID | Homogeneous poly strings of polar residues |  |  |  |  |  |  |  |  |  |  | Homogeneous poly strings of non-polar residues |  |  |  |  |  |  |  |  |
| --- | --- | --- | --- | --- | --- | --- | --- | --- | --- | --- | --- | --- | --- | --- | --- | --- | --- | --- | --- | --- | --- |
|  |  | *l*=1 | *l*=2 | *l*=3 | *l*=4 | *l*=5 | *l*=6 | *l*=7 | *l*=8 | *l*=9 | *l*=10 | *l*=13 | *l*=1 | *l*=2 | *l*=3 | *l*=4 | *l*=5 | *l*=6 | *l*=7 | *l*=8 | *l*=11 |
| 1 | P47543 | 48 | 17 | 12 | 8 | 2 | 1 | 1 | 0 | 0 | 0 | 0 | 46 | 24 | 14 | 4 | 1 | 0 | 1 | 0 | 0 |
| 2 | Q9N655 | 55 | 17 | 9 | 7 | 1 | 1 | 0 | 0 | 0 | 0 | 0 | 38 | 23 | 19 | 7 | 3 | 0 | 0 | 0 | 0 |
| 3 | P58839 | 49 | 24 | 16 | 1 | 3 | 1 | 0 | 0 | 0 | 0 | 0 | 45 | 27 | 15 | 4 | 4 | 0 | 0 | 0 | 0 |
| 4 | P44304 | 58 | 16 | 9 | 5 | 2 | 2 | 0 | 0 | 0 | 0 | 0 | 44 | 25 | 15 | 5 | 3 | 1 | 0 | 0 | 0 |
| 5 | Q92211 | 47 | 17 | 9 | 6 | 2 | 3 | 0 | 0 | 0 | 0 | 0 | 35 | 25 | 17 | 4 | 3 | 0 | 0 | 1 | 0 |
| 6 | Q6CCU7 | 58 | 16 | 7 | 5 | 3 | 2 | 0 | 0 | 0 | 0 | 0 | 46 | 21 | 17 | 3 | 3 | 1 | 0 | 1 | 0 |
| 7 | O67161 | 46 | 23 | 7 | 5 | 2 | 4 | 0 | 0 | 0 | 0 | 0 | 39 | 19 | 23 | 6 | 1 | 0 | 0 | 0 | 0 |
| 8 | Q6BMK0 | 45 | 14 | 10 | 8 | 1 | 4 | 0 | 0 | 0 | 0 | 0 | 33 | 23 | 19 | 3 | 3 | 0 | 0 | 1 | 0 |
| 9 | Q9UVC0 | 42 | 18 | 9 | 5 | 3 | 4 | 0 | 0 | 0 | 0 | 0 | 33 | 22 | 18 | 4 | 3 | 0 | 0 | 1 | 0 |
| 10 | Q92263 | 51 | 17 | 6 | 5 | 2 | 1 | 1 | 2 | 0 | 0 | 0 | 39 | 22 | 18 | 4 | 2 | 0 | 0 | 1 | 0 |
| 11 | O52631 | 51 | 22 | 7 | 3 | 2 | 3 | 0 | 0 | 0 | 0 | 0 | 38 | 21 | 20 | 7 | 2 | 0 | 0 | 0 | 0 |
| 12 | P34783 | 49 | 14 | 7 | 8 | 5 | 3 | 0 | 1 | 0 | 0 | 0 | 37 | 23 | 18 | 6 | 2 | 0 | 0 | 1 | 0 |
| 13 | A6UUN9 | 41 | 23 | 15 | 2 | 4 | 1 | 1 | 0 | 0 | 0 | 0 | 37 | 33 | 8 | 6 | 2 | 1 | 0 | 0 | 0 |
| 14 | Q6M0E6 | 44 | 25 | 8 | 5 | 5 | 1 | 0 | 0 | 0 | 0 | 0 | 38 | 30 | 12 | 4 | 3 | 1 | 0 | 0 | 0 |
| 15 | Q00640 | 41 | 19 | 7 | 7 | 3 | 3 | 0 | 1 | 0 | 0 | 0 | 34 | 22 | 15 | 7 | 2 | 0 | 0 | 1 | 0 |
| 16 | P35143 | 48 | 17 | 9 | 6 | 3 | 3 | 0 | 0 | 0 | 0 | 0 | 36 | 24 | 19 | 5 | 1 | 1 | 0 | 0 | 0 |
| 17 | P54117 | 50 | 19 | 6 | 6 | 3 | 3 | 0 | 0 | 0 | 0 | 0 | 35 | 26 | 20 | 4 | 1 | 1 | 0 | 0 | 0 |
| 18 | P20445 | 45 | 15 | 7 | 7 | 5 | 2 | 0 | 1 | 0 | 0 | 0 | 34 | 26 | 13 | 6 | 2 | 0 | 0 | 1 | 0 |
| 19 | Q12552 | 41 | 15 | 9 | 6 | 5 | 2 | 0 | 1 | 0 | 0 | 0 | 32 | 22 | 16 | 5 | 2 | 0 | 1 | 1 | 0 |
| 20 | Q96US8 | 50 | 18 | 6 | 8 | 4 | 3 | 0 | 0 | 0 | 0 | 0 | 40 | 29 | 15 | 4 | 1 | 0 | 0 | 0 | 0 |
| 21 | P17721 | 46 | 17 | 7 | 3 | 6 | 2 | 0 | 1 | 0 | 0 | 0 | 39 | 14 | 21 | 7 | 1 | 0 | 1 | 0 | 0 |
| 22 | A2BLC6 | 52 | 25 | 8 | 4 | 3 | 0 | 0 | 0 | 0 | 0 | 0 | 39 | 29 | 15 | 7 | 2 | 1 | 0 | 0 | 0 |
| 23 | Q9Y796 | 49 | 18 | 8 | 5 | 4 | 2 | 0 | 0 | 0 | 0 | 0 | 38 | 23 | 18 | 4 | 3 | 0 | 1 | 0 | 0 |
| 24 | Q9HGY7 | 52 | 10 | 9 | 6 | 3 | 3 | 0 | 1 | 0 | 0 | 0 | 37 | 22 | 16 | 5 | 2 | 0 | 1 | 1 | 0 |
| 25 | P23722 | 50 | 17 | 8 | 6 | 1 | 3 | 0 | 1 | 0 | 0 | 0 | 37 | 23 | 21 | 4 | 2 | 0 | 0 | 0 | 0 |
| 26 | P09316 | 47 | 18 | 12 | 5 | 4 | 1 | 0 | 0 | 0 | 0 | 0 | 41 | 20 | 21 | 4 | 1 | 0 | 1 | 0 | 0 |
| 27 | P0A038 | 47 | 21 | 8 | 7 | 1 | 4 | 0 | 0 | 0 | 0 | 0 | 43 | 19 | 20 | 5 | 1 | 0 | 0 | 0 | 0 |
| 28 | Q00584 | 58 | 14 | 5 | 5 | 5 | 3 | 0 | 0 | 0 | 0 | 0 | 42 | 23 | 18 | 5 | 1 | 1 | 0 | 0 | 0 |
| 29 | O13507 | 57 | 16 | 8 | 5 | 2 | 3 | 0 | 0 | 0 | 0 | 0 | 45 | 20 | 20 | 4 | 2 | 1 | 0 | 0 | 0 |
| 30 | Q9UR38 | 50 | 16 | 8 | 5 | 2 | 2 | 1 | 0 | 0 | 0 | 0 | 35 | 18 | 23 | 5 | 2 | 2 | 0 | 0 | 0 |
| 31 | P32638 | 47 | 15 | 10 | 9 | 0 | 1 | 0 | 1 | 0 | 0 | 0 | 33 | 22 | 21 | 3 | 4 | 0 | 0 | 1 | 0 |
| 32 | P00362 | 50 | 19 | 7 | 4 | 3 | 2 | 0 | 1 | 0 | 0 | 0 | 41 | 17 | 21 | 5 | 2 | 0 | 1 | 0 | 0 |
| 33 | P15115 | 55 | 23 | 3 | 5 | 3 | 1 | 0 | 0 | 0 | 0 | 0 | 45 | 19 | 17 | 6 | 2 | 1 | 0 | 0 | 1 |
| 34 | Q971K2 | 46 | 22 | 16 | 4 | 2 | 1 | 0 | 0 | 0 | 0 | 0 | 43 | 31 | 8 | 7 | 2 | 1 | 0 | 0 | 0 |
| 35 | Q5RAB4 | 50 | 19 | 6 | 5 | 1 | 2 | 0 | 2 | 0 | 0 | 0 | 35 | 25 | 17 | 5 | 1 | 0 | 1 | 1 | 0 |
| 36 | P04406 | 50 | 19 | 6 | 5 | 1 | 2 | 0 | 2 | 0 | 0 | 0 | 35 | 25 | 17 | 5 | 1 | 0 | 1 | 1 | 0 |
| 37 | A7IB57 | 46 | 21 | 13 | 6 | 3 | 0 | 0 | 0 | 0 | 0 | 0 | 36 | 30 | 13 | 9 | 0 | 1 | 0 | 0 | 0 |
| 38 | A3CYG8 | 51 | 17 | 10 | 5 | 2 | 1 | 0 | 1 | 0 | 0 | 0 | 37 | 27 | 13 | 7 | 2 | 1 | 0 | 1 | 0 |
| 39 | Q2FNA2 | 54 | 15 | 11 | 6 | 2 | 0 | 0 | 0 | 0 | 0 | 0 | 36 | 29 | 11 | 7 | 4 | 1 | 1 | 0 | 0 |
| 40 | A4YHA5 | 51 | 27 | 9 | 6 | 2 | 0 | 0 | 0 | 0 | 0 | 0 | 48 | 29 | 10 | 8 | 0 | 1 | 0 | 0 | 0 |
| 41 | A1RV79 | 50 | 22 | 5 | 2 | 3 | 2 | 1 | 0 | 0 | 0 | 1 | 37 | 28 | 9 | 9 | 2 | 0 | 2 | 0 | 0 |
| 42 | Q59199 | 52 | 16 | 8 | 8 | 2 | 1 | 0 | 0 | 0 | 0 | 0 | 37 | 23 | 17 | 7 | 3 | 0 | 0 | 0 | 0 |
| 43 | Q6L125 | 46 | 23 | 11 | 5 | 5 | 2 | 1 | 0 | 0 | 0 | 0 | 52 | 27 | 10 | 4 | 0 | 0 | 0 | 0 | 0 |
| 44 | A3MTU1 | 53 | 23 | 6 | 3 | 3 | 1 | 0 | 0 | 0 | 1 | 0 | 40 | 30 | 8 | 9 | 2 | 0 | 2 | 0 | 0 |
| 45 | P39460 | 46 | 19 | 14 | 6 | 3 | 1 | 0 | 0 | 0 | 0 | 0 | 44 | 28 | 8 | 7 | 1 | 2 | 0 | 0 | 0 |
| 46 | B0R2M2 | 50 | 23 | 11 | 3 | 2 | 1 | 0 | 0 | 0 | 0 | 0 | 42 | 25 | 14 | 7 | 2 | 1 | 0 | 0 | 0 |
| 47 | Q97BJ8 | 47 | 19 | 9 | 3 | 7 | 2 | 0 | 1 | 0 | 0 | 0 | 42 | 28 | 14 | 3 | 0 | 0 | 1 | 0 | 0 |
| 48 | Q9HJ69 | 50 | 19 | 12 | 3 | 8 | 1 | 0 | 0 | 0 | 0 | 0 | 48 | 32 | 12 | 2 | 0 | 0 | 0 | 0 | 0 |
| 49 | Q8ZWK7 | 54 | 26 | 6 | 1 | 3 | 1 | 0 | 1 | 0 | 0 | 0 | 41 | 30 | 12 | 6 | 1 | 0 | 3 | 0 | 0 |
| 50 | A4WIW2 | 53 | 26 | 5 | 2 | 2 | 1 | 0 | 0 | 1 | 0 | 0 | 40 | 27 | 11 | 8 | 1 | 1 | 3 | 0 | 0 |
| 51 | B1YD06 | 49 | 26 | 9 | 2 | 2 | 1 | 0 | 0 | 0 | 0 | 1 | 42 | 28 | 9 | 8 | 3 | 0 | 1 | 0 | 0 |
| 52 | O27090 | 48 | 22 | 4 | 8 | 5 | 0 | 0 | 0 | 0 | 0 | 0 | 37 | 29 | 11 | 8 | 2 | 1 | 0 | 0 | 0 |
| 53 | C3MWN0 | 43 | 20 | 16 | 5 | 3 | 1 | 0 | 0 | 0 | 0 | 0 | 43 | 28 | 8 | 7 | 1 | 2 | 0 | 0 | 0 |
| 54 | C3MQN0 | 43 | 21 | 15 | 5 | 3 | 1 | 0 | 0 | 0 | 0 | 0 | 43 | 28 | 8 | 7 | 1 | 1 | 1 | 0 | 0 |
| 55 | C3N6F0 | 43 | 20 | 16 | 5 | 3 | 1 | 0 | 0 | 0 | 0 | 0 | 43 | 28 | 8 | 7 | 1 | 2 | 0 | 0 | 0 |

**Table 10.**
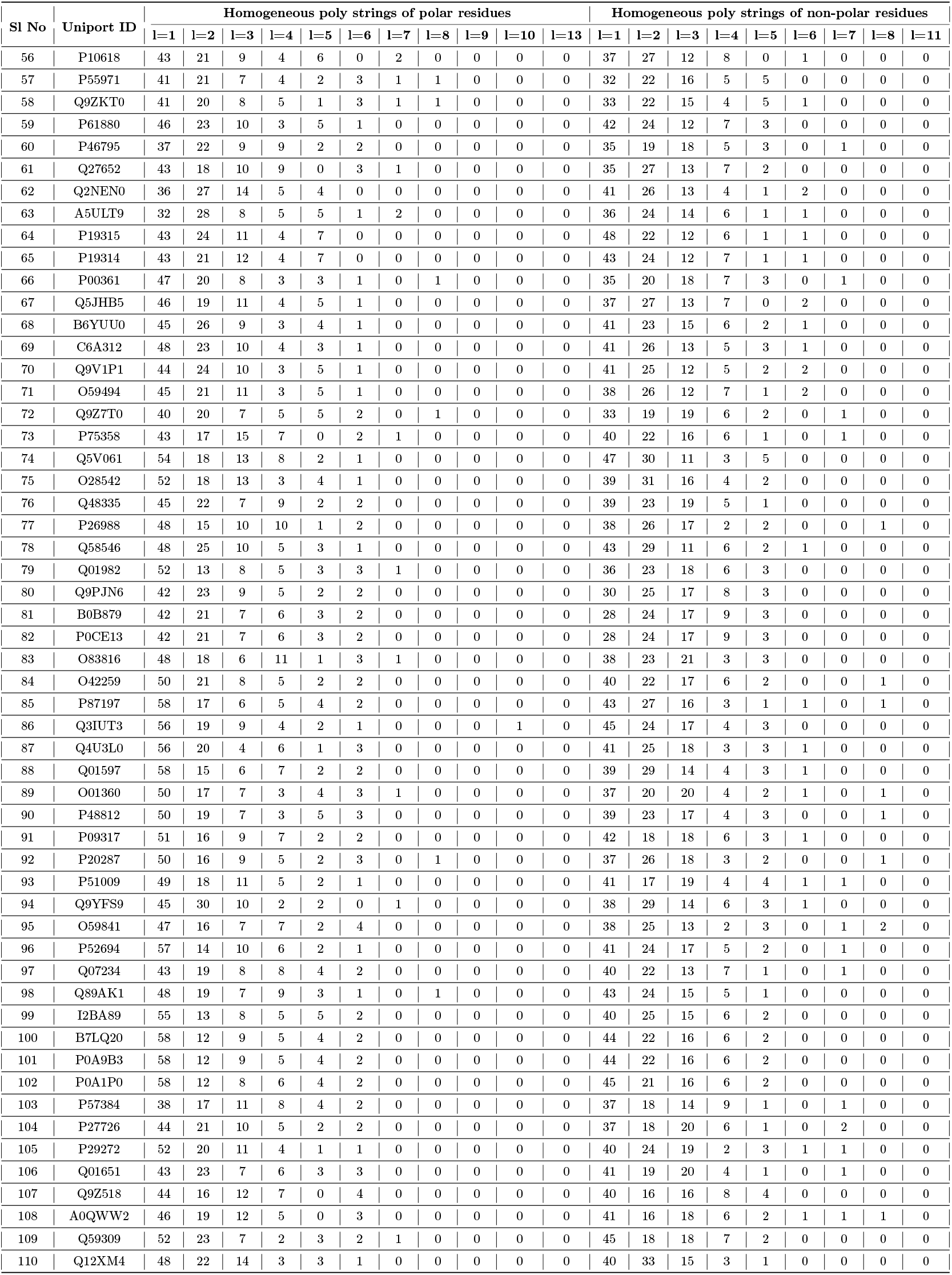
Frequency of homogeneous poly-string of polar and non-polar residues of *GAPDH* moonlighting proteins.

| Sl No | Uniport ID | Homogeneous poly strings of polar residues |  |  |  |  |  |  |  |  |  |  | Homogeneous poly strings of non-polar residues |  |  |  |  |  |  |  |  |
| --- | --- | --- | --- | --- | --- | --- | --- | --- | --- | --- | --- | --- | --- | --- | --- | --- | --- | --- | --- | --- | --- |
|  |  | l=1 | l=2 | l=3 | l=4 | l=5 | l=6 | l=7 | l=8 | l=9 | l=10 | l=13 | l=1 | l=2 | l=3 | l=4 | l=5 | l=6 | l=7 | l=8 | l=11 |
| 56 | P10618 | 43 | 21 | 9 | 4 | 6 | 0 | 2 | 0 | 0 | 0 | 0 | 37 | 27 | 12 | 8 | 0 | 1 | 0 | 0 | 0 |
| 57 | P55971 | 41 | 21 | 7 | 4 | 2 | 3 | 1 | 1 | 0 | 0 | 0 | 32 | 22 | 16 | 5 | 5 | 0 | 0 | 0 | 0 |
| 58 | Q9ZKT0 | 41 | 20 | 8 | 5 | 1 | 3 | 1 | 1 | 0 | 0 | 0 | 33 | 22 | 15 | 4 | 5 | 1 | 0 | 0 | 0 |
| 59 | P61880 | 46 | 23 | 10 | 3 | 5 | 1 | 0 | 0 | 0 | 0 | 0 | 42 | 24 | 12 | 7 | 3 | 0 | 0 | 0 | 0 |
| 60 | P46795 | 37 | 22 | 9 | 9 | 2 | 2 | 0 | 0 | 0 | 0 | 0 | 35 | 19 | 18 | 5 | 3 | 0 | 1 | 0 | 0 |
| 61 | Q27652 | 43 | 18 | 10 | 9 | 0 | 3 | 1 | 0 | 0 | 0 | 0 | 35 | 27 | 13 | 7 | 2 | 0 | 0 | 0 | 0 |
| 62 | Q2NEN0 | 36 | 27 | 14 | 5 | 4 | 0 | 0 | 0 | 0 | 0 | 0 | 41 | 26 | 13 | 4 | 1 | 2 | 0 | 0 | 0 |
| 63 | A5ULT9 | 32 | 28 | 8 | 5 | 5 | 1 | 2 | 0 | 0 | 0 | 0 | 36 | 24 | 14 | 6 | 1 | 1 | 0 | 0 | 0 |
| 64 | P19315 | 43 | 24 | 11 | 4 | 7 | 0 | 0 | 0 | 0 | 0 | 0 | 48 | 22 | 12 | 6 | 1 | 1 | 0 | 0 | 0 |
| 65 | P19314 | 43 | 21 | 12 | 4 | 7 | 0 | 0 | 0 | 0 | 0 | 0 | 43 | 24 | 12 | 7 | 1 | 1 | 0 | 0 | 0 |
| 66 | P00361 | 47 | 20 | 8 | 3 | 3 | 1 | 0 | 1 | 0 | 0 | 0 | 35 | 20 | 18 | 7 | 3 | 0 | 1 | 0 | 0 |
| 67 | Q5JHB5 | 46 | 19 | 11 | 4 | 5 | 1 | 0 | 0 | 0 | 0 | 0 | 37 | 27 | 13 | 7 | 0 | 2 | 0 | 0 | 0 |
| 68 | B6YUU0 | 45 | 26 | 9 | 3 | 4 | 1 | 0 | 0 | 0 | 0 | 0 | 41 | 23 | 15 | 6 | 2 | 1 | 0 | 0 | 0 |
| 69 | C6A312 | 48 | 23 | 10 | 4 | 3 | 1 | 0 | 0 | 0 | 0 | 0 | 41 | 26 | 13 | 5 | 3 | 1 | 0 | 0 | 0 |
| 70 | Q9V1P1 | 44 | 24 | 10 | 3 | 5 | 1 | 0 | 0 | 0 | 0 | 0 | 41 | 25 | 12 | 5 | 2 | 2 | 0 | 0 | 0 |
| 71 | O59494 | 45 | 21 | 11 | 3 | 5 | 1 | 0 | 0 | 0 | 0 | 0 | 38 | 26 | 12 | 7 | 1 | 2 | 0 | 0 | 0 |
| 72 | Q9Z7T0 | 40 | 20 | 7 | 5 | 5 | 2 | 0 | 1 | 0 | 0 | 0 | 33 | 19 | 19 | 6 | 2 | 0 | 1 | 0 | 0 |
| 73 | P75358 | 43 | 17 | 15 | 7 | 0 | 2 | 1 | 0 | 0 | 0 | 0 | 40 | 22 | 16 | 6 | 1 | 0 | 1 | 0 | 0 |
| 74 | Q5V061 | 54 | 18 | 13 | 8 | 2 | 1 | 0 | 0 | 0 | 0 | 0 | 47 | 30 | 11 | 3 | 5 | 0 | 0 | 0 | 0 |
| 75 | O28542 | 52 | 18 | 13 | 3 | 4 | 1 | 0 | 0 | 0 | 0 | 0 | 39 | 31 | 16 | 4 | 2 | 0 | 0 | 0 | 0 |
| 76 | Q48335 | 45 | 22 | 7 | 9 | 2 | 2 | 0 | 0 | 0 | 0 | 0 | 39 | 23 | 19 | 5 | 1 | 0 | 0 | 0 | 0 |
| 77 | P26988 | 48 | 15 | 10 | 10 | 1 | 2 | 0 | 0 | 0 | 0 | 0 | 38 | 26 | 17 | 2 | 2 | 0 | 0 | 1 | 0 |
| 78 | Q58546 | 48 | 25 | 10 | 5 | 3 | 1 | 0 | 0 | 0 | 0 | 0 | 43 | 29 | 11 | 6 | 2 | 1 | 0 | 0 | 0 |
| 79 | Q01982 | 52 | 13 | 8 | 5 | 3 | 3 | 1 | 0 | 0 | 0 | 0 | 36 | 23 | 18 | 6 | 3 | 0 | 0 | 0 | 0 |
| 80 | Q9PJN6 | 42 | 23 | 9 | 5 | 2 | 2 | 0 | 0 | 0 | 0 | 0 | 30 | 25 | 17 | 8 | 3 | 0 | 0 | 0 | 0 |
| 81 | B0B879 | 42 | 21 | 7 | 6 | 3 | 2 | 0 | 0 | 0 | 0 | 0 | 28 | 24 | 17 | 9 | 3 | 0 | 0 | 0 | 0 |
| 82 | P0CE13 | 42 | 21 | 7 | 6 | 3 | 2 | 0 | 0 | 0 | 0 | 0 | 28 | 24 | 17 | 9 | 3 | 0 | 0 | 0 | 0 |
| 83 | O83816 | 48 | 18 | 6 | 11 | 1 | 3 | 1 | 0 | 0 | 0 | 0 | 38 | 23 | 21 | 3 | 3 | 0 | 0 | 0 | 0 |
| 84 | O42259 | 50 | 21 | 8 | 5 | 2 | 2 | 0 | 0 | 0 | 0 | 0 | 40 | 22 | 17 | 6 | 2 | 0 | 0 | 1 | 0 |
| 85 | P87197 | 58 | 17 | 6 | 5 | 4 | 2 | 0 | 0 | 0 | 0 | 0 | 43 | 27 | 16 | 3 | 1 | 1 | 0 | 1 | 0 |
| 86 | Q3IUT3 | 56 | 19 | 9 | 4 | 2 | 1 | 0 | 0 | 0 | 1 | 0 | 45 | 24 | 17 | 4 | 3 | 0 | 0 | 0 | 0 |
| 87 | Q4U3L0 | 56 | 20 | 4 | 6 | 1 | 3 | 0 | 0 | 0 | 0 | 0 | 41 | 25 | 18 | 3 | 3 | 1 | 0 | 0 | 0 |
| 88 | Q01597 | 58 | 15 | 6 | 7 | 2 | 2 | 0 | 0 | 0 | 0 | 0 | 39 | 29 | 14 | 4 | 3 | 1 | 0 | 0 | 0 |
| 89 | O01360 | 50 | 17 | 7 | 3 | 4 | 3 | 1 | 0 | 0 | 0 | 0 | 37 | 20 | 20 | 4 | 2 | 1 | 0 | 1 | 0 |
| 90 | P48812 | 50 | 19 | 7 | 3 | 5 | 3 | 0 | 0 | 0 | 0 | 0 | 39 | 23 | 17 | 4 | 3 | 0 | 0 | 1 | 0 |
| 91 | P09317 | 51 | 16 | 9 | 7 | 2 | 2 | 0 | 0 | 0 | 0 | 0 | 42 | 18 | 18 | 6 | 3 | 1 | 0 | 0 | 0 |
| 92 | P20287 | 50 | 16 | 9 | 5 | 2 | 3 | 0 | 1 | 0 | 0 | 0 | 37 | 26 | 18 | 3 | 2 | 0 | 0 | 1 | 0 |
| 93 | P51009 | 49 | 18 | 11 | 5 | 2 | 1 | 0 | 0 | 0 | 0 | 0 | 41 | 17 | 19 | 4 | 4 | 1 | 1 | 0 | 0 |
| 94 | Q9YFS9 | 45 | 30 | 10 | 2 | 2 | 0 | 1 | 0 | 0 | 0 | 0 | 38 | 29 | 14 | 6 | 3 | 1 | 0 | 0 | 0 |
| 95 | O59841 | 47 | 16 | 7 | 7 | 2 | 4 | 0 | 0 | 0 | 0 | 0 | 38 | 25 | 13 | 2 | 3 | 0 | 1 | 2 | 0 |
| 96 | P52694 | 57 | 14 | 10 | 6 | 2 | 1 | 0 | 0 | 0 | 0 | 0 | 41 | 24 | 17 | 5 | 2 | 0 | 1 | 0 | 0 |
| 97 | Q07234 | 43 | 19 | 8 | 8 | 4 | 2 | 0 | 0 | 0 | 0 | 0 | 40 | 22 | 13 | 7 | 1 | 0 | 1 | 0 | 0 |
| 98 | Q89AK1 | 48 | 19 | 7 | 9 | 3 | 1 | 0 | 1 | 0 | 0 | 0 | 43 | 24 | 15 | 5 | 1 | 0 | 0 | 0 | 0 |
| 99 | I2BA89 | 55 | 13 | 8 | 5 | 5 | 2 | 0 | 0 | 0 | 0 | 0 | 40 | 25 | 15 | 6 | 2 | 0 | 0 | 0 | 0 |
| 100 | B7LQ20 | 58 | 12 | 9 | 5 | 4 | 2 | 0 | 0 | 0 | 0 | 0 | 44 | 22 | 16 | 6 | 2 | 0 | 0 | 0 | 0 |
| 101 | P0A9B3 | 58 | 12 | 9 | 5 | 4 | 2 | 0 | 0 | 0 | 0 | 0 | 44 | 22 | 16 | 6 | 2 | 0 | 0 | 0 | 0 |
| 102 | P0A1P0 | 58 | 12 | 8 | 6 | 4 | 2 | 0 | 0 | 0 | 0 | 0 | 45 | 21 | 16 | 6 | 2 | 0 | 0 | 0 | 0 |
| 103 | P57384 | 38 | 17 | 11 | 8 | 4 | 2 | 0 | 0 | 0 | 0 | 0 | 37 | 18 | 14 | 9 | 1 | 0 | 1 | 0 | 0 |
| 104 | P27726 | 44 | 21 | 10 | 5 | 2 | 2 | 0 | 0 | 0 | 0 | 0 | 37 | 18 | 20 | 6 | 1 | 0 | 2 | 0 | 0 |
| 105 | P29272 | 52 | 20 | 11 | 4 | 1 | 1 | 0 | 0 | 0 | 0 | 0 | 40 | 24 | 19 | 2 | 3 | 1 | 1 | 0 | 0 |
| 106 | Q01651 | 43 | 23 | 7 | 6 | 3 | 3 | 0 | 0 | 0 | 0 | 0 | 41 | 19 | 20 | 4 | 1 | 0 | 1 | 0 | 0 |
| 107 | Q9Z518 | 44 | 16 | 12 | 7 | 0 | 4 | 0 | 0 | 0 | 0 | 0 | 40 | 16 | 16 | 8 | 4 | 0 | 0 | 0 | 0 |
| 108 | A0QWW2 | 46 | 19 | 12 | 5 | 0 | 3 | 0 | 0 | 0 | 0 | 0 | 41 | 16 | 18 | 6 | 2 | 1 | 1 | 1 | 0 |
| 109 | Q59309 | 52 | 23 | 7 | 2 | 3 | 2 | 1 | 0 | 0 | 0 | 0 | 45 | 18 | 18 | 7 | 2 | 0 | 0 | 0 | 0 |
| 110 | Q12XM4 | 48 | 22 | 14 | 3 | 3 | 1 | 0 | 0 | 0 | 0 | 0 | 40 | 33 | 15 | 3 | 1 | 0 | 0 | 0 | 0 |

**Table 11.** Frequency of homogeneous poly-string of polar and non-polar residues of *GAPDH* moonlighting proteins.

| Sl No | Uniport ID | Homogeneous poly strings of polar residues |  |  |  |  |  |  |  |  |  |  | Homogeneous poly strings of non-polar residues |  |  |  |  |  |  |  |  |
| --- | --- | --- | --- | --- | --- | --- | --- | --- | --- | --- | --- | --- | --- | --- | --- | --- | --- | --- | --- | --- | --- |
|  |  | l=1 | l=2 | l=3 | l=4 | l=5 | l=6 | l=7 | l=8 | l=9 | l=10 | l=13 | l=1 | l=2 | l=3 | l=4 | l=5 | l=6 | l=7 | l=8 | l=11 |
| 111 | Q59800 | 49 | 22 | 8 | 2 | 4 | 1 | 0 | 0 | 0 | 0 | 0 | 41 | 18 | 15 | 7 | 1 | 2 | 2 | 0 | 0 |
| 112 | Q8NK47 | 65 | 13 | 5 | 3 | 4 | 3 | 0 | 0 | 0 | 0 | 0 | 48 | 19 | 19 | 6 | 1 | 0 | 0 | 1 | 0 |
| 113 | O32755 | 42 | 21 | 12 | 6 | 2 | 3 | 0 | 0 | 0 | 0 | 0 | 41 | 25 | 14 | 4 | 2 | 0 | 1 | 0 | 0 |
| 114 | P32637 | 49 | 18 | 10 | 5 | 3 | 3 | 0 | 0 | 0 | 0 | 0 | 42 | 20 | 18 | 7 | 1 | 0 | 0 | 0 | 0 |
| 115 | P46713 | 45 | 22 | 9 | 3 | 2 | 3 | 0 | 0 | 0 | 0 | 0 | 40 | 16 | 15 | 9 | 2 | 1 | 2 | 0 | 0 |
| 116 | P94915 | 47 | 20 | 9 | 5 | 1 | 3 | 0 | 0 | 0 | 0 | 0 | 41 | 18 | 15 | 7 | 1 | 2 | 1 | 1 | 0 |
| 117 | P9WN83 | 48 | 18 | 10 | 5 | 1 | 3 | 0 | 0 | 0 | 0 | 0 | 40 | 17 | 17 | 8 | 2 | 0 | 1 | 1 | 0 |
| 118 | A6URW1 | 43 | 24 | 11 | 4 | 5 | 1 | 0 | 0 | 0 | 0 | 0 | 41 | 27 | 12 | 4 | 2 | 2 | 0 | 0 | 0 |
| 119 | Q8PTD3 | 39 | 24 | 12 | 5 | 4 | 0 | 0 | 0 | 0 | 0 | 0 | 34 | 30 | 11 | 7 | 2 | 0 | 1 | 0 | 0 |
| 120 | Q5R2J2 | 52 | 14 | 5 | 7 | 3 | 2 | 0 | 1 | 0 | 0 | 0 | 32 | 28 | 17 | 4 | 1 | 0 | 1 | 1 | 0 |
| 121 | Q757I2 | 48 | 19 | 7 | 4 | 3 | 2 | 0 | 1 | 0 | 0 | 0 | 35 | 22 | 20 | 4 | 2 | 0 | 0 | 1 | 0 |
| 122 | P51469 | 51 | 16 | 7 | 4 | 4 | 2 | 0 | 1 | 0 | 0 | 0 | 36 | 25 | 15 | 6 | 2 | 0 | 0 | 1 | 0 |
| 123 | Q5XJ10 | 53 | 14 | 8 | 4 | 3 | 2 | 0 | 1 | 0 | 0 | 0 | 34 | 27 | 15 | 6 | 1 | 0 | 1 | 1 | 0 |
| 124 | P00355 | 50 | 18 | 6 | 5 | 1 | 2 | 0 | 2 | 0 | 0 | 0 | 33 | 26 | 17 | 5 | 1 | 0 | 1 | 1 | 0 |
| 125 | P10096 | 50 | 18 | 6 | 5 | 1 | 2 | 0 | 2 | 0 | 0 | 0 | 33 | 26 | 17 | 5 | 1 | 0 | 1 | 1 | 0 |
| 126 | P80534 | 52 | 21 | 7 | 5 | 1 | 4 | 0 | 1 | 0 | 0 | 0 | 37 | 27 | 17 | 6 | 2 | 0 | 1 | 1 | 0 |
| 127 | Q9N2D5 | 50 | 18 | 6 | 5 | 1 | 2 | 0 | 2 | 0 | 0 | 0 | 33 | 26 | 17 | 5 | 1 | 0 | 1 | 1 | 0 |
| 128 | Q28259 | 49 | 19 | 6 | 5 | 1 | 2 | 0 | 2 | 0 | 0 | 0 | 34 | 25 | 17 | 5 | 1 | 0 | 1 | 1 | 0 |
| 129 | Q4KYY3 | 49 | 19 | 6 | 5 | 1 | 2 | 0 | 2 | 0 | 0 | 0 | 34 | 25 | 17 | 5 | 1 | 0 | 1 | 1 | 0 |
| 130 | A3FKF7 | 49 | 19 | 6 | 5 | 1 | 2 | 0 | 2 | 0 | 0 | 0 | 34 | 25 | 17 | 5 | 1 | 0 | 1 | 1 | 0 |
| 131 | P46406 | 50 | 19 | 6 | 5 | 2 | 2 | 0 | 1 | 0 | 0 | 0 | 34 | 26 | 17 | 5 | 1 | 0 | 1 | 1 | 0 |
| 132 | P04797 | 50 | 16 | 6 | 5 | 2 | 2 | 0 | 2 | 0 | 0 | 0 | 32 | 26 | 17 | 5 | 1 | 0 | 1 | 1 | 0 |
| 133 | P17244 | 50 | 18 | 6 | 5 | 1 | 2 | 0 | 2 | 0 | 0 | 0 | 33 | 26 | 17 | 5 | 1 | 0 | 1 | 1 | 0 |
| 134 | P16858 | 49 | 19 | 6 | 4 | 2 | 2 | 0 | 2 | 0 | 0 | 0 | 35 | 24 | 17 | 5 | 1 | 0 | 1 | 1 | 0 |
| 135 | O57479 | 53 | 16 | 6 | 4 | 2 | 3 | 0 | 1 | 0 | 0 | 0 | 35 | 24 | 17 | 6 | 1 | 0 | 1 | 1 | 0 |
| 136 | P00356 | 53 | 16 | 6 | 5 | 1 | 3 | 0 | 1 | 0 | 0 | 0 | 34 | 25 | 17 | 6 | 1 | 0 | 1 | 1 | 0 |
| 137 | Q05025 | 53 | 16 | 6 | 5 | 1 | 3 | 0 | 1 | 0 | 0 | 0 | 34 | 25 | 17 | 6 | 1 | 0 | 1 | 1 | 0 |
| 138 | Q4J940 | 44 | 22 | 8 | 9 | 5 | 1 | 0 | 0 | 0 | 0 | 0 | 41 | 34 | 9 | 3 | 2 | 1 | 0 | 0 | 0 |
| 139 | Q9UW96 | 52 | 15 | 9 | 7 | 2 | 2 | 0 | 0 | 0 | 0 | 0 | 35 | 26 | 18 | 5 | 3 | 0 | 0 | 0 | 0 |
| 140 | Q92243 | 52 | 22 | 8 | 7 | 0 | 2 | 0 | 0 | 0 | 0 | 0 | 45 | 19 | 20 | 4 | 1 | 1 | 1 | 0 | 0 |
| 141 | A6VIW4 | 44 | 25 | 8 | 5 | 5 | 1 | 0 | 0 | 0 | 0 | 0 | 38 | 30 | 12 | 4 | 3 | 1 | 0 | 0 | 0 |
| 142 | A9A6M9 | 44 | 25 | 8 | 5 | 5 | 1 | 0 | 0 | 0 | 0 | 0 | 38 | 30 | 12 | 4 | 3 | 1 | 0 | 0 | 0 |
| 143 | A4FZL2 | 44 | 23 | 8 | 5 | 6 | 1 | 0 | 0 | 0 | 0 | 0 | 37 | 30 | 12 | 4 | 3 | 1 | 0 | 0 | 0 |
| 144 | P08439 | 48 | 15 | 7 | 7 | 1 | 4 | 1 | 0 | 0 | 0 | 0 | 37 | 21 | 18 | 5 | 2 | 0 | 1 | 0 | 0 |
| 145 | P19089 | 52 | 16 | 8 | 5 | 4 | 3 | 0 | 0 | 0 | 0 | 0 | 41 | 25 | 16 | 4 | 2 | 1 | 0 | 0 | 0 |
| 146 | P54118 | 52 | 19 | 6 | 6 | 3 | 3 | 0 | 0 | 0 | 0 | 0 | 42 | 26 | 16 | 3 | 1 | 1 | 0 | 1 | 0 |
| 147 | Q8WZN0 | 51 | 19 | 6 | 7 | 3 | 3 | 0 | 0 | 0 | 0 | 0 | 42 | 26 | 17 | 3 | 1 | 0 | 0 | 1 | 0 |
| 148 | Q9HFX1 | 51 | 13 | 7 | 8 | 3 | 4 | 0 | 0 | 0 | 0 | 0 | 38 | 26 | 16 | 3 | 2 | 0 | 0 | 1 | 0 |
| 149 | P29497 | 56 | 16 | 7 | 5 | 4 | 3 | 0 | 0 | 0 | 0 | 0 | 45 | 26 | 16 | 3 | 1 | 0 | 0 | 1 | 0 |
| 150 | Q9P8C0 | 55 | 14 | 8 | 5 | 4 | 2 | 0 | 1 | 0 | 0 | 0 | 41 | 27 | 18 | 2 | 1 | 0 | 0 | 1 | 0 |
| 151 | P28844 | 54 | 15 | 7 | 5 | 4 | 1 | 0 | 1 | 1 | 0 | 0 | 41 | 26 | 17 | 3 | 1 | 0 | 0 | 1 | 0 |
| 152 | Q8X1X3 | 49 | 10 | 6 | 10 | 2 | 4 | 1 | 0 | 0 | 0 | 0 | 34 | 24 | 15 | 5 | 3 | 0 | 0 | 1 | 0 |
| 153 | Q1DTF9 | 47 | 13 | 7 | 7 | 4 | 2 | 1 | 1 | 0 | 0 | 0 | 35 | 22 | 17 | 5 | 2 | 0 | 0 | 1 | 0 |
| 154 | Q8J1H3 | 47 | 14 | 8 | 7 | 3 | 2 | 1 | 1 | 0 | 0 | 0 | 36 | 23 | 16 | 5 | 2 | 0 | 0 | 1 | 0 |
| 155 | P0CN74 | 54 | 17 | 9 | 5 | 3 | 2 | 0 | 0 | 0 | 0 | 0 | 41 | 24 | 17 | 4 | 3 | 1 | 0 | 0 | 0 |
| 156 | P0C0G7 | 52 | 25 | 8 | 4 | 1 | 2 | 0 | 0 | 0 | 0 | 0 | 44 | 23 | 17 | 6 | 1 | 0 | 1 | 0 | 0 |
| 157 | P0C0G6 | 51 | 26 | 8 | 4 | 1 | 2 | 0 | 0 | 0 | 0 | 0 | 44 | 24 | 16 | 6 | 1 | 0 | 1 | 0 | 0 |
| 158 | P0DB18 | 50 | 25 | 9 | 4 | 1 | 2 | 0 | 0 | 0 | 0 | 0 | 43 | 23 | 17 | 6 | 1 | 0 | 1 | 0 | 0 |
| 159 | Q59906 | 51 | 28 | 8 | 4 | 0 | 2 | 0 | 0 | 0 | 0 | 0 | 45 | 24 | 16 | 6 | 1 | 0 | 1 | 0 | 0 |
| 160 | P52987 | 52 | 28 | 8 | 3 | 1 | 2 | 0 | 0 | 0 | 0 | 0 | 47 | 26 | 13 | 4 | 3 | 0 | 1 | 0 | 0 |
| 161 | Q8TFJ2 | 54 | 16 | 10 | 5 | 2 | 1 | 0 | 0 | 0 | 0 | 0 | 40 | 21 | 18 | 3 | 5 | 2 | 0 | 0 | 0 |
| 162 | Q94469 | 45 | 15 | 9 | 6 | 2 | 2 | 0 | 0 | 1 | 1 | 0 | 33 | 24 | 15 | 5 | 3 | 0 | 1 | 0 | 0 |
| 163 | P53430 | 49 | 15 | 11 | 6 | 2 | 2 | 0 | 1 | 0 | 0 | 0 | 36 | 27 | 16 | 4 | 2 | 0 | 1 | 0 | 0 |
| 164 | P56649 | 54 | 20 | 5 | 3 | 2 | 3 | 1 | 0 | 0 | 0 | 0 | 37 | 24 | 20 | 3 | 4 | 0 | 0 | 0 | 0 |
| 165 | P00357 | 51 | 22 | 5 | 3 | 2 | 3 | 1 | 0 | 0 | 0 | 0 | 36 | 24 | 20 | 3 | 4 | 0 | 0 | 0 | 0 |

##### Poly-string of polar residues (ps-P)

Each of two sequences, A1RV79 and B1YD06, contained a poly-string of polar residues of length 13 (Table 9). Only three sequences (A3MTU1, Q3IUT3, and Q94469) had a ps-P of length 10 while length 9 was observed in three sequences (A4WIW2, P28844, and Q94469) (Table 9). Maximum frequencies for length 7 and 8 ps-P in a single sequence were 2. Two sequences contained two ps-Ps of length 7, while twelve sequences contained 8 ps-Ps. No ps-Ps of length 5 was found in seven sequences, while ten sequences did not have ps-Ps of length 6 (Tables 9, 10, and 11).

##### Poly-string of non-polar residues (ps-N)

The sequence P15115 possessed a single ps-N of length 11 (Tables 9, 10, and 11), and no other sequence contains any ps-N of length 11. One sequence, O59841, contained an 8-length ps-N with frequency 2. A single occurrence of 8-length ps-N was observed in 51 GAPDH sequences, while 113 sequences did not possess any such poly-strings. Among the 165 GAPDH sequences, two sequences—Q8ZWK7 and A4WIW2—contained ps-N of length 7 with a frequency of 3. Five sequences contained double occurrence of ps-N of length 7. A single occurrence of 7-length ps-N was observed in fifty-four GAPDH sequences, while one hundred and four sequences do not contain any such poly-strings. Ps-N of length 6 with frequency 2 were found in twelve sequences. Seven sequences—A7IB57, A4YHA5, Q6L125, Q97BJ8, Q9HJ69, P10618, and Q5JHB5—did not possess any ps-N of length 5, whereas the remaining sequences contained at least one such poly-string (Tables 9, 10, and 11). Moreover, ps-N of length 5 with the corresponding highest frequency of 5 were found in each of four sequences—P55971, Q9ZKT0, Q5V061, and Q8TFJ2..

#### 4.5.2. Change response sequences of polar, non-polar profiles in GAPDH moonlighting sequences

Q12552 had the lowest percentages of ‘PN’ (23.28%), ‘NP’ (23.58%) changes, while the highest percentage of ‘PN’ (28.02%) and ‘NP’ (28.02%) changes were noticed in A4YHA5 (Figure 7). Two GAPDH sequences Q6L125 and Q9HJ69 possessed significantly low amount amount of changes in ‘NN’ compared to other sequences with percentages 17.35% and 18.40%, respectively. The highest percentage of ‘PP’ (28.49%) and ‘NN (29.29%) changes were found in A5ULT9 and Q2FNA2, respectively. Seven clusters were formed based on a distance threshold of 3 (Figure 9). The largest cluster comprised of 43 sequences, highlighted in red in Figure 9. Based on the threshold sets to 1, total of 140 moonlighting sequences were grouped into 32 disjoint sets, as illustrated in the dendrogram (Figure 9) and as detailed in Table 12.

**Table 12.** List of sets consist of proximal GAPDH protein sequences based on frequency of changes between adjacent residues from polarnonpolar profiles.

| Sets based on the proximity |  |  |  |  |  |
| --- | --- | --- | --- | --- | --- |
| Set-1 | {57, 58, 9, 8} | Set-9 | {73, 83, 106, 143, 148} | Set-17 | {74, 110, 149} |
| Set-2 | {21, 107, 144, 119, 24, 95} | Set-10 | {3, 75, 28, 100, 101, 102, 86} | Set-18 | {2, 22, 49, 105} |
| Set-3 | {12, 18, 153, 154} | Set-11 | {78, 99, 146} | Set-19 | {39, 50} |
| Set-4 | {15, 60, 152, 162, 72} | Set-12 | {51, 155, 69, 109, 68} | Set-20 | {111, 161} |
| Set-5 | {1, 20, 98, 64} | Set-13 | {7, 92, 10, 25, 71, 26, 67, 16, 163} | Set-21 | {4, 6, 46, 88, 87, 96} |
| Set-6 | {13, 113, 62, 65} | Set-14 | {17, 37, 90} | Set-22 | {29, 140, 158, 85} |
| Set-7 | {56, 97, 61} | Set-15 | {14, 141, 142, 77, 114, 151, 59, 145, 70} | Set-23 | {156, 157, 159} |
| Set-8 | {27, 53, 55, 45, 54, 76, 118} | Set-16 | {34, 150, 147} | Set-24 | {11, 94, 44, 164, 42, 84, 165} |
|  |  |  |  | Set-25 | {5, 41, 134, 89, 121, 79, 122} |
|  |  |  |  | Set-26 | {35, 36, 120, 128, 129, 130, 104, 126, 124, 125, 127, 133, 132} |
|  |  |  |  | Set-27 | {23, 32, 52, 91, 139} |
|  |  |  |  | Set-28 | {30, 115, 66, 108, 116, 117} |
|  |  |  |  | Set-29 | {38, 123, 131, 135} |
|  |  |  |  | Set-30 | {93, 136, 137} |
|  |  |  |  | Set-31 | {31, 80} |
|  |  |  |  | Set-32 | {81, 82} |

**Figure 7.**
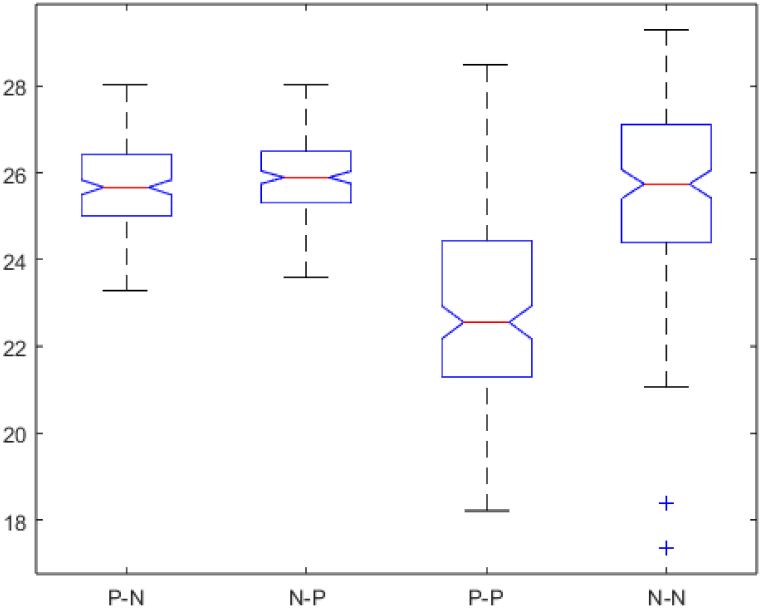
Box-plot of the relative frequency of PN, NP, PP, and NN changes in GAPDH moonlighting proteins

**Figure 8.**
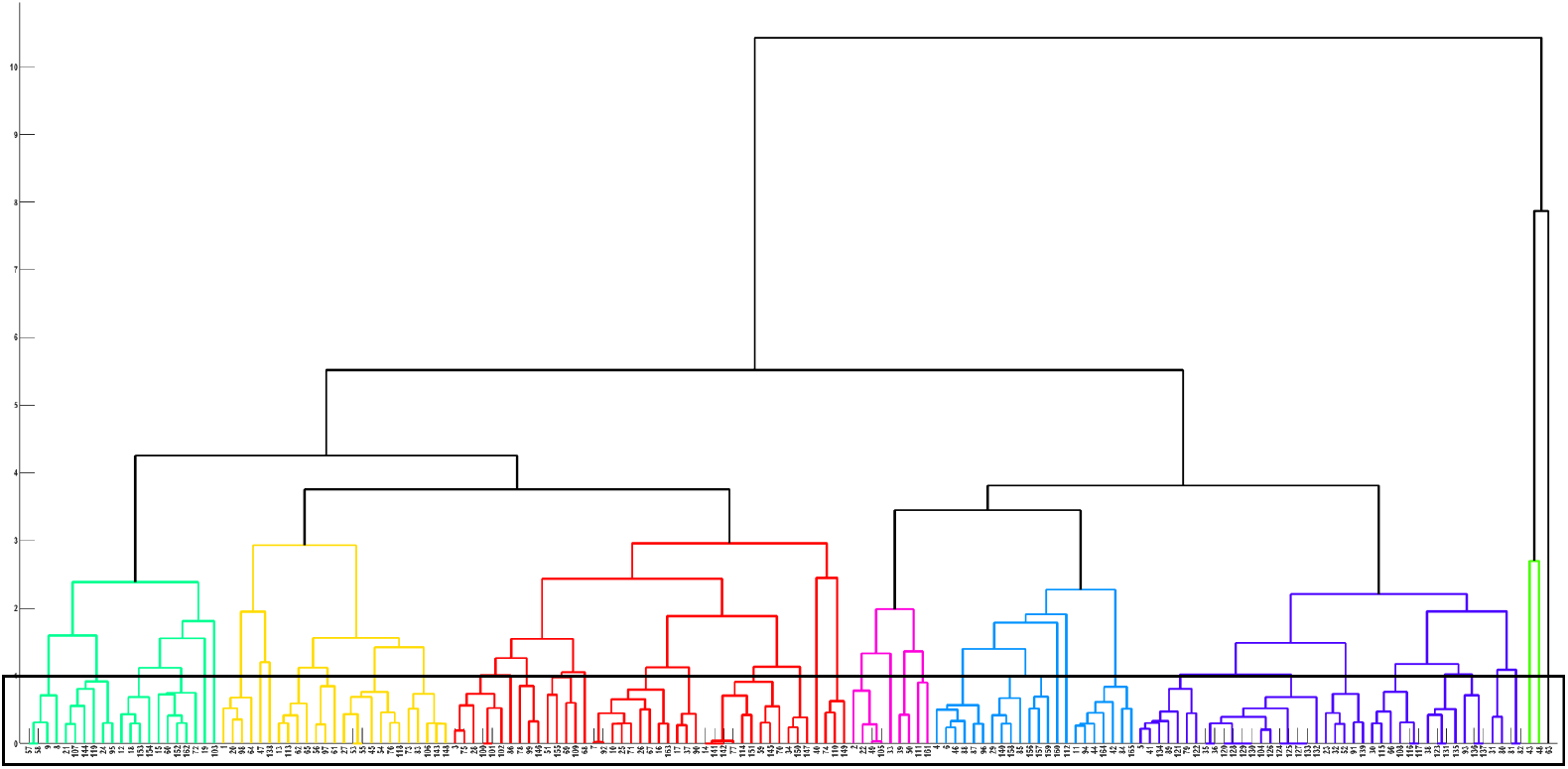
Phylogenetic relationship among the GAPDH moonlighting proteins based on relative frequency of PP, NP, PP, and NN changes as obtained from polar, non-polar profiles.

Domain-wise classification of these sets revealed a diverse taxonomic composition. All the proximal sets as demonstrated in Table 12, except Set-3, 19, 23 and 32 were composed of GAPDH sequences spanning more than one domain, including combinations of Archaea, Bacteria, and Eukarya, reflecting extensive evolutionary overlap. Specifically, members of the Set-6 and Set-8 belonged to Archaea and Bacteria, while members of the Set-14, Set-15, Set-16, Set-17, Set-25, and Set-29 belonged to Archaea and Eukarya; and members of the Set-20, Set-22, Set-26, Set-28, Set-30, and Set-31 belonged to Eukarya and Bacteria. In contrast, Set-3 consisted solely of Eukaryotic GAPDH sequences. Additionally, Set-23 and Set-32 consisted exclusively of Bacterial GAPDH sequences, while the Set-19 was composed solely of Archaeal GAPDH members.

#### 4.5.3. Phylogenetic relationships among GAPDH sequences based on polar, non-polar walk

Polar, non-polar walk of 165 GAPDH sequences were derived as the way discussed in the method **section**. P-NP walk of only two sequences (P47543 and P58839) are shown here as example (Figure 9). While calculating Hausdorff distance between this pair of sequences, both the walks were scaled w.r.t P47543 as P47543 is of shorter length compared to P58839 (Figure 9). Hausdorff distances between P-NP walks of all possible pair of sequences were enumerated in similar way.

Eight clusters were formed based on a distance threshold of 10 (Figure 10), of which the two larger clusters contained 68 and 65 GAPDH sequences (marked red in red and green in dendrogram Figure 10). While setting the distance threshold to 2 (20% of the original threshold), 11 disjoint sets were emerged containing 42 GAPDH sequences (Table 13).

**Table 13.** List of sets consist of proximal GAPDH protein sequences based on P-NP walk.

| Sets based on the proximity derived from P-NP walk |  |  |  |
| --- | --- | --- | --- |
| **Set-1** | {99, 100, 101, 102} | **Set-7** | {67, 71, 70} |
| **Set-2** | {150, 151} | **Set-8** | {35, 36, 134, 124, 125, 127, 133, 132, 128, 129, 130} |
| **Set-3** | {17, 95} | **Set-9** | {135, 136, 137} |
| **Set-4** | {14, 141, 142, 143} | **Set-10** | {80, 81, 82} |
| **Set-5** | {45, 53, 55, 54} | **Set-11** | {156, 157, 158, 159} |
| **Set-6** | {153, 154} |  |  |

**Figure 9.**
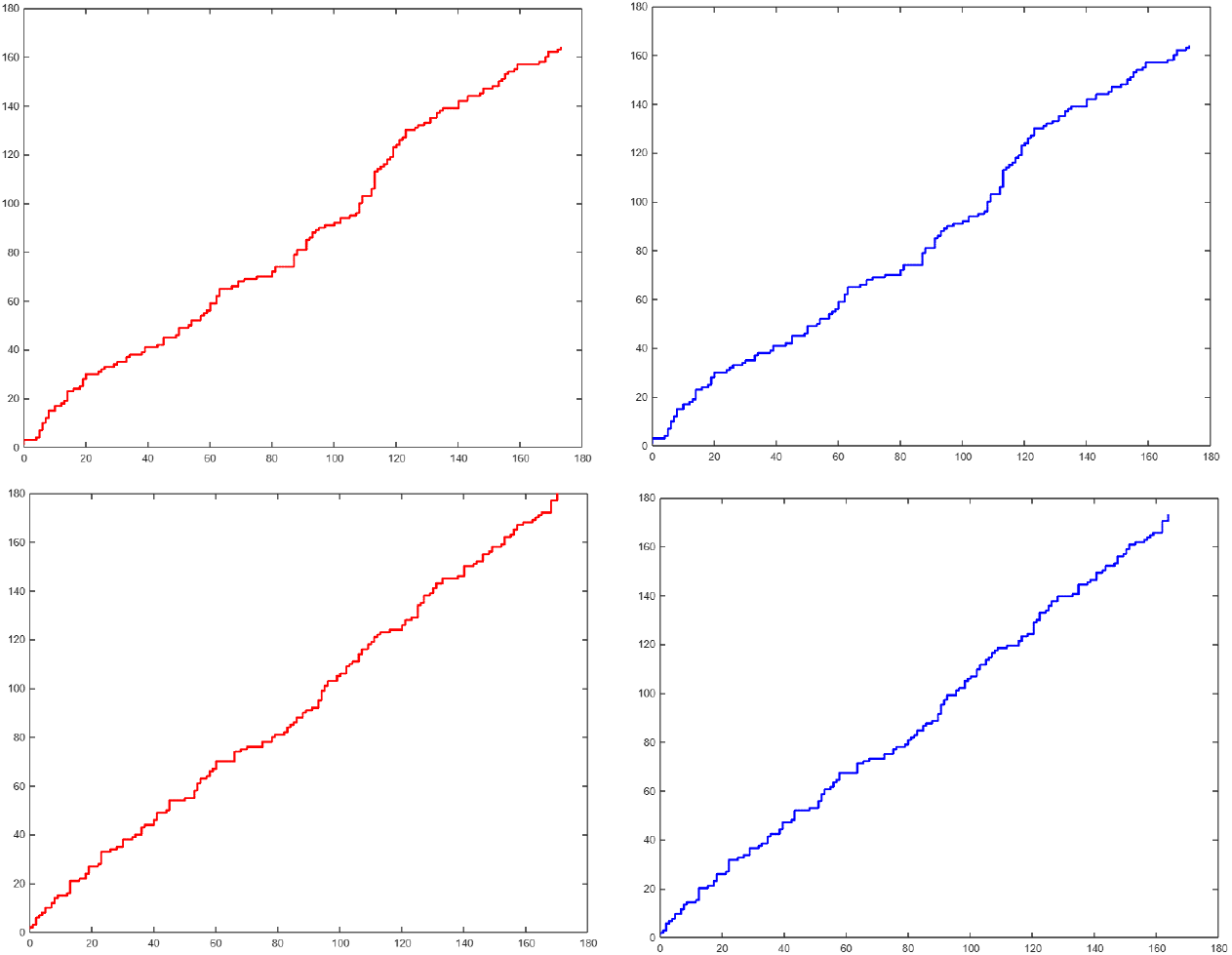
P-NP walks of P47543 (top left) and P58839 (bottom left) and their respective scaled P-NP walks (top right and bottom right)

**Figure 10.**
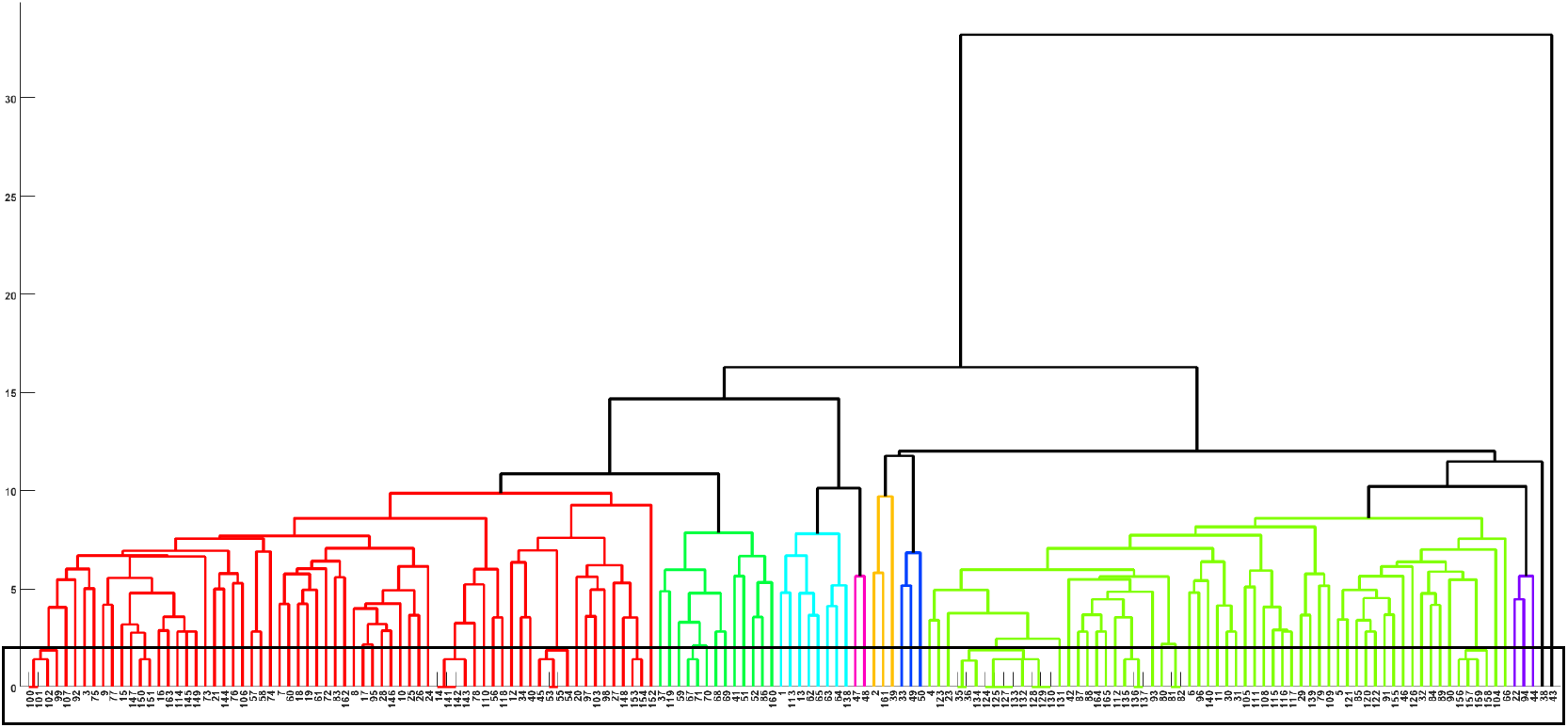
Phylogenetic relationships among the GAPDH moonlighting proteins based on P-NP walk.

Domain-wise and taxonomic classification of the 11 proximal sets derived from the P-NP walk-based clustering (Table 13) revealed distinct evolutionary groupings. The proximal Set-2, Set-3, and Set-6 consisted exclusively of the sequences from the domain Eukaryota, specifically belonging to the kingdom Fungi. The Set-4, Set-5, and Set-7 were composed entirely of Archaeal sequences. The Set-8 included only Eukaryotic members, all classified under phylum Chordata, class Mammalia, while the Set-9 was also strictly Eukaryotic, represented by phylum Chordata, class Aves. On the other hand, Set-10 and Set-11 consisted of GAPDH sequences exclusively from the domain Bacteria, highlighting conserved polar, non-polar profiles within this group.

### 4.6. Intrinsic protein disorder analysis of GAPDH moonlighting proteins

The proportions of disordered, highly flexible, moderately flexible, and other residues in each GAPDH moonlighting protein were calculated (Figure 11). Three GAPDH moonlighting proteins showed notably high percentages of disordered residues: Q5V061 (38.98%), Q8J1H3 (32.34%), and Q1DTF9 (31.16%). In contrast, proteins with significantly low percentages of disordered residues included P17721 (3.90%), P00361 (5.74%). The highest proportions of highly flexible residues were observed in P23722 (70.45%) followed by B0R2M2 (68.95%), while the lowest was found in O13507 (35.50%). For moderately flexible residues, Q59199 exhibited the highest percentage (45.35%) followed by P46713 (44.54%), whereas B0R2M2 had the lowest (9.55%). Notably, eighteen GAPDH moonlighting sequences showed zero residues in the “other” category: P23722, P00362, Q59199, P39460, Q97BJ8, Q9HJ69, O27090, Q2NEN0, A5ULT9, P19315, P19314, A0QWW2, Q59309, P9WN83, P0C0G7, P0C0G6, P0DB18, and Q59906 (Figure 11).

**Figure 11.**
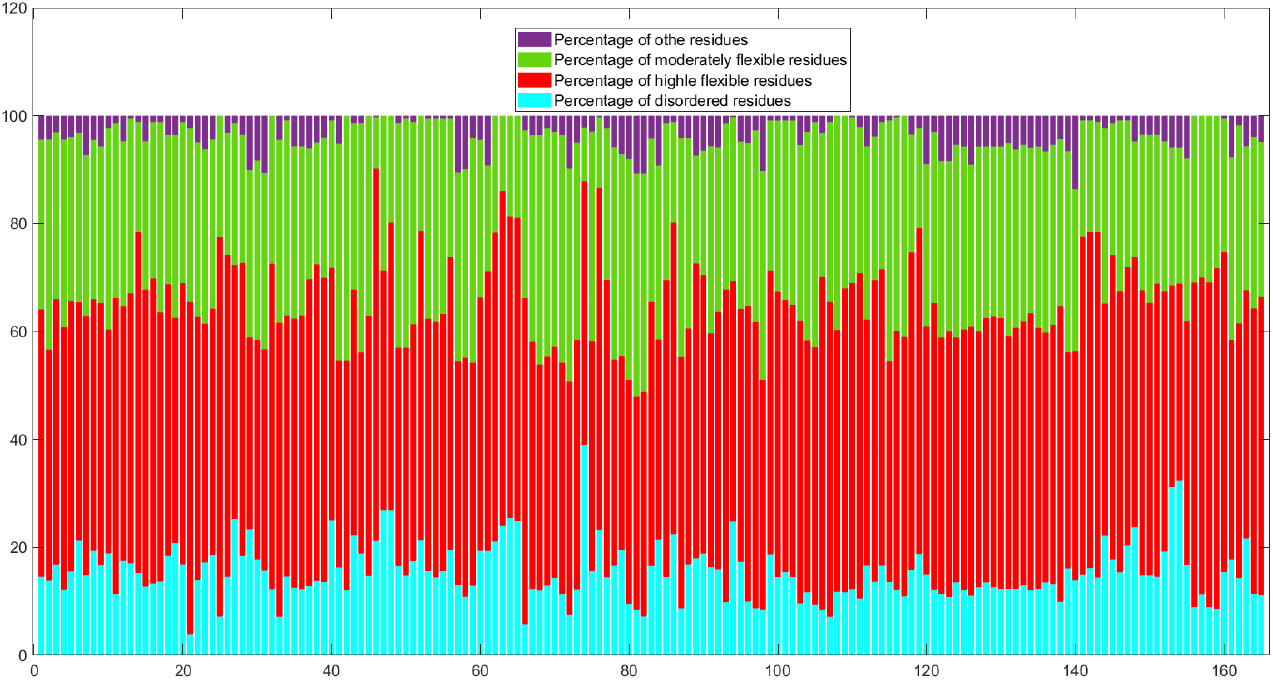
Percentages of disordered, highly flexible, moderately flexible and other residues in GAPDH moonlighting proteins

#### 4.6.1. Percentage of four different IPD residue types in each amino acid

Each of the four figures in Figures 12 and 13) corresponds to one residue type (disordered (D), highly flexible (HF), moderately flexible (MF), and other (O) residues) and displays the percentage of that residue in each amino acid. It needs to be pointed out that percentages were calculated based on individual amino acid frequency in a sequence implying the sum of the percentages of four different IPD residue types will be 100 for a given amino acid in a sequence.

**Figure 12.**
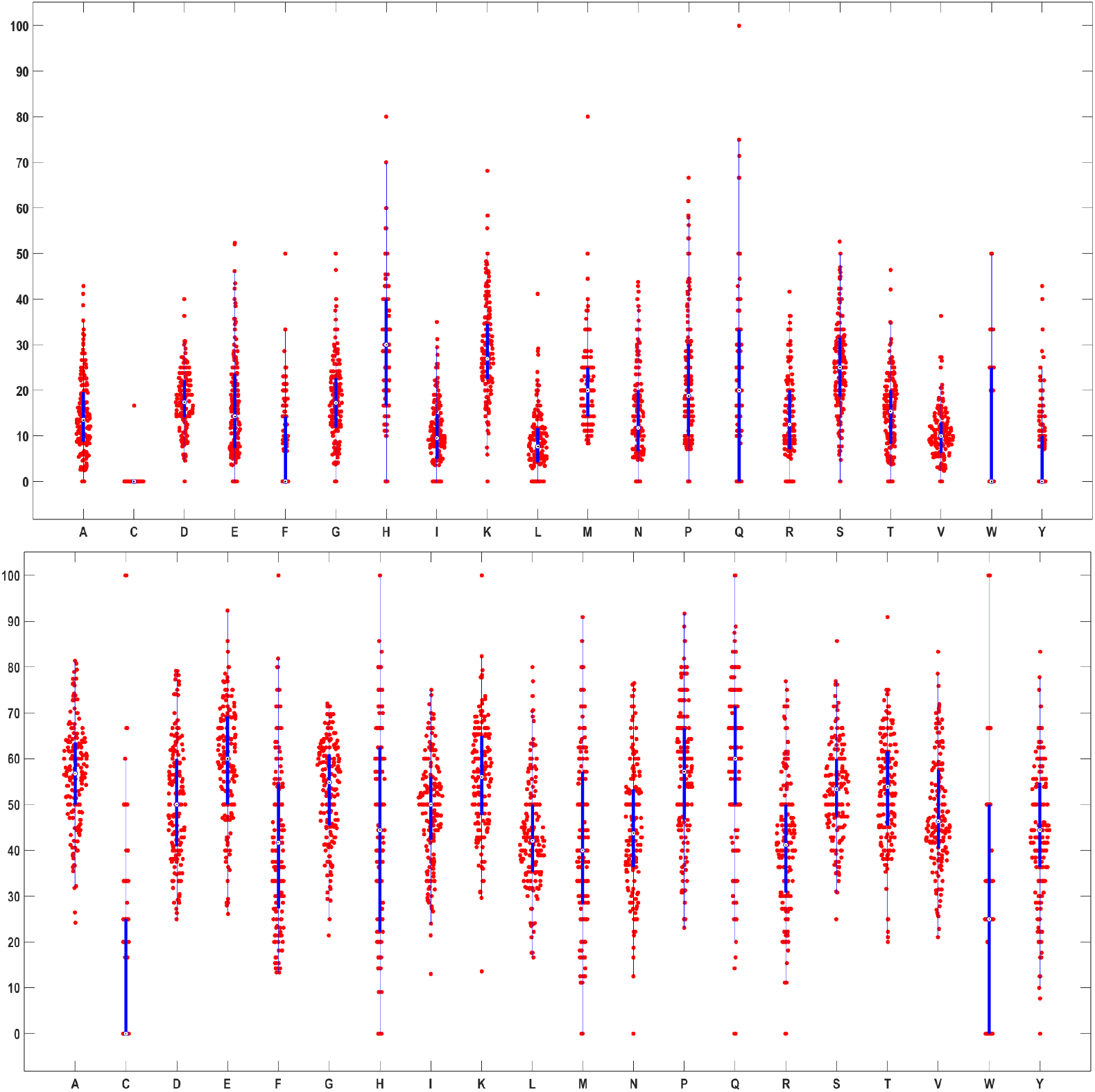
Percentage of disordered (Top) and highly flexible (Bottom) regions in each amino acid in 165 GAPDH sequences

**Figure 13.**
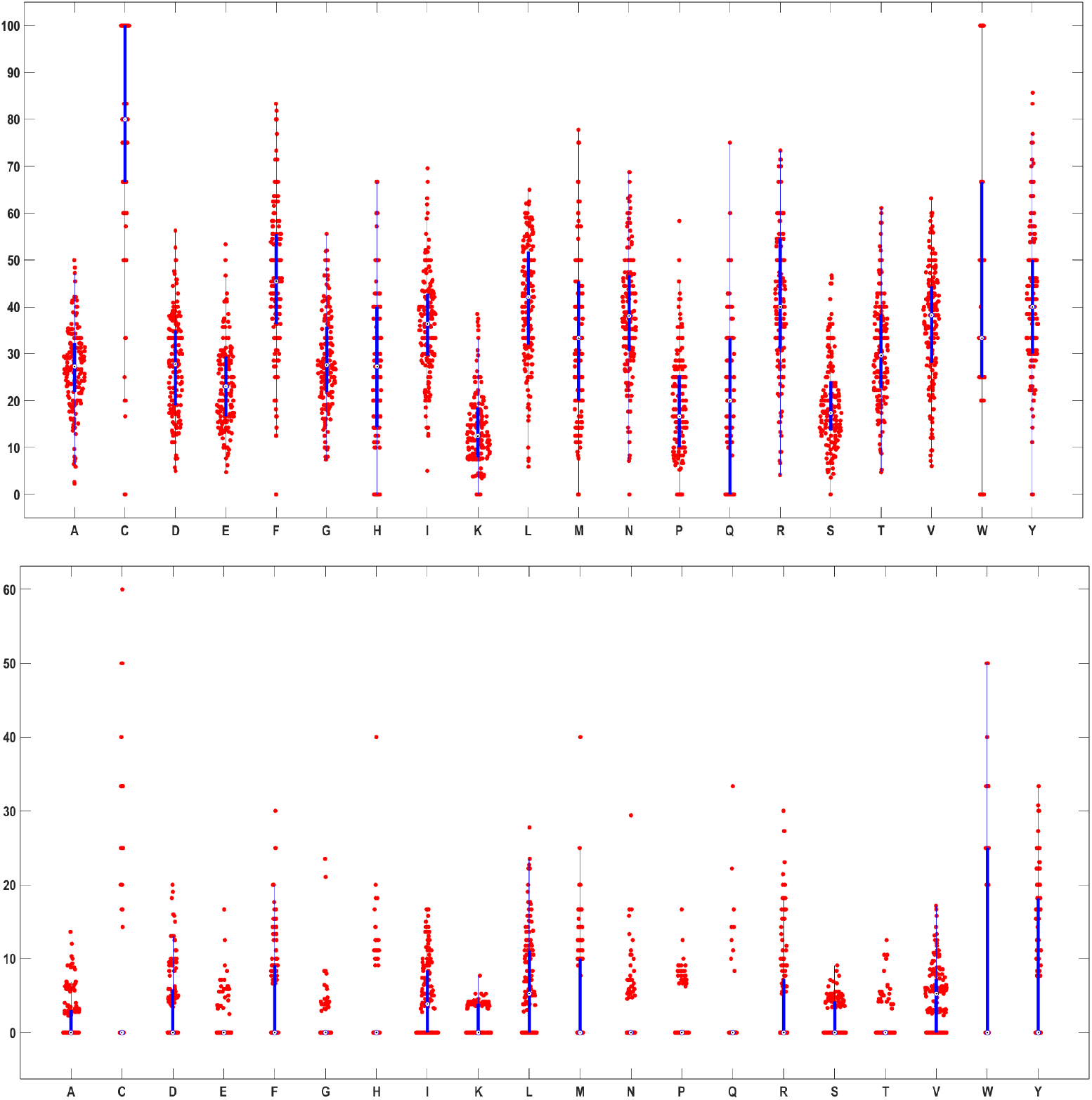
Percentage of moderately flexible (Top) and other regions (Bottom) in each amino acid in 165 GAPDH sequences

The only sequence with disordered cysteine residues is P34783, which contains 16.667% disordered residues (denoted as ‘D’) specifically in cysteine (C). No other GAPDH sequence shows disordered cysteine residues. Additionally, all glutamine (Q) present in the sequence Q00584 were found to be disordered. Among the 165 GAPDH protein sequences analyzed, five amino acids—alanine (A), cysteine (C), phenylalanine (F), tryptophan (W), and tyrosine (Y)—were found to have no disordered residues in sequences P61880 and O59494. In sequence P17721, no disordered residues were present in the amino acids C, D, F, H, I, N, Q, S, T, W, and Y. Thirteen sequences showed no disordered residues in glutamic acid (E), although glutamic acid displayed a high disorder content of approximately 52% in sequences Q97BJ8 and Q9YFS9. Similarly, 82 sequences contained no disordered phenylalanine residues, whereas sequence Q6L125 had 50% of its phenylalanine residues disordered. No disordered glycine (G) residues were found across four sequences—Q9Z7T0, Q07234, Q89AK1, and Q01651. Histidine (H) showed no disorder in 13 GAPDH sequences, while it exhibited the highest disorder percentage (80%) in Q9UVC0. Twenty-seven sequences contained no disordered isoleucine (I) residues. Disordered leucine (L) residues were absent in 32 sequences, while sequences Q1DTF9 and Q8J1H3 had 41.18% of their leucine residues disordered. No disordered methionine (M) was observed in Q6M0E6 and Q971K2, but Q5V061 had 80% of its methionine residues disordered. Thirteen sequences lacked disordered asparagine (N), while Q8TFJ2 contained 43.75% disordered asparagine. In five sequences—P23722, P15115, Q9ZKT0, Q07234, and P80534—proline (P) residues were present but not disordered, whereas Q9HJ69 showed 66.67% disordered proline. Glutamine (Q) showed no disorder in 45 sequences; however, Q9HGY7, Q1DTF9, and Q8J1H3 had 75% of their glutamine residues disordered. Thirty-three sequences showed no disordered arginine (R). No disordered serine (S) residues were found in P17721, P15115, and Q9Z518, whereas A5ULT9 and Q5V061 exhibited 52.63% disordered serine. Threonine (T) showed no disorder in 13 sequences, but Q5V061 had 46.43% disordered threonine residues. Valine (V) showed no disordered residues in O52631, Q89AK1, and Q9Z518. Tryptophan (W) had no disorder in 92 sequences, yet 50% of the tryptophan residues were disordered in Q9YFS9 and Q8NK47. Tyrosine (Y) showed no disorder in 91 sequences, while Q9YFS9 had 42.857% disordered tyrosine residues (**Supplementary file-2**).

The amino acids A, D, E, F, G, I, K, L, P, S, T, and V each exhibited a minimum presence ranging from 13.04% to 26.09% and a maximum presence of up to 100% among the ‘HF’ residues across 165 GAPDH sequences. All cysteine residues in P00361, Q48335, P87197, Q59800, and P52987 were found to be highly flexible, whereas no ‘HF’ cysteine residues were identified in 95 GAPDH sequences. Furthermore, it was noted that 50% of the cysteine residues in P47543, P0A038, P46795, A5ULT9, P75358, Q5V061, Q12XM4, Q9UW96, and P28844 were classified as ‘HF’. It was observed that 79.17% of the aspartic acid (D) residues in P23722 and Q01651 were classified as ‘HF’. Additionally, nineteen sequences were identified in which no histidine residues were marked as ‘HF’, whereas all histidine residues in P87197 and Q89AK1 were classified as ‘HF’. None of the methionine (M) residues were classified as ‘HF’ in the following nine sequences: O67161, Q9UR38, Q01982, Q01651, Q59800, Q8NK47, P9WN83, Q9UW96, and Q92243. In contrast, 90.91% of the methionine residues in A7IB57 were identified as ‘HF’. More than 75% of the asparagine (N) residues in each of the GAPDH sequences– P00362, P15115, Q07234, and Q59906 were classified as ‘HF’, whereas none of the asparagine residues in A2BLC6 were identified as ‘HF’. None of the glutamine (Q) residues were classified as ‘HF’ in Q6CCU7, Q00584, Q01982, Q9UW96, and Q8WZN0, whereas all glutamine residues in P17721, P20287, P80534, and Q94469 were identified as ‘HF’.None of the arginine (R) residues were classified as ‘HF’ in Q92263 and P08439, while 76.92% of the arginine residues in P19314 were identified as ‘HF’. All tryptophan (W) residues in A7IB57, A3CYG8, Q2FNA2, A3MTU1, Q8ZWK7, A4WIW2, B1YD06, and Q59309 were identified as ‘HF’, whereas no tryptophan residues were classified as ‘HF’ in 53 of the 165 GAPDH sequences. A total of 83.33% of the tyrosine (Y) residues in Q59800 were classified as ‘HF’, while none of the tyrosine residues in Q9Z518 were identified as ‘HF’ (**Supplementary file-2**).

The amino acids A, D, E, G, I, L, R, T, and V each exhibited a minimum presence ranging from 2.33% to 7.71% among the moderately flexible (‘MF’) residues across 165 GAPDH sequences. Additionally, 77 GAPDH sequences were identified in which all cysteine residues were found to be ‘MF’. Similarly, 31 sequences were identified where all tryptophan (W) residues were classified as ‘MF’. None of the cysteine residues in P00361, Q5V061, Q48335, P87197, Q59800, and P52987 were classified as ‘MF’, whereas 77 GAPDH sequences were identified in which all cysteine residues were marked as ‘MF’. In Q6M0E6 and A5ULT9, none of the phenylalanine (F) residues were classified as ‘MF’, whereas 83.33% of the F residues in P44304 were identified as ‘MF’. None of the histidine residues in 30 GAPDH sequences were marked as ‘MF’, whereas 66.67% of the histidine residues in P39460, C3MWN0, C3MQN0, C3N6F0, P61880, and O59494 were classified as ‘MF’. No ‘MF’ residues were identified among the lysine (K) residues in P58839, B0R2M2, Q9HJ69, A5ULT9, Q3IUT3, and Q8PTD3, whereas the highest percentage—38.46%—of lysine residues classified as ‘MF’ was observed in Q8ZWK7. None of the methionine residues in Q9UVC0, A7IB57, Q5V061, Q3IUT3, and Q9HFX1 were classified as ‘MF’, whereas 77.78% of the methionine residues in A4WIW2 were identified as ‘MF’. None of the asparagine (N) residues in Q5V061 were classified as ‘MF’, whereas 68.75% of the N residues in Q9N655 and P0CN74 were identified as ‘MF’. None of the proline residues were classified as ‘MF’ in 14 GAPDH sequences, whereas 58.33% of the proline residues in Q92263 were marked as ‘MF’. In 53% of the GAPDH sequences, no glutamine (Q) residues were classified as ‘MF’. A total of 46.67% of the serine residues in GAPDH sequences P17721 and B6YUU0 were identified as ‘MF’, while among all sequences, only Q48335 contained no serine residues classified as ‘MF’. Tryptophan had not been marked as ‘MF’ residues in 30 GAPDH sequences. Amino acid Y in P00362, Q27652, and Q59800 was not identified as ‘MF’, while 85.71% of Y present in Q9Z518 were all marked as ‘MF’ (**Supplementary file-2**).

Among the 165 GAPDH sequences, 31 were identified in which no amino acid residues were classified as ‘Other’ (‘O’). Only the amino acids A, D, G, and F were marked as ‘O’ in the sequences O52631, Q9Z518, A3MTU1, and P46713, respectively, with corresponding percentages of 2.56%, 3.45%, 3.70%, and 11.11%. The highest proportion of ‘O’ residues was observed in cysteine residues in sequence P55971. Additionally, 50% of the tryptophan (W) residues in Q9UVC0, A2BLC6, P09316, Q6L125, Q5V061, Q58546, and P51009 were classified as ‘O’. The amino acid glutamine (Q) was not marked as ‘O’ in 156 GAPDH sequences; however, in one sequence, Q01651, Q was identified as ‘O’ (**Supplementary file-2**).

#### 4.6.2. Change response sequences of disordered, highly flexible, moderately flexible, and other residues profiles

Figure 14 illustrates the distribution of sixteen transition types derived from intrinsic disorder profiles. The analysis, visualized through boxplots, highlights distinct patterns of structural flexibility that differentiate moonlighting proteins from their non-moonlighting counterparts. No transitions from ‘other’ to ‘highly flexible’ (O_HF), O_D, MF_D, HF_O, D_O, or D_MF were observed across all 165 intrinsic protein disorder (IPD) profiles of GAPDH sequences. The protein Q92243 exhibited the highest percentage (12.8%) of O_O transitions. Identical percentages of transitions in O_MF and MF_O, were recorded across all 165 GAPDH sequences. The highest percentage of MF_MF transitions (8.215%) was observed in Q59199 and it had highest percentage of moderately flexible residues. Significantly low percentage of MF_MF transitions were found in B0R2M2 (8.38%), Q5V061 (8.21%) and Q48335 (10.47%). The protein P0CE13 showed the highest percentage (4.505%) of MF_HF and HF_MF transitions. Additionally, identical percentages of MF_HF and HF_MF transitions were consistently observed across all 165 sequences. Among all 16 transition types, HF_HF transitions were found to be the most prevalent across the 165 GAPDH sequences. Specifically, the lowest (31.55%) and highest (66.77%) HF_HF transition percentages were found in Q01982 and P23722, respectively (Figure 14). Percentages of HF_D and D_HF transitions were identical across all sequences. Sequences P17721 and Q5V061 displayed the lowest (2.41%) and highest (37.68%) percentages of D_D transitions, respectively as they had lowest and highest percentage of disordered residues.

**Figure 14.**
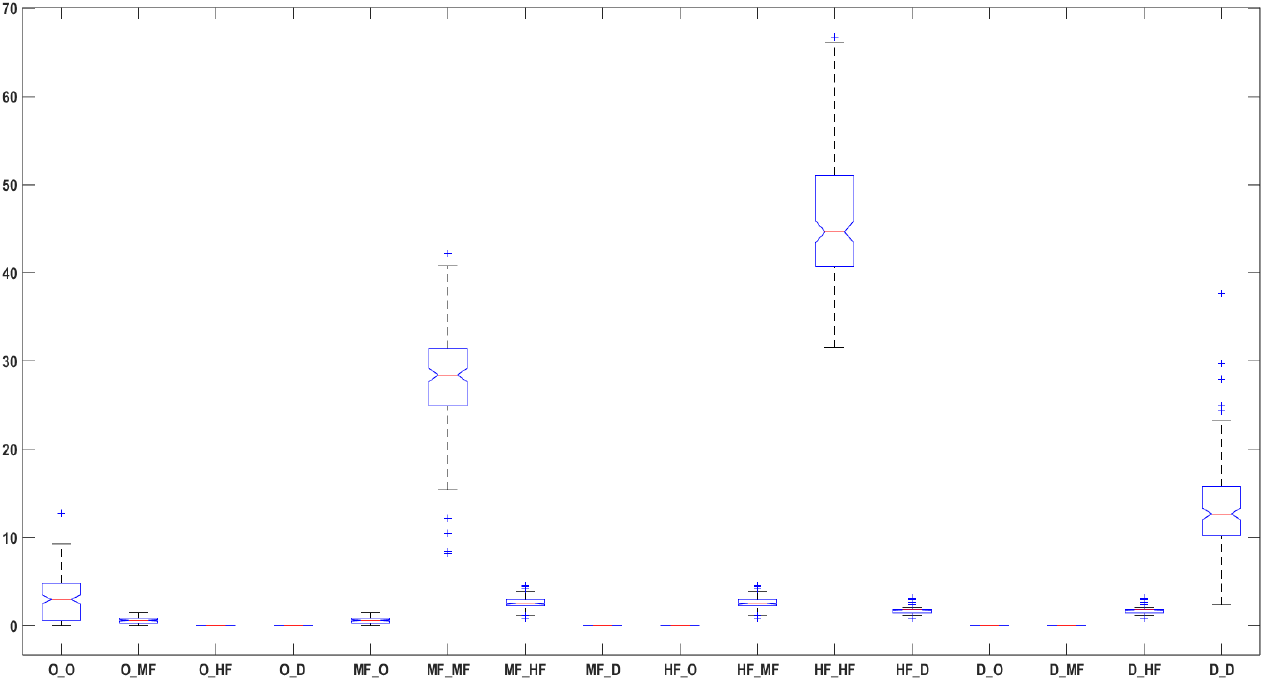
Box-plot of the percentages of disordered, highly flexible, moderately flexible and other residues changes in GAPDH moonlighting protein sequences

An analysis of 165 GAPDH moonlighting sequences, using a distance threshold of 16, resulted in the formation of seven distinct clusters, which can be observed in the dendrogram (Figure 15. The most substantial cluster consisted of 69 GAPDH sequences (marked in green (Figure 15)). Clusters range from small (e.g., rightmost cluster in blue with only 3 GAPDH sequences) to large (e.g., Cluster marked in green with 69 sequences), suggesting that some proteins share common flexibility characteristics while others are more specialized.

**Figure 15.**
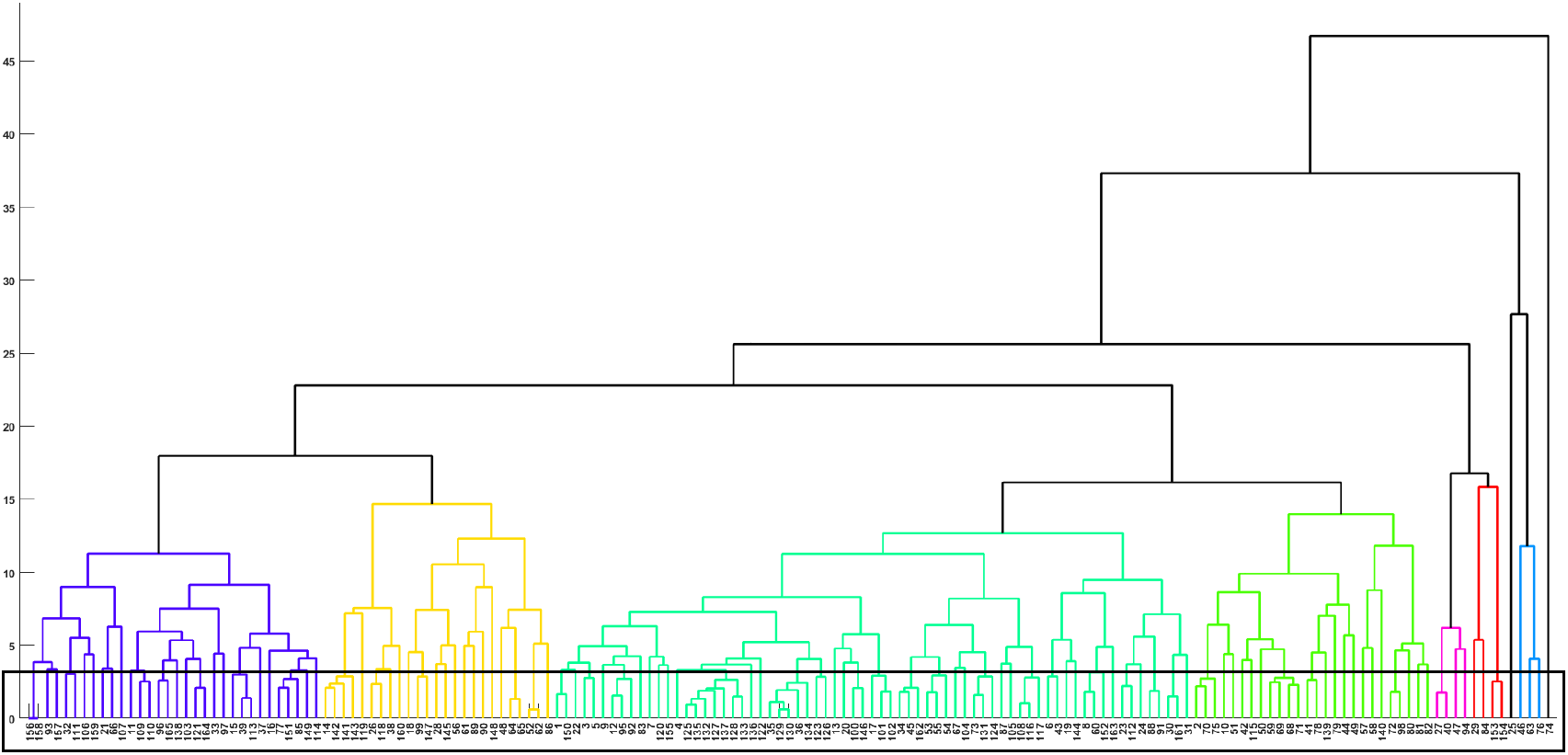
Phylogenetic relationship among the GAPDH moonlighting proteins based on change response sequences of disordered, highly flexible, moderately flexible, and other residues profiles.

Upon thresholding the distance threshold to 3.2, 105 GAPDH sequences were clubbed into 36 disjoint sets (Table 14). Domain-wise and taxonomic analysis of the 36 proximal sets derived from IPD profile revealed widespread diversity across evolutionary lineages (Table 14). Set-1, Set-2, Set-3, Set-25, and Set-34 consisted exclusively of Bacterial GAPDH sequences. Six sets (Set-9, Set-12, Set-13, Set-23, Set-32, and Set-33) were solely composed of Archaeal GAPDH sequences. Set-4, Set-10, and Set-35 included both Bacterial and Archaeal members. The Set-5 comprised GAPDH sequences from Bacteria and Eukarya, specifically from Arthropoda (Malacostraca). Set-6 contained Eukaryotic sequences, spanning two distinct kingdoms—Fungi and Animalia (Malacostraca). Broad domain diversity was observed in Set-7 and Set-14, which included representatives from Bacteria, Archaea, and Eukaryota. Notably, Set-8 featured exclusively Eukaryotic GAPDH sequences from Fungi and Mold. Several proximal sets showed cross-domain representations involving Eukarya and Bacteria: Set-11, Set-20, Set-21, and Set-27 included GAPDH sequences from Bacteria and Eukaryotic fungi organisms, while Set-24 combined Bacterial and Mammalian sequences. Other sets demonstrated specific cross-domain distributions, such as Set-15 and Set-26, which included both Archaea and Eukaryotic fungi, and Set-22 and Set-31, comprising Archaea and Eukarya (Amoebozoa). Several sets were strictly Eukaryotic: Set-16 included Fungi, Plants, and Animals (Platyhelminthes); Set-17 contained Mammals and Aves; Set-18 included only Mammals; and Set-19 included both Actinopterygii and Mammals. Likewise, Set-28 and Set-36 were limited to Eukaryotic fungi, Set-29 included Insects and Fungi, and Set-30 contained only Fungal entries.

**Table 14.** List of sets consist of proximal GAPDH protein sequences based on frequency of changes between adjacent residues of intrinsic protein disorder profiles.

| Sets of proximal GAPDH sequences |  |  |  |
| --- | --- | --- | --- |
| **Set-1** | {156, 158} | **Set-13** | {52, 62} |
| **Set-2** | {93, 157} | **Set-14** | {1, 150, 22} |
| **Set-3** | {32, 111} | **Set-15** | {3,5} |
| **Set-4** | {11, 109, 110} | **Set-16** | {12, 95, 92} |
| **Set-5** | {96, 165} | **Set-17** | {125,127, 128, 132, 133, 135, 137} |
| **Set-6** | {121, 164} | **Set-18** | {35, 36, 129, 130, 134} |
| **Set-7** | {15, 39, 113} | **Set-19** | {123, 126} |
| **Set-8** | {77, 151, 85, 149} | **Set-20** | {100, 146} |
| **Set-9** | {14, 141, 142, 143} | **Set-21** | {17, 101, 102} |
| **Set-10** | {26, 118, 38} | **Set-22** | {34, 45, 162} |
| **Set-11** | {99, 147} | **Set-23** | {53, 54, 55} |
| **Set-12** | {64, 65} | **Set-24** | {73, 124, 131} |
| **Set-25** | {108, 116, 117} | **Set-26** | {6, 43} |
| **Set-27** | {8, 60, 152} | **Set-28** | {23, 112} |
| **Set-29** | {88, 91} | **Set-30** | {30, 161} |
| **Set-31** | {2, 70, 75} | **Set-32** | {59, 68, 69, 71} |
| **Set-33** | {41, 78} | **Set-34** | {72,98} |
| **Set-35** | {27, 40} | **Set-36** | {153, 154} |

#### 4.6.3. Phylogenetic relationships among GAPDH sequences based on IPD walk

Intrinsic protein disorder walk (IPD walk) of 165 GAPDH sequences were derived as the way discussed in the method **section**. IPD walk of only two sequences (P47543 and P58839) were presented as example in Figure 16). While calculating Hausdorff distance between this pair of sequences, both the IPD walks were scaled w.r.t P47543 as P47543 is of shorter length compared to P58839 (Figure 16). Hausdorff distances between IPD walks of all possible pair of sequences were enumerated in similar way.

**Figure 16.**
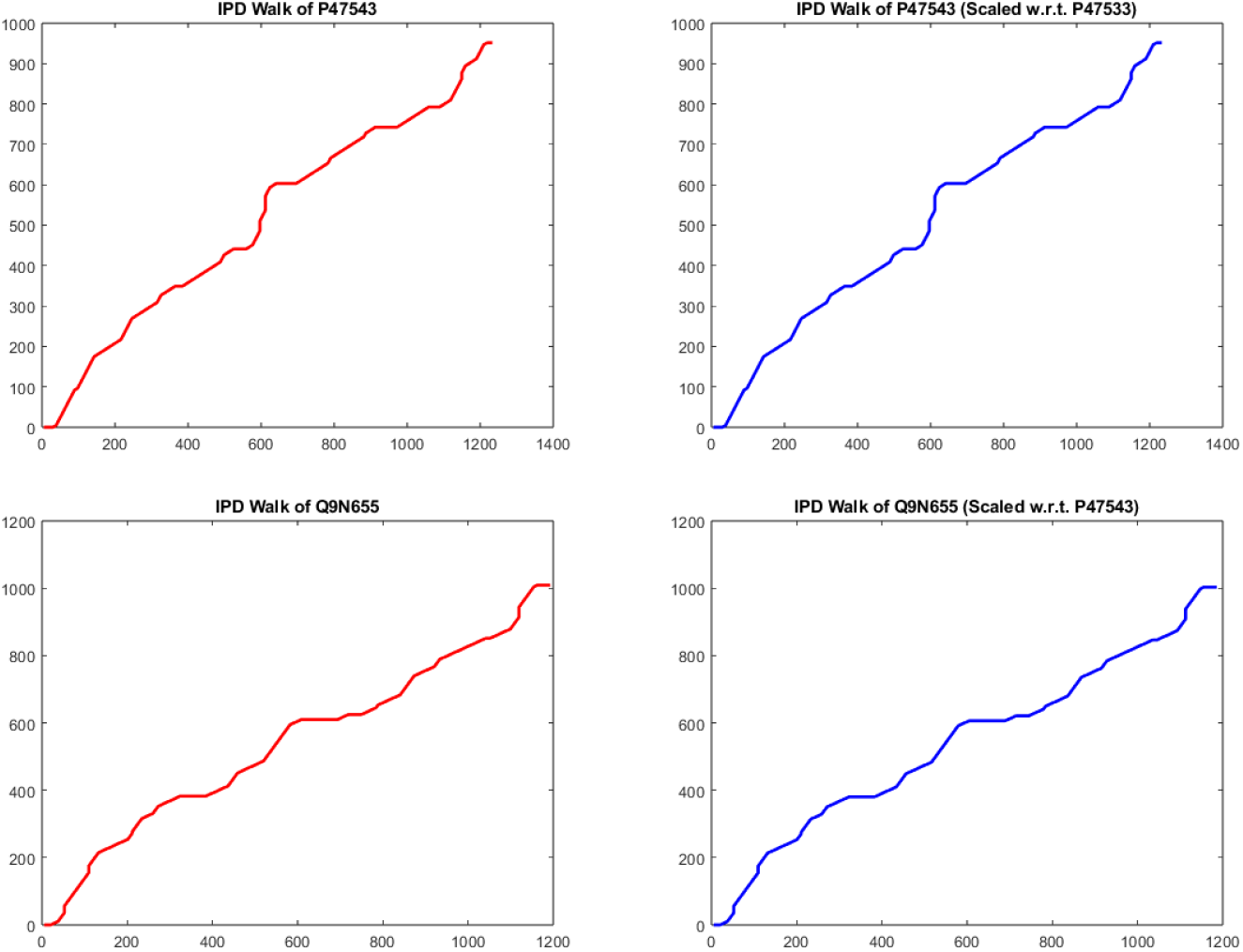
IPD Walk of P47543 (top left) and P58839 (bottom left) and their respective scaled IPD walks (top right and bottom right)

An analysis of IPD profiles of 165 GAPDH moonlighting sequences, using a distance threshold of 100, resulted in the formation of ten distinct clusters, which can be observed in the dendrogram (Figure 17). The most substantial cluster consisted of 62 GAPDH sequences and was highlighted in red, while the smallest cluster contained 2 GAPDH sequences (Figure 17). While setting the distance threshold to 20 (20% of the original threshold), 17 disjoint sets were emerged containing 42 GAPDH sequences (Table 15). Taxonomic characterization of the 17 disjoint sets generated from intrinsic protein disorder (IPD) walk-based clustering (Table 15) revealed domain-specific and lineage-specific clustering patterns. Set-1, Set-2, Set-6, and Set-16 were composed exclusively of sequences from the domain Bacteria, suggesting conserved disordered structural patterns within prokaryotic GAPDH homologs. Set-3, Set-4, and Set-17 contained only Eukaryotic sequences, specifically belonging to the kingdom Fungi, highlighting lineage-specific flexibility profiles among fungal proteins. In contrast, Set-5, Set-13, Set-14, and Set-15 were formed solely by Archaeal sequences, underscoring distinct patterns of intrinsic disorder within extremophilic organisms. Among the remaining sets, Set-7, Set-9, and Set-12 were composed of Eukaryotic sequences restricted to the class Mammalia, reflecting conserved disorder architecture within this vertebrate lineage. Set-8 and Set-10 also fell under the domain Eukaryota, but comprised a mix of sequences from both Mammals and Aves, indicating shared disordered features across endothermic vertebrates. Notably, Set-11 included sequences from Eukaryotic Arthropods, specifically within the class Malacostraca, suggesting that disorder-driven profiles in invertebrates can also form cohesive phylogenetic clusters.

**Table 15.** List of sets consist of proximal GAPDH protein sequences based on IPD walk.

| Sets of proximal GAPDH sequences based on IPD walk |  |  |  |
| --- | --- | --- | --- |
| **Set-1** | {156, 158} | **Set-9** | {124, 125, 127, 131} |
| **Set-2** | {100, 101, 102} | **Set-10** | {132, 135, 136} |
| **Set-3** | {149, 151} | **Set-11** | {164, 165} |
| **Set-4** | {5, 95} | **Set-12** | {128, 129, 130} |
| **Set-5** | {14, 142, 141, 143} | **Set-13** | {53, 55, 54} |
| **Set-6** | {108, 116} | **Set-14** | {59, 71} |
| **Set-7** | {35, 36} | **Set-15** | {67, 70} |
| **Set-8** | {134, 137} | **Set-16** | {81, 82} |
|  |  | **Set-17** | {153, 154} |

**Figure 17.**
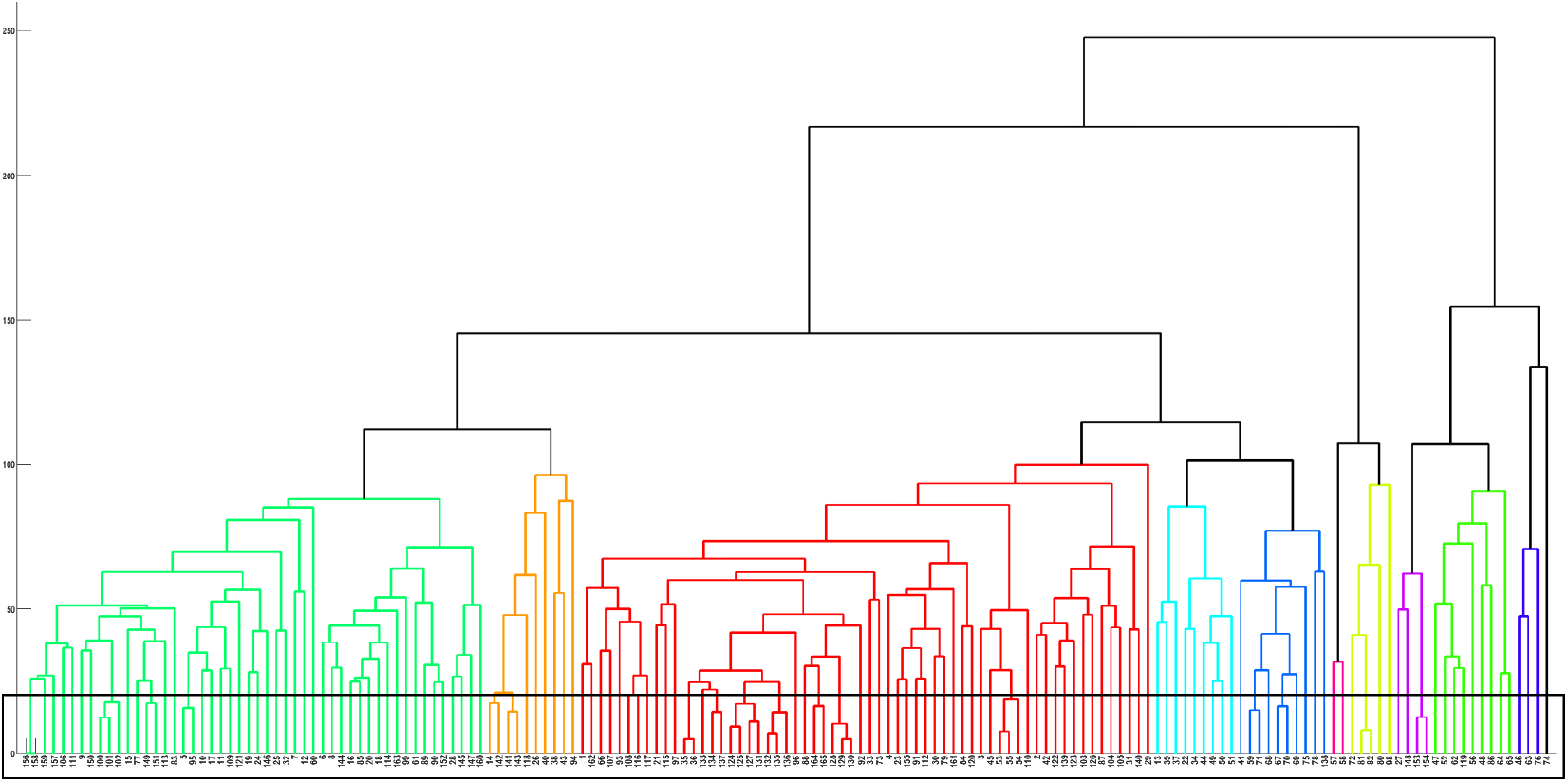
Phylogenetic relationship among the GAPDH moonlighting proteins based on IPD walk.

### 4.7. Derivatives from indicator matrices formed by GAPDH sequences

Indicator matrices for all 165 GAPDH sequences based on spatial amino acid compositions were calculated and accordingly binary images were generated. Such an example of indicator matrix for the GAPDH sequence–P10096 was determined (Figure 18).

**Figure 18.**
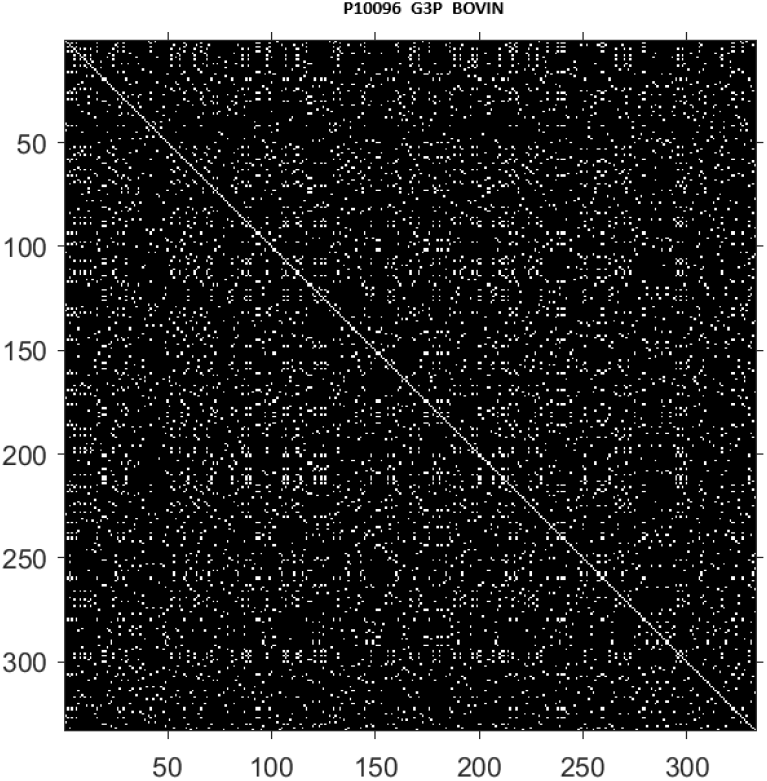
Indicator matrices for the GAPDH sequence–P10096

Fractal dimension, density of ones, frequency of connected components with connectivity 4 and 8 were determined for all indicator matrices (Table 16). A comparative scatter plot with histogram of these four derivatives from indicator matrices was presented in Figure 19. From the Table 16, it was observed that low variance in FD and density of ones indicates conserved complexity and sparsity patterns in indicator matrices. On the other side, the cluster counts (both 4- and 8-connectivity) show noticeable but bounded variability.

**Table 16.** Normalized statistical features derived from the indicator matrices of 165 GAPDH moonlighting sequences.

| Features and statistics | FD | Density of Ones | Number of clusters with 4 connectivity | Number of clusters with 8 connectivity |
| --- | --- | --- | --- | --- |
| *Maximum* | 1.443 | 0.077 | 8037.271 | 6467.347 |
| *Minimum* | 1.390 | 0.062 | 6364.384 | 5169.509 |
| *Average* | 1.411 | 0.067 | 6923.530 | 5654.358 |
| *Standard deviation* | 0.007 | 0.002 | 287.292 | 219.626 |
| *Median* | 1.411 | 0.066 | 6883.958 | 5622.707 |

**Figure 19.**
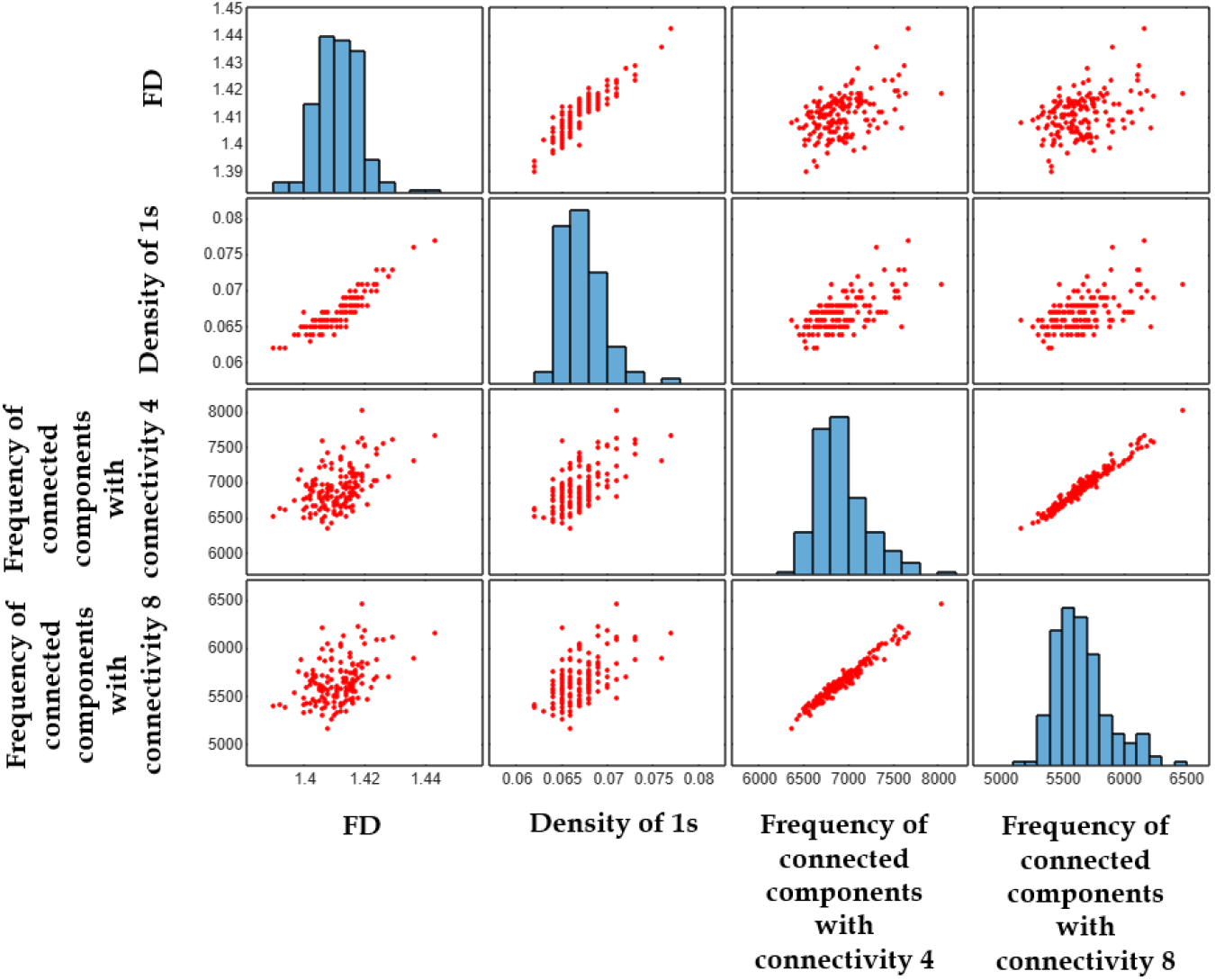
Fractal dimensions, density of ones, number of connected components with connectivity 4, and number of connected components with connectivity 4 in the indicator matrices derived from GAPDH sequences.

### 4.8. Proximal relationships among GAPDH moonlighting proteins

From the pool of 165 GAPDH proteins, a maximal intersecting family of proximal sets comprising 16 GAPDH moonlighting proteins was derived from Tables (3, 4, 12, 13, 14, and 15) as tabulated in Table 17.

**Table 17.** Maximal intersecting family of proximal sets of GAPDH moonlighting proteins based on various quantitative features.

| Family of proximal sets |  |  |  |
| --- | --- | --- | --- |
| *Proximal set-1* | {C3MWN0, C3MQN0, C3N6F0} all from *Sulfolobus islandicus* | *Proximal set-5* | P10096 (*Bos taurus*), Q9N2D5 (*Felis catus*) |
| *Proximal set-2* | {Q6M0E6, A6VIW4, A9A6M9} all from *Methanococcus maripaludis* | *Proximal set-6* | {Q1DTF9 (*Coccidioides immitis*), Q8J1H3 (*Coccidioides posadasii*)} |
| *Proximal set-3* | P0A9B3 (*Escherichia coli*), P0A1P0 (*Salmonella typhimurium*) | *Proximal set-7* | {Q5RAB4 (*Pongo abelii*), P04406 (*Homo sapiens*)} |
| *Proximal set-4* | Q4KYY3 (*Spermophilus citellus*), A3FKF7 (*Mustela putorius furo*) |  |  |

#### 4.8.1. Typical and atypical roles of GAPDH proteins within the proximal sets

The GAPDH proteins within each of the seven proximal sets were found to be involved in a range of both common and unique functions.

##### Canonical function

Sixteen GAPDH proteins belonging to seven proximal sets catalyze a conserved step in glycolysis [59, 60, 61].Specifically, they mediate the oxidative phosphorylation of glyceraldehyde 3-phosphate (G3P) to 1, 3-bisphosphoglycerate (BPG), using *NAD*^+^ or *NAD*(*P*)^+^ as a cofactor. This reaction proceeds through the formation of a hemiacetal intermediate with a catalytic cysteine, which is oxidized to a thioester, accompanied by the reduction of *NAD*^+^ to NADH [59, 60, 61]. After NADH is exchanged for a new *NAD*^+^ molecule, the thioester is attacked by inorganic phosphate to produce BPG.

##### Moonlighting/secondary function(s)

Here for each proximal set, moonlighting/secondary functions were outlined here.

###### C3MQN0

C3MQN0 exhibits a range of moonlighting activities. It has been shown to bind nucleic acids, thereby influencing both transcriptional regulation and mRNA stability. There is also growing evidence of its participation in DNA repair processes, which is particularly vital for the survival of *Sulfolobus islandicus* in its extreme environmental niche [62]. Additionally, this protein may be involved in the regulation of gene expression, membrane trafficking, and apoptosis-like responses under cellular stress. These multifunctional roles are believed to be modulated by its subcellular localization and post-translational modifications, highlighting the dynamic regulatory potential of GAPDH in archaea [62].

###### C3MWN0

While direct experimental characterization of the GAPDH protein’s (C3MWN0) secondary roles is currently lacking, it was found from the close sequence similarity to C3MQN0 that it shares similar moonlighting functions. These may include interactions with nucleic acids that affect transcription and RNA stability, as well as involvement in DNA repair pathways and cellular stress responses [62]. Its multifunctional character likely enables it to contribute to processes such as membrane organization, trafficking, and regulatory signaling, all of which are essential for adaptation to high-temperature, acidic environments. The extent of these roles may depend on post-translational modifications and dynamic changes in localization within the cell [62].

###### C3N6F0

C3N6F0 is part of this conserved genomic framework, implying evolutionary pressure to retain its function due to its central role in energy metabolism. The stability of such core components suggests essentiality for survival in the thermoacidophilic environments of *Sulfolobus islandicus* [62].

###### Q6M0E6

Specific experimental studies in *Methanococcus maripaludis* are lacking, Q6M0E6 is likely to moonlight in several secondary cellular roles. These may include nucleic acid binding, affecting mRNA stability or transcription regulation, participation in redox sensing and oxidative stress responses, and potential involvement in regulatory protein–protein interactions that are independent of its glycolytic activity [63].

###### A6VIW4

As with other GAPDH homologs, A6VIW4 is presumed to have additional moonlighting functions. These may include interactions with nucleic acids, enabling it to participate in the regulation of gene expression and RNA processing. It may also play a role in stress adaptation and serve as part of intracellular signaling pathways, particularly in response to environmental or redox stress [63].

###### A9A6M9

It may serve as an isoform with specialized or redundant roles in metabolism, potentially offering flexibility under different environmental or physiological conditions [63]. Like other GAPDH proteins, A9A6M9 is expected to have moonlighting capabilities. These may include nucleic acid binding, involvement in cellular stress response networks, and interaction with other macromolecules in roles beyond metabolism, although direct experimental evidence in *Methanococcus maripaludis* is currently lacking.

###### P0A9B3

Beyond its metabolic role, the GAPDH protein P0A9B3 in *Escherichia coli* was known to perform several non-glycolytic functions. It was reported to bind to DNA and RNA, participating in the regulation of gene expression and mRNA stability. GAPDH also associates with the bacterial membrane, where it may play roles in adhesion or hostpathogen interactions. Additionally, under oxidative stress or stationary-phase conditions, it participated in redox signaling and may act as a sensor of the cellular redox state. These secondary roles were often regulated by post-translational modifications such as oxidation of cysteine residues, which modulate the enzyme’s localization and functional state.

###### P0A1P0

Beyond its canonical role in glycolysis, the GAPDH protein P0A1P0 in *Salmonella typhimurium* was increasingly recognized for its moonlighting functions, particularly those relevant to bacterial pathogenesis and host interaction. Like its homologs in other prokaryotes, this GAPDH can localize to the cell surface, where it participated in adhesion to host tissues, enhancing colonization during infection. Additionally, P0A1P0 may play a regulatory role in gene expression by binding nucleic acids, and it may act as a redox sensor under oxidative stress conditions—common during host immune responses. These secondary functions reflect the enzyme’s ability to contribute to *Salmonella*’s virulence and stress adaptation, extending its role well beyond metabolism to include survival and persistence within the host.

###### *Proximal set-4* : Q4KYY3 (*Spermophilus citellus*), A3FKF7 (*Mustela putorius furo*)}

Beyond their canonical role in glycolysis, GAPDH proteins Q4KYY3 and A3FKF7 exhibit a wide range of moonlighting functions that extend into nuclear signaling, cytoskeletal organization, immune regulation, and post-translational modification. These proteins have S-nitrosylase activity, enabling them to mediate S-nitrosylation of nuclear targets such as SIRT1, HDAC2, and PRKDC, thereby participating in processes like transcriptional regulation, apoptosis, and DNA repair. They also play a role in microtubule dynamics by enhancing CHP1-dependent microtubule and membrane association, suggesting involvement in cytoskeletal remodeling. Additionally, both are components of the GAIT (gamma interferon-activated inhibitor of translation) complex, which selectively represses the translation of inflammatory mRNAs in response to interferon-*γ* signaling. Furthermore, they contribute to innate immunity by interacting with TRAF2 and TRAF3 to promote TNF-induced NF-*κ*B activation and Type-I interferon production. These diverse regulatory and structural roles highlight the multifunctionality of GAPDH beyond metabolism, integrating it into fundamental cellular processes across compartments [64, 65].

###### Canonical function

The GAPDH proteins encoded by P10096 in *Bos taurus* and Q9N2D5 in *Felis catus* catalyze a central step in glycolysis: the *NAD*^+^-dependent oxidative phosphorylation of D-glyceraldehyde 3-phosphate (G3P) to 3-phospho-D-glyceroyl phosphate [66, 67]. This reaction produced NADH and was found to be essential for cellular energy production and redox homeostasis. The catalytic mechanism involved the formation of a hemiacetal intermediate with a catalytic cysteine residue, its oxidation to a thioester, and subsequent attack by inorganic phosphate to generate the high-energy product used downstream in ATP synthesis [68, 69, 70].

Beyond their glycolytic role, GAPDH proteins P10096 (*Bos taurus*) and Q9N2D5 (*Felis catus*) possessed S-nitrosylase activity, enabling the S-nitrosylation of nuclear proteins such as SIRT1 and HDAC2, thus participating in transcriptional regulation and apoptosis [71]. Both GAPDH proteins contributed to cytoskeletal dynamics by facilitating CHP1-dependent microtubule interactions and are involved in immune modulation through the GAIT complex, suppressing translation of inflammatory mRNAs in response to interferon-*γ*. Additionally, they promote NF-*κ*B activation and Type-I interferon production via interactions with TRAF2 and TRAF3 [72].

###### *Proximal set-6* : {Q1DTF9 (*Coccidioides immitis*), Q8J1H3 (*Coccidioides posadasii*)}

GAPDH in fungi was well-recognized as a moonlighting adhesin that facilitates interactions with host components. For instance, in human-pathogenic fungi like *Candida albicans* and *Paracoccidioides* species, GAPDH was displayed on the cell surface, where it bound to extracellular matrix proteins such as fibronectin, laminin, and Type-I collagen, contributing to adhesion and invasion [73]. It also acted as a plasminogen receptor, enhancing extracellular matrix degradation to aid infection [74]. By analogy, Q1DTF9 and Q8J1H3 likely to serve similar non-glycolytic roles in host adhesion, tissue invasion, and possibly modulation of infection, thereby extending their functions beyond metabolism.

###### Q5RAB4

In addition to its metabolic role, Q5RAB4 possessed nitrosylase activity, mediating S-nitrosylation of critical nuclear proteins such as SIRT1, HDAC2, and PRKDC, thereby contributing to the regulation of transcription, DNA replication, RNA transport, and apoptosis [75]. It also played a role in cytoskeletal organization by promoting CHP1-dependent associations with microtubules and membranes [76]. Furthermore, Q5RAB4 was a component of the GAIT complex, which suppresses translation of inflammatory mRNAs upon interferon-gamma stimulation. It additionally contributeed to innate immune signaling by enhancing TNF-induced NF-*κ*B activation and Type-I interferon production through its interactions with TRAF2 and TRAF3 [75].

###### P04406

Beyond glycolysis, P04406 serves multiple moonlighting roles. It acted as a nitrosylase, mediating the Snitrosylation of nuclear proteins such as SIRT1, HDAC2, and PRKDC, influencing diverse nuclear processes including transcription, RNA transport, DNA replication, and apoptosis [77, 78]. P04406 also regulated cytoskeletal dynamics by supporting CHP1-mediated microtubule and membrane associations [79, 80]. It played an important role in immune regulation as a part of the GAIT complex, which bound to 3’-UTR GAIT elements in inflammatory mRNAs and suppresses their translation upon IFN-*γ* signaling [81, 82]. Moreover, it promoted the innate immunity by facilitating TNF-induced NF-*κ*B activation and Type-I interferon production through interactions with TRAF2 and TRAF3 [83, 84].

## 5. Discussion

This study presents a quantitative investigation of 165 GAPDH moonlighting proteins across Archaea, Bacteria, and Eukarya, revealing conserved canonical functions and diverse secondary roles shaped by compositional features, evolutionary constraints, and functional adaptations.

From the multiple sequence alignment and observed sequence variations among the 165 GAPDH moonlighting proteins, the presence of invariant residues strongly indicates conserved structural or catalytic roles essential to core functionality. Meanwhile, the presence of limited non-conservative substitutions and moderate overall variability suggests evolutionary constraints that permit divergence for functional innovation without compromising metabolic integrity.

The enrichment of valine and alanine across these sequences points to a structural basis for multifunctionality [85]. These hydrophobic residues likely contribute to core packing and interface flexibility—features crucial for enabling moonlighting behavior, especially under diverse or extreme environmental conditions [86]. Compositional analysis further confirmed the evolutionary selection of such residues, with valine and alanine exhibiting median frequencies of 10.21% and 9.61%, respectively. Though valine is typically a poor helix-former, Gregoret and Sauer demonstrated its improved stability within tertiary structures—much like alanine—highlighting the importance of structural context in protein flexibility [87]. In particular, extremophilic sequences such as P0A038, O83816, Q9Z518, Q8X1X3, and Q2FNA2 revealed elevated levels of small non-polar and charged residues, supporting adaptations to environmental stress. These findings align with prior studies emphasizing salt bridge networks and the spatial context of hydrophobic residues in protein architecture [88, 89].

This dual conservation and adaptability support the notion that GAPDH multifunctionality is encoded both at the sequence level and through spatial hydrophobic motifs [90]. Canonical examples, such as the use of GAPDH in glycolysis and extracellularly for plasminogen binding, further underscore the protein’s capacity for context-dependent functional switching [91]. The presence of solvent-exposed, non-conserved hydrophobic residues like valine and alanine may facilitate the evolution of novel binding interfaces, enabling functional diversification without the need for major structural reorganization.

Moreover, a distinct pattern emerged from the residue-level composition: GAPDH proteins were selectively enriched in both order-promoting residues (I, V, N, M) and disorder-promoting ones (T, S, G, A, K), while being depleted in classical order-promoting (C, W, Y, F, L, H) and disorder-promoting (R, Q, E) residues. This compositional duality likely reflects a balance between structural stability and the conformational flexibility required for multifunctionality [92].

Poly-string analysis further revealed tight compositional control [45]. Long homogeneous runs (length ≥ 4) were rare, suggesting an avoidance of excessive local homogeneity. Tri-residue repeats were most frequent for alanine and lysine, whereas residues like CCC, MMM, and WWW were completely absent, reinforcing selective compositional restraint. Di-residue repeats like TT and VV were widely present, indicating that short stretches of local uniformity are tolerated—perhaps to facilitate small-scale structural motifs that support dynamic functionality. Extended poly-string analysis showed the longest polar and non-polar runs to be of lengths 13 and 11, respectively—observed only in a few sequences. Most sequences lacked such extended homogeneity, again pointing toward structural restraint. However, selective enrichment of shorter polar and non-polar poly-strings indicates a functional compromise between local flexibility and global structural coherence.

Spatial analysis through binary indicator matrices, constructed for all 165 sequences, revealed low variance in fractal dimension and density of ones. This suggests a conserved underlying spatial complexity and distribution pattern of amino acids across sequences. In contrast, moderate variability in cluster counts using 4- and 8-connectivity hinted at subtle, but meaningful local differences in spatial organization that could influence folding or interactions unique to certain GAPDH homologs.

Functionally, all GAPDH proteins from the seven proximal sets catalyze the key glycolytic reaction: the *NAD*^+^ or *NAD*(*P*)^+^-dependent oxidation of D-glyceraldehyde-3-phosphate to 1,3-bisphosphoglycerate. This core function is deeply conserved across Archaea, Bacteria, and Eukarya, underscoring GAPDH’s role as a metabolic cornerstone. In archaeal proteins (e.g., *Sulfolobus islandicus* and *Methanococcus maripaludis*), this function supports autotrophic energy metabolism and redox balance under extreme conditions [91]. In bacteria such as *E. coli* and *S. typhimurium*, the reaction mechanism involves a cysteine-mediated thioester intermediate, contributing to robust ATP generation. Mammalian GAPDHs, including those from *Bos taurus, Felis catus*, and *Homo sapiens*, share this core glycolytic activity, but are also involved in redox signaling [91]. Similarly, fungal GAPDHs like those in *Coccidioides* species maintain classical catalytic roles necessary for pathogenic metabolism [73].

Beyond their canonical role, GAPDH proteins across the seven proximal sets demonstrate a broad array of moonlighting or secondary functions shaped by evolutionary and environmental pressures [93]. In Archaea, GAPDHs contribute to nucleic acid binding, transcription regulation, DNA repair, and cellular stress responses—functions critical to extremophile survival [94]. In bacteria, these proteins facilitate host adhesion, membrane localization, and act as redox sensors, contributing to both virulence and environmental resilience [95]. In mammalian systems, GAPDH moonlighting involves S-nitrosylase activity regulating transcription and apoptosis, microtubule organization via CHP1, and immune regulation via the GAIT complex and TRAF-mediated NF-*κ*B signaling. In fungi, GAPDHs act as surface adhesins and plasminogen receptors, aiding host invasion and infection [74]. These multifunctional roles are often modulated by subcellular localization and post-translational modifications, reflecting the protein’s structural plasticity and regulatory versatility [96, 97].

This comprehensive study reveals that GAPDH moonlighting functions are underpinned by subtle yet significant variations in sequence composition, structural motifs, and spatial organization. The conservation of canonical glycolytic roles across all domains of life affirms GAPDH’s indispensable metabolic function. Simultaneously, the evolution of secondary functions—enabled through compositional balance, structural plasticity, and localization dynamics—highlights the adaptability of this protein as a model for moonlighting behavior. Future studies could explore how these multifunctional roles are regulated in a domain-specific manner, particularly under stress or disease conditions. Structural modeling and mutagenesis studies could further elucidate how non-conserved residues contribute to binding promiscuity and regulatory interactions. Comparative studies across other moonlighting protein families would also help determine whether the compositional and spatial principles observed here are generalizable.

## Acknowledgments

The authors sincerely thank Mr. Ashim Dhar for his logistical support in the laboratory. A special thanks is extended to Mr. Arindam Samanta for his valuable and insightful comments. The authors also express their gratitude to the Indian Statistical Institute (ISI/TAC/PROJECT-1/2024-25) for their financial support. Notably, feature extractions and subsequent computations were carried out in MATLAB (2025a).

## Author contributions statement

SSH and DN formulated the problem and designed the theoretical experiments. DN, SSH, and NM carried out the experiments and performed the analyses. DN, SSH, NM, VNU, and MS drafted the initial manuscript. All authors contributed to reviewing and editing the manuscript. SSH and AG supervised the overall project. All authors reviewed, checked, and approved the final manuscript.

## References

[1] J. A. Shapiro, Revisiting the central dogma in the 21st century, Annals of the New York Academy of Sciences 1178 (1) (2009) 6–28.

[2] C. L. Tan, E. Anderson, The new central dogma of molecular biology, Resonance 14 (3) (2020) 1–32.

[3] J. Ule, B. J. Blencowe, Alternative splicing regulatory networks: functions, mechanisms, and evolution, Molecular cell 76 (2) (2019) 329–345.

[4] C. J. Jeffery, Protein moonlighting: what is it, and why is it important?, Philosophical transactions of the Royal Society B: biological sciences 373 (1738) (2018) 20160523.

[5] C. J. Jeffery, An introduction to protein moonlighting, Biochemical Society Transactions 42 (6) (2014) 1679–1683.

[6] N. Singh, N. Bhalla, Moonlighting proteins, Annual review of genetics 54 (1) (2020) 265–285.

[7] D. H. Huberts, I. J. van der Klei, Moonlighting proteins: an intriguing mode of multitasking, Biochimica et Biophysica Acta (BBA)-Molecular Cell Research 1803 (4) (2010) 520–525.

[8] A. Espinosa-Cantú, E. Cruz-Bonilla, L. Noda-Garcia, A. DeLuna, Multiple forms of multifunctional proteins in health and disease, Frontiers in Cell and Developmental Biology 8 (2020) 451.

[9] A. Gizak, Multitasking proteins and their involvement in pathogenesis (2023).

[10] P. Werelusz, S. Galiniak, M. Mołoń, Molecular functions of moonlighting proteins in cell metabolic processes, Biochimica et Biophysica Acta (BBA)-Molecular Cell Research 1871 (1) (2024) 119598.

[11] A. Kosova, S. Khodyreva, O. Lavrik, Role of glyceraldehyde-3-phosphate dehydrogenase (gapdh) in dna repair, Biochemistry (Moscow) 82 (2017) 643–654.

[12] N. El Kadmiri, I. Slassi, B. El Moutawakil, S. Nadifi, A. Tadevosyan, A. Hachem, A. Soukri, Glyceraldehyde-3-phosphate dehydrogenase (gapdh) and alzheimer’s disease, Pathologie Biologie 62 (6) (2014) 333–336.

[13] D.-M. Chuang, C. Hough, V. V. Senatorov, Glyceraldehyde-3-phosphate dehydrogenase, apoptosis, and neurodegenerative diseases, Annu. Rev. Pharmacol. Toxicol. 45 (1) (2005) 269–290.

[14] N. W. Seidler, N. W. Seidler, Compartmentation of gapdh, GAPDH: Biological Properties and Diversity (2013) 61–101.

[15] N. W. Seidler, Gapdh: biological properties and diversity (2012).

[16] D. Xu, F. Shao, X. Bian, Y. Meng, T. Liang, Z. Lu, The evolving landscape of noncanonical functions of metabolic enzymes in cancer and other pathologies, Cell metabolism 33 (1) (2021) 33–50.

[17] V. N. Uversky, Looking at the recent advances in understanding α-synuclein and its aggregation through the proteoform prism, F1000Research 6 (2017) 525.

[18] M. A. Sirover, Gapdh: a multifunctional moonlighting protein in eukaryotes and prokaryotes, Moonlighting Proteins: Novel Virulence Factors in Bacterial Infections (2017) 147–167.

[19] M. A. Sirover, Moonlighting glyceraldehyde-3-phosphate dehydrogenase: Posttranslational modification, protein and nucleic acid interactions in normal cells and in human pathology, Critical Reviews in Biochemistry and Molecular Biology 55 (4) (2020) 354–371.

[20] H. M. Ng, L. X. Kin, S. G. Dashper, N. Slakeski, C. A. Butler, E. C. Reynolds, Bacterial interactions in pathogenic subgingival plaque, Microbial Pathogenesis 94 (2016) 60–69.

[21] T. J. Foster, Colonization and infection of the human host by staphylococci: adhesion, survival and immune evasion, Veterinary dermatology 20 (5-6) (2009) 456–470.

[22] A. Thakur, H. Mikkelsen, G. Jungersen, Intracellular pathogens: host immunity and microbial persistence strategies, Journal of immunology research 2019 (1) (2019) 1356540.

[23] P.-C. Chen, M.-H. Hsieh, W.-S. Kuo, L. S.-H. Wu, H.-F. Kao, L.-F. Liu, Z.-G. Liu, W.-Y. Jeng, J.-Y. Wang, Moonlighting glyceraldehyde-3-phosphate dehydrogenase (gapdh) protein of lactobacillus gasseri attenuates allergic asthma via immunometabolic change in macrophages, Journal of biomedical science 29 (1) (2022) 75.

[24] M.-H. Hsieh, R.-L. Jan, L. S.-H. Wu, P.-C. Chen, H.-F. Kao, W.-S. Kuo, J.-Y. Wang, Lactobacillus gasseri attenuates allergic airway inflammation through pparγ activation in dendritic cells, Journal of Molecular Medicine 96 (2018) 39–51.

[25] C. Ye, L. Zhang, L. Tang, Y. Duan, J. Liu, H. Zhou, Host genetic backgrounds: the key to determining parasite-host adaptation, Frontiers in Cellular and Infection Microbiology 13 (2023) 1228206.

[26] A. El-Ansary, Biochemical and immunological adaptation in schistosome parasitism, Comparative Biochemistry and Physiology Part B: Biochemistry and Molecular Biology 136 (2) (2003) 227–243.

[27] W. F. Martin, R. Cerff, Physiology, phylogeny, early evolution, and gapdh, Protoplasma 254 (2017) 1823–1834.

[28] P. V. Shegay, O. P. Shatova, A. A. Zabolotneva, A. V. Shestopalov, A. D. Kaprin, Moonlight functions of glycolytic enzymes in cancer, Frontiers in Molecular Biosciences 10 (2023) 1076138.

[29] M. N. Gupta, V. N. Uversky, Moonlighting enzymes: when cellular context defines specificity, Cellular and Molecular Life Sciences 80 (5) (2023) 130.

[30] D. Nawn, S. S. Hassan, M. Sil, A. Ghosh, A. Goswami, P. Basu, G. W. Dayhoff II, K. Lundstrom, V. N. Uversky, The distal-proximal relationships among the human moonlighting proteins: Evolutionary hotspots and darwinian checkpoints, International Journal of Biological Macromolecules 259 (2024) 128998.

[31] C. Sulmon, J. Van Baaren, F. Cabello-Hurtado, G. Gouesbet, F. Hennion, C. Mony, D. Renault, M. Bormans, A. El Amrani, C. Wiegand, et al., Abiotic stressors and stress responses: What commonalities appear between species across biological organization levels?, Environmental Pollution 202 (2015) 66–77.

[32] B. I. Tieleman, Understanding immune function as a pace of life trait requires environmental context, Behavioral Ecology and Sociobiology 72 (3) (2018) 55.

[33] D. Nawn, S. S. Hassan, M. Sil, A. Ghosh, A. Goswami, V. N. Uversky, Proximal relationships of moonlighting proteins in escherichia coli: a mathematical genomic perspective, International Journal of Biological Macromolecules 308 (2025) 142766.

[34] F. Madeira, N. Madhusoodanan, J. Lee, A. Eusebi, A. Niewielska, A. R. Tivey, R. Lopez, S. Butcher, The embl-ebi job dispatcher sequence analysis tools framework in 2024, Nucleic acids research 52 (W1) (2024) W521–W525.

[35] M. Garcia-Boronat, C. M. Diez-Rivero, E. L. Reinherz, P. A. Reche, Pvs: a web server for protein sequence variability analysis tuned to facilitate conserved epitope discovery, Nucleic acids research 36 (suppl_2) (2008) W35–W41.

[36] C. M. Díez-Rivero, P. Reche, Discovery of conserved epitopes through sequence variability analyses, in: Bioinformatics for immunomics, Springer, 2009, pp. 95–101.

[37] M. H. Smith, The amino acid composition of proteins, Journal of Theoretical Biology 13 (1966) 261–282.

[38] S. S. Hassan, M. Sil, S. Chakraborty, A. Goswami, P. Basu, D. Nawn, V. N. Uversky, Possible functional proximity of various organisms based on the bioinformatics analysis of their taste receptors, International Journal of Biological Macromolecules 222 (2022) 2105–2121.

[39] S. S. Hassan, D. Nawn, A. Ghosh, M. Sil, A. Goswami, P. Basu, K. Lundstrom, V. N. Uversky, Methuselah proteins in drosophila: Structural and evolutionary insights from mathematical genomics, Biochemical and Biophysical Research Communications (2025) 152240.

[40] V. Vacic, V. N. Uversky, A. K. Dunker, S. Lonardi, Composition profiler: a tool for discovery and visualization of amino acid composition differences, BMC bioinformatics 8 (2007) 1–7.

[41] M. Sickmeier, J. A. Hamilton, T. LeGall, V. Vacic, M. S. Cortese, A. Tantos, B. Szabo, P. Tompa, J. Chen, V. N. Uversky, et al., Disprot: the database of disordered proteins, Nucleic acids research 35 (suppl_1) (2007) D786–D793.

[42] D. Piovesan, F. Tabaro, I. Mičetić, M. Necci, F. Quaglia, C. J. Oldfield, M. C. Aspromonte, N. E. Davey, R. Davidović, Z. Dosztányi, et al., Disprot 7.0: a major update of the database of disordered proteins, Nucleic acids research 45 (D1) (2017) D219–D227.

[43] M. C. Aspromonte, M. V. Nugnes, F. Quaglia, A. Bouharoua, S. C. Tosatto, D. Piovesan, Disprot in 2024: improving function annotation of intrinsically disordered proteins, Nucleic Acids Research 52 (D1) (2024) D434–D441.

[44] D. Nawn, S. S. Hassan, E. M. Redwan, T. Bhattacharya, P. Basu, K. Lundstrom, V. N. Uversky, Unveiling the genetic tapestry: Rare disease genomics of spinal muscular atrophy and phenylketonuria proteins, International Journal of Biological Macromolecules 269 (2024) 131960.

[45] D. Nawn, S. S. Hassan, A. Hromić-Jahjefendić, T. Bhattacharya, P. Basu, E. M. Redwan, D. Barh, B. S. Andrade, A. A. Aljabali, Á. Serrano-Aroca, et al., Molecular genomic insights into melanoma associated proteins prame and bap1, Journal of Biomolecular Structure and Dynamics (2025) 1–31.

[46] M. Y. Karpeisky, V. Ilyin, Analysis of non-polar regions in proteins, Journal of molecular biology 224 (3) (1992) 629–638.

[47] Z. M. Frenkel, E. N. Trifonov, Walking through protein sequence space, Journal of theoretical biology 244 (1) (2007) 77–80.

[48] G. W. Dayhoff, V. N. Uversky, Rapid prediction and analysis of protein intrinsic disorder, Protein Science 31 (12) (2022) e4496.

[49] W.-L. Hsu, C. Oldfield, J. Meng, F. Huang, B. Xue, V. N. Uversky, P. Romero, A. K. Dunker, Intrinsic protein disorder and protein-protein interactions, in: Biocomputing 2012, World Scientific, 2012, pp. 116–127.

[50] G. W. Dayhoff, M. H. van Regenmortel, V. N. Uversky, Intrinsic disorder in protein sense-antisense recognition, Journal of Molecular Recognition 33 (10) (2020) e2868.

[51] J. Habchi, P. Tompa, S. Longhi, V. N. Uversky, Introducing protein intrinsic disorder, Chemical reviews 114 (13) (2014) 6561–6588.

[52] C. Cattani, Fractals and hidden symmetries in dna, Mathematical problems in engineering 2010 (1) (2010) 507056.

[53] R. J. Boys, D. A. Henderson, D. J. Wilkinson, Detecting homogeneous segments in dna sequences by using hidden markov models, Journal of the Royal Statistical Society: Series C (Applied Statistics) 49 (2) (2000) 269–285.

[54] A. K. Dunker, J. D. Lawson, C. J. Brown, R. M. Williams, P. Romero, J. S. Oh, C. J. Oldfield, A. M. Campen, C. M. Ratliff, K. W. Hipps, et al., Intrinsically disordered protein, Journal of molecular graphics and modelling 19 (1) (2001) 26–59.

[55] P. Romero, Z. Obradovic, X. Li, E. C. Garner, C. J. Brown, A. K. Dunker, Sequence complexity of disordered protein, Proteins: Structure, Function, and Bioinformatics 42 (1) (2001) 38–48.

[56] R. Williams, Z. Obradovic, V. Mathura, W. Braun, E. Garner, J. Young, S. Takayama, C. J. Brown, A. K. Dunker, The protein non-folding problem: amino acid determinants of intrinsic order and disorder, in: Biocomputing 2001, World Scientific, 2000, pp. 89–100.

[57] P. Radivojac, L. M. Iakoucheva, C. J. Oldfield, Z. Obradovic, V. N. Uversky, A. K. Dunker, Intrinsic disorder and functional proteomics, Biophysical journal 92 (5) (2007) 1439–1456.

[58] A. Campen, R. M. Williams, C. J. Brown, J. Meng, V. N. Uversky, A. K. Dunker, Top-idp-scale: a new amino acid scale measuring propensity for intrinsic disorder, Protein and peptide letters 15 (9) (2008) 956–963.

[59] R. A. Welch, V. Burland, G. Plunkett III, P. Redford, P. Roesch, D. Rasko, E. L. Buckles, S.-R. Liou, A. Boutin, J. Hackett, et al., Extensive mosaic structure revealed by the complete genome sequence of uropathogenic escherichia coli, Proceedings of the National Academy of Sciences 99 (26) (2002) 17020–17024.

[60] M. McClelland, K. E. Sanderson, J. Spieth, S. W. Clifton, P. Latreille, L. Courtney, S. Porwollik, J. Ali, M. Dante, F. Du, et al., Complete genome sequence of salmonella enterica serovar typhimurium lt2, Nature 413 (6858) (2001) 852–856.

[61] J. G. Lawrence, H. Ochman, D. L. Hartl, Molecular and evolutionary relationships among enteric bacteria, Microbiology 137 (8) (1991) 1911–1921.

[62] M. L. Reno, N. L. Held, C. J. Fields, P. V. Burke, R. J. Whitaker, Biogeography of the sulfolobus islandicus pangenome, Proceedings of the National Academy of Sciences 106 (21) (2009) 8605–8610.

[63] E. Hendrickson, R. Kaul, Y. Zhou, D. Bovee, P. Chapman, J. Chung, E. Conway de Macario, J. Dodsworth, W. Gillett, D. Graham, et al., Complete genome sequence of the genetically tractable hydrogenotrophic methanogen methanococcus maripaludis, Journal of bacteriology 186 (20) (2004) 6956–6969.

[64] I. Naletova, E. Schmalhausen, B. Tomasello, D. Pozdyshev, F. Attanasio, V. Muronetz, The role of sperm-specific glyceraldehyde-3-phosphate dehydrogenase in the development of pathologies—from asthenozoospermia to carcinogenesis, Frontiers in Molecular Biosciences 10 (2023) 1256963.

[65] N. Svitek, V. von Messling, Early cytokine mrna expression profiles predict morbillivirus disease outcome in ferrets, Virology 362 (2) (2007) 404–410.

[66] K. D. Kulbe, K. W. Jackson, J. Tang, Structural evidence for a liver-specific glyceraldehyde-3-phosphate dehydrogenase, Biochemical and Biophysical Research Communications 67 (1) (1975) 35–42.

[67] J. Oláh, N. Tőkési, O. Vincze, I. Horváth, A. Lehotzky, A. Erdei, E. Szájli, K. F. Medzihradszky, F. Orosz, G. G. Kovács, et al., Interaction of tppp/p25 protein with glyceraldehyde-3-phosphate dehydrogenase and their co-localization in lewy bodies, FEBS letters 580 (25) (2006) 5807–5814.

[68] A. K. Helfer-Hungerbuehler, S. Widmer, R. Hofmann-Lehmann, Gapdh pseudogenes and the quantification of feline genomic dna equivalents, Molecular biology international 2013 (1) (2013) 587680.

[69] C. E. Morgan, Z. Zhang, M. Miyagi, M. Golczak, E. W. Yu, Toward structural-omics of the bovine retinal pigment epithelium, Cell Reports 41 (13) (2022).

[70] B. Y. Baker, W. Shi, B. Wang, K. Palczewski, High-resolution crystal structures of the photoreceptor glyceraldehyde 3-phosphate dehydrogenase (gapdh) with three and four-bound nad molecules, Protein Science 23 (11) (2014) 1629–1639.

[71] G. Russell, D. Veal, J. T. Hancock, Is glyceraldehyde-3-phosphate dehydrogenase a central redox mediator, React. Oxyg. Species 9 (2020) 48–69.

[72] C. Tristan, N. Shahani, T. W. Sedlak, A. Sawa, The diverse functions of gapdh: views from different subcellular compartments, Cellular signalling 23 (2) (2011) 317–323.

[73] C. M. Marcos, H. C. d. Oliveira, J. d. F. da Silva, P. A. Assato, A. M. Fusco-Almeida, M. J. Mendes-Giannini, The multifaceted roles of metabolic enzymes in the paracoccidioides species complex, Frontiers in Microbiology 5 (2014) 719.

[74] J. Wang, Y. Li, L. Pan, J. Li, Y. Yu, B. Liu, M. Zubair, Y. Wei, B. Pillay, A. O. Olaniran, et al., Glyceraldehyde-3-phosphate dehydrogenase (gapdh) moonlights as an adhesin in mycoplasma hyorhinis adhesion to epithelial cells as well as a plasminogen receptor mediating extracellular matrix degradation, Veterinary Research 52 (1) (2021) 80.

[75] S. Bechtel, H. Rosenfelder, A. Duda, C. P. Schmidt, U. Ernst, R. Wellenreuther, A. Mehrle, C. Schuster, A. Bahr, H. Blöcker, et al., The full-orf clone resource of the german cdna consortium, BMC genomics 8 (2007) 1–12.

[76] A. Hanauer, J. L. Mandel, The glyceraldehyde 3 phosphate dehydrogenase gene family: structure of a human cdna and of an x chromosome linked pseudogene; amazing complexity of the gene family in mouse., The EMBO journal 3 (11) (1984) 2627–2633.

[77] K. Nowak, M. Wolny, T. Banaś, The complete amino acid sequence of human muscle glyceraldehyde 3-phosphate dehydrogenase, FEBS letters 134 (2) (1981) 143–146.

[78] J. Y. Tso, X.-H. Sun, T.-h. Kao, K. S. Reece, R. Wu, Isolation and characterization of rat and human glyceraldehyde-3-phosphate dehydrogenase cdnas: genomic complexity and molecular evolution of the gene, Nucleic acids research 13 (7) (1985) 2485–2502.

[79] K. Tokunaga, Y. Nakamura, K. Sakata, K. Fujimori, M. Ohkubo, K. Sawada, S. Sakiyama, Enhanced expression of a glyceraldehyde-3-phosphate dehydrogenase gene in human lung cancers, Cancer research 47 (21) (1987) 5616–5619.

[80] R. Allen, K. Trach, J. Hoch, Identification of the 37-kda protein displaying a variable interaction with the erythroid cell membrane as glyceraldehyde-3-phosphate dehydrogenase., Journal of Biological Chemistry 262 (2) (1987) 649–653.

[81] A. Arif, P. Chatterjee, R. A. Moodt, P. L. Fox, Heterotrimeric gait complex drives transcript-selective translation inhibition in murine macrophages, Molecular and Cellular Biology 32 (24) (2012) 5046–5055.

[82] L. Ercolani, B. Florence, M. Denaro, M. Alexander, Isolation and complete sequence of a functional human glyceraldehyde-3-phosphate dehydrogenase gene., Journal of Biological Chemistry 263 (30) (1988) 15335–15341.

[83] X. Gao, X. Wang, T. H. Pham, L. A. Feuerbacher, M.-L. Lubos, M. Huang, R. Olsen, A. Mushegian, C. Slawson, P. R. Hardwidge, Nleb, a bacterial effector with glycosyltransferase activity, targets gapdh function to inhibit nf-κb activation, Cell host & microbe 13 (1) (2013) 87–99.

[84] X. Gao, T. H. Pham, L. A. Feuerbacher, K. Chen, M. P. Hays, G. Singh, C. Rueter, R. Hurtado-Guerrero, P. R. Hardwidge, Citrobacter rodentium nleb protein inhibits tumor necrosis factor (tnf) receptor-associated factor 3 (traf3) ubiquitination to reduce host type i interferon production, Journal of Biological Chemistry 291 (35) (2016) 18232–18238.

[85] M. V. Katti, R. Sami-Subbu, P. K. Ranjekar, V. S. Gupta, Amino acid repeat patterns in protein sequences: their diversity and structural-functional implications, Protein Science 9 (6) (2000) 1203–1209.

[86] A. Ghaffarkhah, S. A. Hashemi, A. A. Isari, M. Panahi-Sarmad, F. Jiang, T. P. Russell, O. J. Rojas, M. Arjmand, Chemistry, applications, and future prospects of structured liquids, Chemical Society Reviews (2024).

[87] L. M. Gregoret, R. T. Sauer, Tolerance of a protein helix to multiple alanine and valine substitutions, Folding and Design 3 (2) (1998) 119–126.

[88] A. S. Panja, S. Maiti, B. Bandyopadhyay, Protein stability governed by its structural plasticity is inferred by physicochemical factors and salt bridges, Scientific reports 10 (1) (2020) 1822.

[89] M. H. Al Mughram, C. Catalano, N. B. Herrington, M. K. Safo, G. E. Kellogg, 3d interaction homology: The hydrophobic residues alanine, isoleucine, leucine, proline and valine play different structural roles in soluble and membrane proteins, Frontiers in Molecular Biosciences 10 (2023) 1116868.

[90] N. W. Seidler, N. W. Seidler, Functional diversity, GAPDH: Biological Properties and Diversity (2013) 103–147.

[91] S. Parvez, M. J. Long, J. R. Poganik, Y. Aye, Redox signaling by reactive electrophiles and oxidants, Chemical reviews 118 (18) (2018) 8798–8888.

[92] Y. Zhang, M. Barboiu, Constitutional dynamic materials toward natural selection of function, Chemical Reviews 116 (3) (2016) 809–834.

[93] M. A. Sirover, Glyceraldehyde-3-phosphate dehydrogenase (GAPDH): the quintessential moonlighting protein in normal cell function and in human disease, Academic Press, 2017.

[94] R. Cavicchioli, T. Thomas, P. M. Curmi, Cold stress response in archaea, Extremophiles 4 (2000) 321–331.

[95] M. L. Reniere, Reduce, induce, thrive: bacterial redox sensing during pathogenesis, Journal of bacteriology 200 (17) (2018) 10–1128.

[96] W. Li, F. Li, X. Zhang, H.-K. Lin, C. Xu, Insights into the post-translational modification and its emerging role in shaping the tumor microenvironment, Signal transduction and targeted therapy 6 (1) (2021) 422.

[97] S. Mittal, D. Saluja, Protein post-translational modifications: role in protein structure, function and stability, Proteostasis and chaperone surveillance (2015) 25–37.

